# Cell-free immuno-profiling on a genetically programmed biochip

**DOI:** 10.1101/2024.10.09.617356

**Authors:** Aurore Dupin, Ohad Vonshak, Valerie Arad, Maya Levanon, Noa Avidan, Yiftach Divon, Steve Peleg, Seth Thompson, Vincent Noireaux, Shirley S. Daube, Roy H. Bar-Ziv

## Abstract

Emerging cell-free synthetic biology approaches provide biosafe, cheap, and versatile genetic tools to advance therapeutic research and development. Combined with micro-fabrication technology, we developed a platform to quantitatively reconstitute interactions of cell-free synthesized antigens with antibodies and human receptors in miniaturized compartments on a silicon chip. Photolithographic surface patterning of protein traps and on chip expression from high density gene brushes generated a continuous surface density gradient of fluorescently labeled antigens. Antibodies binding to the antigen gradient generate a full binding curve in each single compartment for affinity determination. We used the SARS-CoV-2 antigens as a model to profile the specificity and affinity of monoclonal antibodies to > 30 viral epitopes synthesized simultaneously on one chip in a genotype-phenotype linked compartments. We further profiled polyclonal antibodies in minute volumes of human sera, revealing patient-specific epitope profiles that are difficult to detect by conventional approaches. Cell-free co-synthesis of the human ACE2 receptor with the viral Receptor-Binding-Domain yielded relative binding affinities to different SARS-CoV-2 variants. This rapid, quantitative, and on-chip genetically programmed approach allows to study complex protein-protein interactions independent of protein purification steps for human immuno-profiling with a fast response time for combating emerging pathogens.

## Main

Molecular characterization of protein-protein interactions is fundamental for diagnosing and monitoring clinical conditions and for developing drugs to target human disease. The ability to profile the immune response by characterizing antibody (Ab) abundance, epitope specificity and binding strength is valuable for various clinical therapeutic applications, from cancer immunotherapy to vaccine development^1–4^. Using biotherapeutic drugs such as monoclonal antibodies (mAbs) has emerged in recent years as a leading approach to target human disease, tailored to disrupt or enhance protein-protein interactions that are the underlying cause of a clinical condition^5–8^. A common approach in the development of biotherapeutics is to search through large sequence-space libraries for potential leads with high specificity and efficacy^9–12^. Yet, high-throughput approaches fall short in providing detailed and quantitative molecular information, which requires time consuming and expensive iterative protein purification steps. Therefore, quantitative characterization is currently limited to a handful of candidates.

Cell-free gene expression (CFE) is a bio-safe, cheap, fast and versatile approach to advance clinical and biopharmaceutical research^13–15^. Target proteins can be synthesized at high yields with no risk of toxicity^16–20^, gene circuits can be designed for point-of-care biosensing and diagnostics^21–23^, mAbs can be synthesized and screened for specific epitope binding^24–26^ and protein-arrays have been generated from their coding genes for Ab profiling ^27–30^. In the past decade we developed a biochip approach as a minimal model of an artificial cell. We demonstrated reconstitution of complex biochemical interactions, such as the biogenesis of the *E. coli* ribosomal subunit and T4 bacteriophage structural domains by cell-free co-synthesis, localization and confinement of nascent proteins^31,32^. We use dozens of miniaturized compartments carved on a silicon chip, with genes immobilized in each compartment at high density as DNA brushes. Upon addition of a bacterial CFE system and sealing, forming a sub-nanoliter volume reaction chamber, proteins synthesized in situ are captured by surface traps surrounding the DNA, creating a genotype-phenotype linkage.

Here we tailor the biochip platform to screen dozens of protein-protein interactions without compromising detailed and quantitative molecular characterization while requiring minute volumes of human samples. As a model system, we demonstrate CFE of antigens of the SARS-CoV-2 human pathogen, specifically the nucleoprotein (N), the S1 spike protein and the Receptor Binding Domain (RBD), all on one chip in soluble form, available to specific binding by mAbs and polyclonal Abs (pAbs). On-chip co-synthesis of the soluble domain of ACE2 human receptor with different RBD variants revealed their relative binding strength. Each compartment could be patterned by photolithography to generate an antigen concentration gradient, allowing to establish in each single compartment a full binding curve with the corresponding Abs and derive simultaneously dozens of affinity equilibrium constants from one chip. We profiled pAbs in human sera samples and commercial mAbs using a panel of full-length viral antigens and their variants, in addition to a systematic screen of N and S1 sub-fragments, establishing an epitope map for 3 previously uncharacterized mAbs, and providing a rich dataset of human sera that cannot be easily obtained with conventional Ab detection techniques.

### Multiplexed antigens display for on chip quantitative antibody-antigen binding assay

In a first chip layout, 384 (array of 12 by 16) circular compartments of 150 µm radius and 5 µm height were etched in a silicon chip (Fig. 1A, Supplementary Fig. 1). Each compartment was patterned in its centre with surface-bound linear DNA polymers forming a high-density gene brush coding for an antigen of interest under a T7 promoter and fused to the GFP gene and an HA peptide tag (Fig. 1A,B, Methods, Supplementary Table 1). Upon addition of an *E. coli* CFE system^33^ onto the chip, antigens fused to GFP-HA were synthesized simultaneously in each compartment according to its genetic program and captured on anti-HA Abs immobilized on the surface surrounding the DNA (Fig. 1C,D, Methods). Specifically, we synthesized in different compartments on one chip the SARS-CoV-2 Nucleocapsid protein (N-GFP-HA) in addition to two control antigens, His-tagged and FLAG-tagged GFP (His-GFP-HA, FLAG-GFP-HA, respectively). The gene fraction in different compartments was varied by mixing it with different amounts of a promoter-less DNA of the same length to maintain a constant DNA brush density, leading to a protein synthesis proportional to gene fraction^34^. Following CFE and washing, the chip was incubated with the anti-N mAb 1A6, which was later detected by a second incubation with fluorescently labeled secondary Abs (Fig. 1C, Methods). Two more identical chips were exposed after CFE to anti-His and anti-FLAG mAbs, respectively. We detected specific binding of each mAb to its target antigen, with low binding to its non-specific targets (Fig. 1A inset, Supplementary Fig. 2).

**Figure 1.**
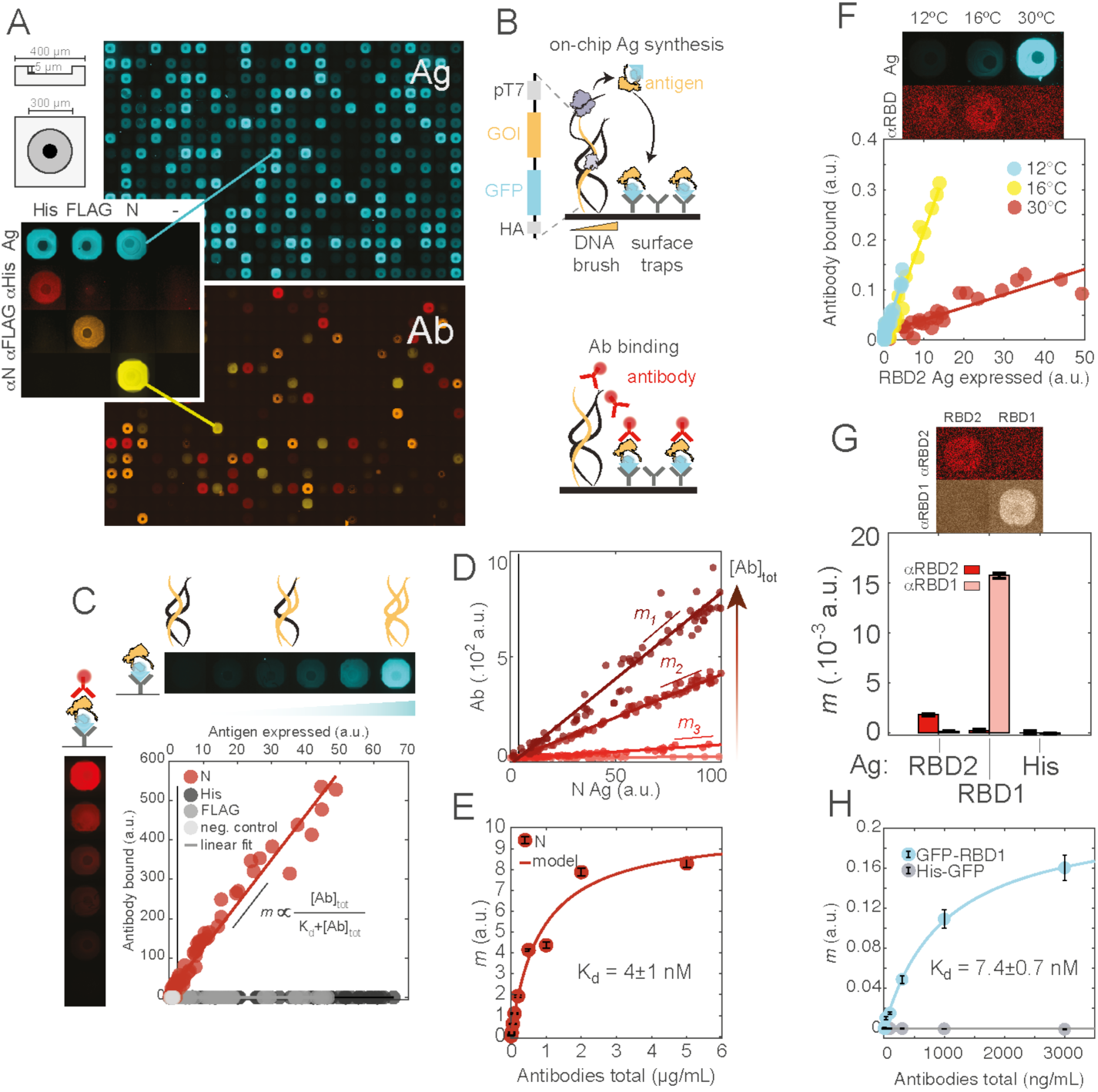
Multiplex quantitative characterization of antigen-antibody binding. A: Microcompartments array for in situ cell-free expression. Left: sketch of a single compartment with a DNA brush (dark circle) spotted in the center. Montage of fluorescence microscopy images of all compartments on one chip, showing captured antigens (top, GFP labeled, blue), and antibody staining (bottom, composite image, anti-His, red, anti-FLAG, orange, anti-N, yellow). Inset: single compartment microscopy images of antigen GFP fluorescence (blue) and monoclonal antibody binding: anti-His (αHis, red), anti-FLAG (αFLAG, orange), anti-N (αN, yellow). Scale of all microscopy images: one compartment image has a width of 400 μm. B: The DNA brush contains a linear synthetic gene coding for the gene of interest (GOI) tagged with a GFP and HA tag under a T7 promoter. Anti-HA antibody traps cover the rest of the surface. Expression from the DNA brush in an *E. coli* cell-free expression system leads to subsequent capture of the antigen on the surface. Antibody samples are characterized by incubation on the chip following washing. C: Quantitative measurement of antigen-antibody binding. Antibody fluorescence against antigen fluorescence for an anti-N antibody binding to N-expressing compartments (red), His (dark grey), FLAG (median grey) or a negative control protein unrelated to SARS-CoV and non-fluorescent but HA-tagged (NC-HA, light grey). Dots represent single data points and lines represent linear regression, with *m* the slope of antibody recognition. Left and top: representative microscopy images of N-expressing compartments (GFP fluorescence, blue) bound by anti-N antibodies (red). D,E: Titration of total antibody concentration and characterization of affinity. D: Linear dependency of bound antibody to bound antigen for 4 concentrations of total antibodies: 5 (light red), 50 (median light red), 500 (median dark red) and 5000 (dark red) ng/mL. Dots and lines are as in C. E: Red circles with black error bars represent fitted *m* value and 95% confidence interval of the fit. *m* values were fitted from 84 data points. Affinity is measured from fitting all antibody titration curves simultaneously. F: RBD2 Ag temperature of cell-free expression and Ab recognition. Fluorescence images of compartments expressing GFP-RBD2 (GFP signal, blue, top row) recognized by the anti-RBD2 mAb CV30 antibody (antibody signal, red, bottom row) at different temperatures (12 °C, left, 16 °C, middle, 30 °C, right). Antigen-antibody binding function for at least 23 compartments at 3 different temperatures (12 °C, blue, 16 °C, yellow, 30 °C, red). Dots represent single data points and lines represent linear regression. G: Orthogonal binding of two specific antibodies to two closely related antigens. Microscopy images of antibody binding to SARS-CoV-2 RBD (RBD2) or SARS-CoV-1 RBD (RBD1), red: anti-RBD2, pink: anti-RBD1. Fluorescence is measured from a labeled secondary antibody. Antigen-antibody binding slope (red: anti-RBD2, pink: anti-RBD1) to RBD2 and RBD1. Bars and error bars represent fitted *m* value and 95% confidence interval of the fit. H: Titration of anti-RBD1 mAb CR3022. 20 compartments expressing each given antigen specie are fitted with a linear fit for each antibody concentration. Circles and error bars represent the fitted slope value and a 95% confidence interval of the fit. Affinity is measured from fitting all antibody titration curves simultaneously.

We evaluated N CFE levels by the GFP signal and the corresponding mAb binding level by the secondary Ab signal, resulting in a linear response curve (Fig. 1C). The linear dependency between N antigens and bound Abs for the specific pair agreed well with a limit case of the chemical equilibrium *Ag* + *Ab* ⇌ *Ag* · *Ab* where Abs are in excess compared to antigens (Supplementary Information, Supplementary Fig. 3). In this limit case, the slope *m* of the linear antigen-Ab binding is proportional to the total Ab concentration and the binding affinity with 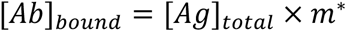 and 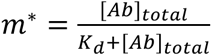. Therefore, by simply diluting genes on-chip we could obtain a titration curve of antigen-Ab interaction.

Based on our model, we hypothesized that antigen-antibody affinity *K*_*d*_ could be derived by obtaining several on-chip antigen-antibody response curves at different total mAb concentration. We patterned twelve identical chips, each with compartments with variable concentration of N genes and a negative control. Following antigen CFE and capture, we incubated each chip with a different concentration of the same anti-N 1A6, ranging over three orders of magnitude (Fig. 1d, Supplementary Fig. 4). We found that the slope *m* increased with increasing Ab concentration (Fig. 1E), consistent with the above model. We determined an on-chip *K*_*d*_ of 558±197 ng/mL (∼4 ±1 nM, averaged from three titrations). This affinity was confirmed with biolayer interferometry (BLI) using cell-free synthesized antigen (*K*_*d*,*BLI*_ = 8±4 nM, Methods, Supplementary Fig. 5). Other studies reported an EC_50_ of 43.50 - 118.4 ng/mL (∼0.3-0.7 nM) with ELISA^35^, and an affinity *K*_*d*_ < 0.7 nM with BLI^36^. As reported, these methods required 50-100 nM purified N protein whereas no purified protein was required to generate our data. mAb binding to N was still distinguishable from non-specific binding to a control antigen at mAb concentration as low as 0.1 ng/mL (Methods, Supplementary Fig. 4). This chip layout required a mAb volume of ∼ 5 μL per compartment compared to ELISA kits with a similar detection limit ^37^ requiring typically 100 μL of Ab sample per well.

### Temperature modulation of antigen synthesis and antibody binding

To demonstrate the generality of the on-chip immuno-profiling platform we tested CFE and Ab detection of the SARS-CoV-2 S1 and its RBD (RBD2). Initial CFE reactions off-chip indicated that the S1 and RBD antigens with a C terminus (C-ter) GFP-HA fusion expressed about one order of magnitude lower than the N-GFP-HA antigen (Supplementary Fig. 6). An N terminus (N-ter) GFP fusion improved the expression yet was still lower than N-GFP-HA expression, and expression of both S1-GFP-HA and GFP-S1-HA in a human cell lysate were even lower, suggesting that the *E. coli* lysate was not the underlying reason for low expression of the mammalian antigens (Methods, Supplementary Fig. 6). Inspired by a common practice in recombinant protein synthesis, we intuited that low synthesis was due to misfolding, incomplete translation or aggregation and that protein detection could be improved by CFE at lower temperatures, at the cost of a slower protein synthesis rates. We found indeed that for all antigen constructs, CFE at temperatures lower than the usual 30°C resulted in increased CFE signals (Supplementary Fig. 6).

Interestingly, on-chip expression of GFP-RBD2-HA at various temperatures revealed a significant increase in binding of the anti-RBD2 mAb CV30 as the temperature decreased, despite lower CFE (Fig. 1F). A similar effect was observed for the N antigen and its specific mAb, and additionally we found that a shorter expression time led to an improved mAb recognition (Supplementary Fig. 7). Taken together, these data suggest that the GFP-fused antigens are better recognized by Abs when synthesized at lower temperatures, most likely due to their improved correct folding and solubility. The slower translation rate may improve co-translational folding and limit the collapse of insoluble proteins into misfolded and aggregating forms. We subsequently chose intermediary on-chip CFE temperatures (16-18°C) to allow for sufficient protein synthesis for detection while minimizing protein misfolding.

With these improved CFE conditions of the RBD antigens on chip we could compare the binding specificity of the anti-RBD2 mAb CV30 to that of CR3022, a mAb recognizing the RBD of the SARS-CoV emerged in 2002-2003 (RBD1). Indeed, we found that each of these two human patients derived mAbs had high selectivity towards their corresponding antigens (RBD1 and RBD2 to CR3022 and CV30, respectively) (Fig. 1G). Similarly to the titration of anti-N mAb (Fig. 1d,e), we performed an on-chip titration of the CR3022 mAb to the RBD1 antigen synthesis on chip and calculated a *K*_*d*_=7.4±0.7 nM (Fig. 1H). Furthermore, we tested the effect of disulfide bond formation on mAb recognition by comparing synthesis at reducing or oxidizing conditions (Methods, Supplementary Fig. 8). mAb binding at the oxidizing conditions, that promote disulfide bonds, was only slightly increased compared to the reducing conditions, suggesting that mAb recognition was not hindered by incorrect formation of disulfide bonds.

### A full antigen-antibody binding curve in one compartment

The quantitative antigen-antibody interaction analysis presented in Fig. 1 used over >10 compartments of variable gene fraction per antigen. To dramatically increase the throughput of each chip and the accuracy of the measurement, we used a different chip layout with elongated compartments of 750 µm x 200 µm and 10 µm height (Fig. 2B), previously shown to provide increased spatial resolution of protein synthesis and capture^32^. We created a spatial gradient of surface traps along the long axis of the compartment by UV lithography (Methods). A DNA brush coding fully for the desired antigen was immobilized on one end of the compartment, bypassing the need to prepare multiple gene fraction solutions. After CFE from the immobilized DNA brush, a linear and reproducible gradient of antigens was displayed on the surface (Fig. 2A,B). Addition of mAbs onto the chip exposed them to a broad range of antigen surface densities, and by averaging over the short axis and considering a pixel size of 1-2 µm, we were able to resolve >300 data points in a single compartment, drastically improving the resolution of the antigen titration. This approach can be considered similarly to a Langmuir adsorption isotherm, where a single compartment displays an array of densities of adsorption sites. The adsorbent (the Ab) adsorbs onto immobile sites of antigens arranged as a monolayer, with no interactions between adjacent sites. A single chip presents 96 different antigen-antibody binding isotherms with improved reproducibility as well as resolution, since the antigen gradient is dominated by the surface patterning and not just by the CFE level. With this approach, we recapitulated the linear antigen-antibody binding function measured in the circular compartments and the Ab-concentration dependent increase of the slope *m*, as well as the affinity measurement (Fig. 2C,D, Supplementary Fig. 9). Performing a total Ab titration, we were able to measure the affinity of anti-RBD2 mAb 5G8 to RBD2 (*K*_*d*_=10.9±0.2 nM, confirmed with BLI, Supplementary Fig. 5).

**Figure 2.**
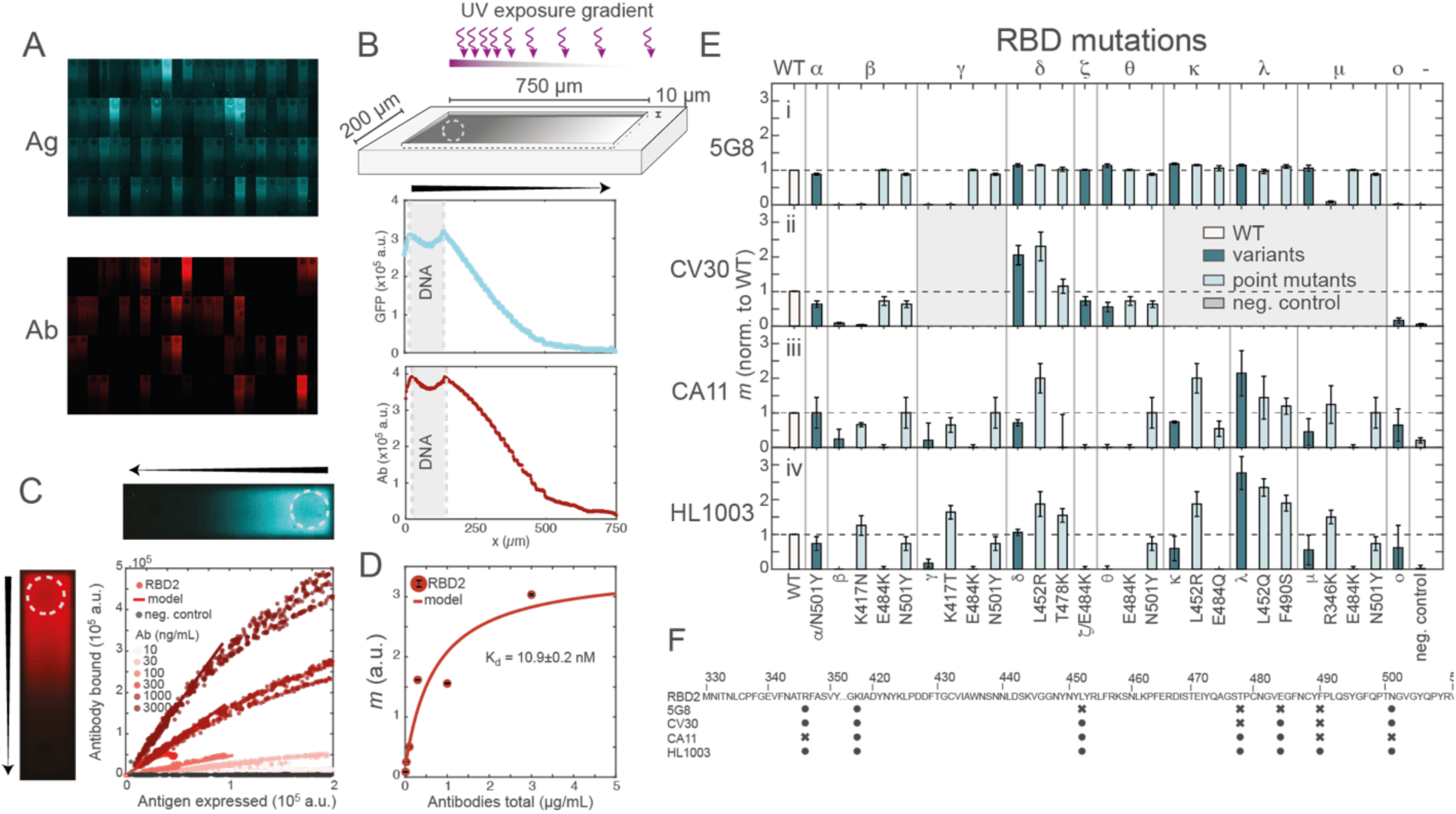
One compartment antigen-antibody titration, variants characterization. A: Fluorescence montage of all compartments on a chip (96 compartments) Ag synthesis (top, blue), and anti-RBD2 mAb 5G8 binding (bottom, red). Scale for all microscopy images: the long axis of one compartment is 750 µm. B: The surface is activated by gradient UV exposure, which leads to a linear gradient of surface activation and of surface traps (top). Following protein synthesis, the antigen is displayed as a gradient (blue, center). The consequent binding of the antibody to the antigen (red, bottom) allows for the titration of the antigen-antibody interaction in a single compartment. DNA brush is indicated with dashed line. Arrow indicates direction of the gradient from high to low density. C,D: Antibody titration and affinity measurement in elongated compartments. C: bound antibody as a function of displayed antigen for four compartments displaying GFP-RBD2 WT-HA (shades of red) or a negative control His-GFP-HA (shades of grey) for different 5G8 mAb incubation concentration. Each compartment consists of 320 individual data points along the long axis of the compartment. DNA brush is indicated with dashed line. Arrow indicates direction of the gradient from high to low density.D: Red dots with black error bars represent fitted *m* value and 95% confidence interval of the fit. *m* values were fitted from 84 data points. Affinity is measured from fitting all antibody titration curves simultaneously. E: Slopes *m* of four anti-RBD2 mAbs (i: 5G8, ii: CV30, iii: CA11, iv: HL1003) binding to RBD2 WT, variants (α, β, γ, δ, ζ, θ, κ, λ, μ, ο), single point mutants (R346K, K417N/T, L452Q/R, E484K/Q, F490S, N501Y) and negative control (His-GFP-HA). Bars and error bars represent average and standard deviation of *m* values fitted on three different chips with at least 28 data points each, normalized by *m*_*WT*_ on each chip. White: WT, dark blue: variants, light blue: single point mutants, grey: negative control. Shaded grey area: variants not tested with slope quantification. Binding was assessed at fixed Ag synthesis in Supplementary Fig. 10. F: RBD2 sequence shows amino acids that were identified to interact (circle) or not interact (cross) with anti-RBD2 mAbs 5G8, CV30, CA11, HL1003.

The improved chip layout with elongated compartments can accommodate many more different antigens on one chip, without compromising the amount of data points. We therefore used this chip layout to profile the binding of Abs to a large panel of antigenic variants. During the COVID-19 pandemic, new SARS-CoV-2 Variants of Concern (VoC) were constantly emerging with variability in transmissibility, virulence and immune evasion^38^. The Spike region of SARS-CoV-2 is particularly prone to mutations that affect virus fitness^38,39^. We patterned several chips with elongated compartments with DNA brushes encoding the RBD2 (WT) antigen and 5 VoC (Alpha, Beta, Delta, Theta, Omicron). These variants differ from the WT sequence by 1 (Alpha) to 15 (Omicron) point mutations (Supplementary Fig. 10). We further included the single point mutations that constitute each of the Beta, Delta, and Theta variants (Fig. 2E). These point mutations are part of or immediately adjacent to the epitope recognized by this Ab, as determined by X-ray crystallography^40^ (Supplementary Fig. 10).

Post on-chip CFE, each chip was incubated with a different anti-RBD2 mAb, 5G8, CA11 and HL1003, and compared to a more limited variant profile of CV30 obtained on a chip with circular compartments (Fig. 2E). In elongated chips, mAb volume could be minimized to ∼1 µL per compartment. Unlike the CV30 whose epitope has been previously characterized^40^, the RBD2 epitopes recognized by the mAbs 5G8, CA11 and HL1003 have not been characterized so far, to our knowledge. Our epitope profiling (summarized in Fig. 2F) demonstrated that all mutations in the RBD region between amino acids 450 to 500 did not affect binding of 5G8, while mutations of amino acid R346 and K417 abolished its binding entirely, suggesting that the epitope of 5G8 is in the N-ter region of the RBD. CV30 and 5G8 lost their binding to the Beta variant due to the K417N mutation, while CA11 and HL1003 lost Beta binding due to the E484K mutation. CV30 had increased binding to the Delta variant due to L452R, but not T478K, and CA11 binding was sensitive to mutations of amino acids K417, L452, T478 and E484, suggesting that these amino acids are part of this mAb’s epitope. Our data suggest that HL1003’s epitope includes amino acids R346, K417, L452, T478 and F490. Interestingly, for some amino acids, such as K417, the K417N substitution did not affect HL1003 binding, while K417T improved HL1003 binding, revealing that this amino acid is most likely part of the epitope of that mAb. The K417N substitution suggests that the interaction of HL1003 is not with the charge on the side chain of K417 but rather with the long side chain of Lysin (K) and Asparagine (N). The K417T substitution might allow the shorter side chain of threonine (T) to form a hydrogen bond previously not possible.

As all antigen variants on one chip were exposed to the same concentration of mAb, the ratio of the slopes obtained for each variant is proportional to the inverse ratio of the affinities: 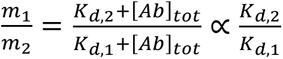, so that 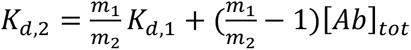. We calculated the relative binding affinity of CV30 to the variants and to the single point mutants compared to WT to be: Alpha 0.30 ± 0.05, Beta 0.024 ± 0.006, Delta 2.7 ± 0.4, Theta 0.23 ± 0.04, Omicron 0.033 ± 0.006 (Supplementary Fig. 10). Huang *et al.* measured a ratio of affinities of 2 for Delta and 0.05 for Omicron compared to WT^41^. To our knowledge, the relative affinities of CV30 to other variants and single point mutants tested here have not been measured before.

### Thorough epitope mapping of by gene truncations

Our approach is based on synthetic DNA, allowing us to easily vary antigen length and sequence without tedious protein purification. Inspired by the fragmentation of pathogen proteins in antigen-presenting cells, we split the full SARS-CoV-2 N gene (419 amino acids aa) into 100 aa-long fragments with a 50 amino acid long overlap (fragments a-h, Fig. 3A). We programmed one chip with DNA encoding all N-fragments and after antigen CFE stained it with two anti-N mAbs: 1A6 (originally raised against the full N protein) and 6H3 (raised against a fragment of N covering aa 121 to 419, see Methods). The chip was stained with the two mAbs simultaneously taking advantage of their orthogonal secondary Abs (Methods). While both mAbs bound strongly to the full N protein (*m*_*Ab*5_ = 0.24 ± 0.04 a.u., *m*_*Ab*4_ = 1.4 ± 0.1 a.u., Fig. 3B, Supplementary Fig. 11), 1A6 showed no recognition of any of the N fragments, whereas 6H3 bound strongly to fragment f, aa 252-351 (*m* = 0.55 ± 0.04 a.u.), did not bind to fragment e, and bound very weakly to fragment g (aa 302-401, *m* = 0.071 ± 0.004 a.u.). According to this analysis it could be deduced that 6H3 recognizes a region within aa 302-351, while 1A6 recognizes structural elements only present in the full tertiary structure, in agreement with the way the two mAbs were raised.

**Figure 3.**
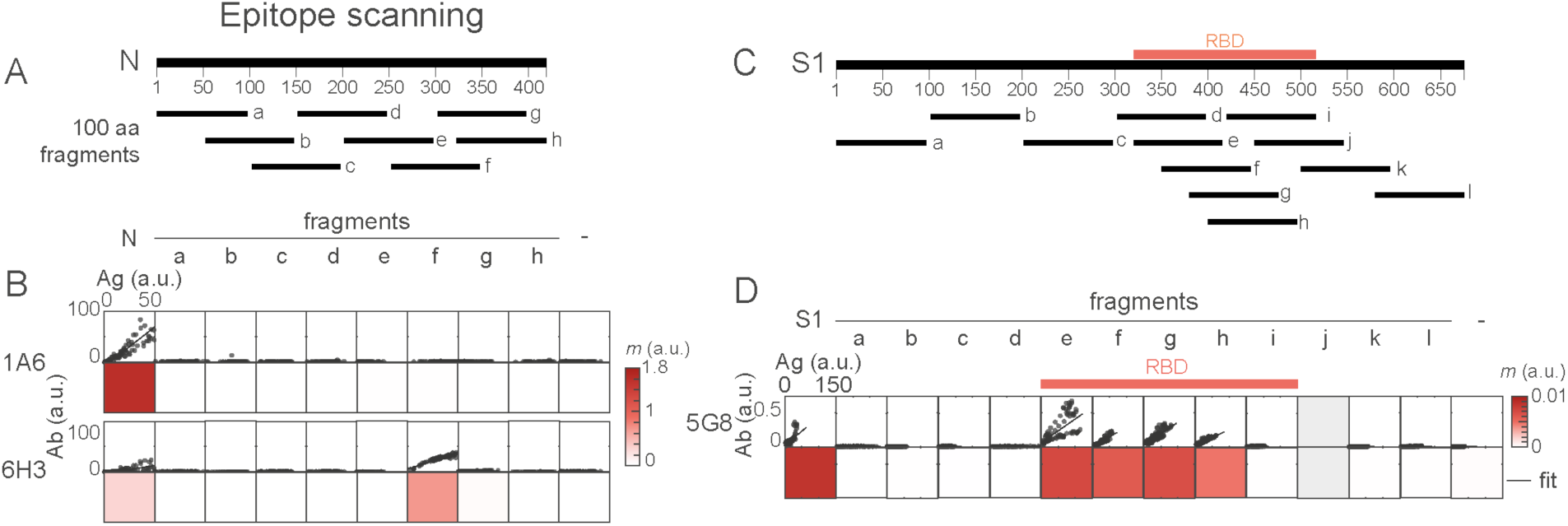
A: Nucleocapsid gene split into 100 amino acids fragments overlapping by 50 aa covering the whole gene. B: Antibody binding for two anti-N mAbs against antigens N, fragments (a-h) and negative control (His-GFP-HA). Dots: data from three chips, lines: fitted slope averaged over three chips. Color: slopes values, *m*, averaged from slope fits of three different chips, each slope fitted from at least 23 data points. Top: anti-N mAb 1A6, bottom: anti-N mAb 6H3. C: S1 gene split into 100 amino acids fragments scanning the RBD2 sequence with 20-30 amino acids overlap and the rest of the sequence with 0-30 amino acids overlap. Fragment j was not tested with 5G8 but used with other Abs (see Supplementary Fig. 11) D: Antibody binding for anti-RBD2 mAb 5G8 against antigens S1, fragments (a-l) and negative control (His-GFP-HA). Dots: data from 1 chip, lines: fitted slope averaged over all data points. Color: slopes values, *m*, each slope fitted from at least 960 data points.

A similar fragmentation of the S1 gene into 100 aa-long fragments revealed binding of 5G8 to the full S1 spike and to 4 fragments within the RBD2, namely fragments e (aa 322-421), f (aa 352-451), g (aa 382-481) and h (aa 402-501), but not to neighboring fragments d (aa 302-401) and i (aa 422-521) (Fig. 3C,D). The common region of fragments that bind 5G8 is aa 402-421, suggesting this specific subsequence of RBD is the epitope of this mAb. mAbs CV30, CA11 and HL1003 did not recognize the full S1 protein nor any of the sub-fragments (Supplementary Fig. 11), although they recognized RBD2 WT and variants.

### Profiling polyclonal Abs in human sera with a broad antigens panel

Next, we investigated whether our platform could detect pAbs in human sera samples. Whole serum is a challenge compared to mAbs, as it contains a spectrum of Abs relating to an individual’s entire immunity history, varying by epitope recognition and relative amounts. To increase the likelihood of identifying as many sera Abs as possible, we patterned chips with a full battery of antigens related to SARS-CoV-2, full and eight fragments of N, full and twelve fragments of S1, as well as some RBD variants (RBD2 WT, Delta, Omicron and RBD1). We analyzed twenty serum samples taken from infected individuals (Methods, PCR-confirmed, sampled between April and August 2020 within 1 to 35 days after onset of the infection) and six negative serum samples predating the COVID-19 outbreak. We applied only 4 µL of each serum per chip (Methods, Supplementary Fig. 12) and following washing, detected the serum Abs with a fluorescently labeled secondary anti-Human IgG Ab. We set a detection threshold at one standard deviation above the mean of the negative control and fitted the Ab signals above this threshold with a linear fit. The vast majority of binding responses were linear, suggesting a saturating Ab concentration in the serum, while some had non-linear binding curves implying a limiting Ab concentration in the serum (all samples’ data in Supplementary Fig. 13). For these limiting concentration cases, the antigen-Ab response was fitted in the low antigen region where the response was still linear.

We found a large diversity in the Ab binding signatures within all 26 serum samples (Fig. 4A, B). Every N and S1 sub-fragment were recognized in at least one patient, while none were recognized in all patients, suggesting that lack of binding of a particular antigen was not due to its poor display but rather for lack of an immune response to this antigen in a particular individual. The strength of antigen recognition (as evident by the slope *m*) was also very variable between samples and between antigens: the N antigens slopes had an 85% coefficient of variation, while the S1/RBD antigens slopes had a 325% coefficient of variation (Supplementary Fig. 14), suggesting that the S1/RBD antigen profiles provide a better signature of a patient-specific immune response.

**Figure 4.**
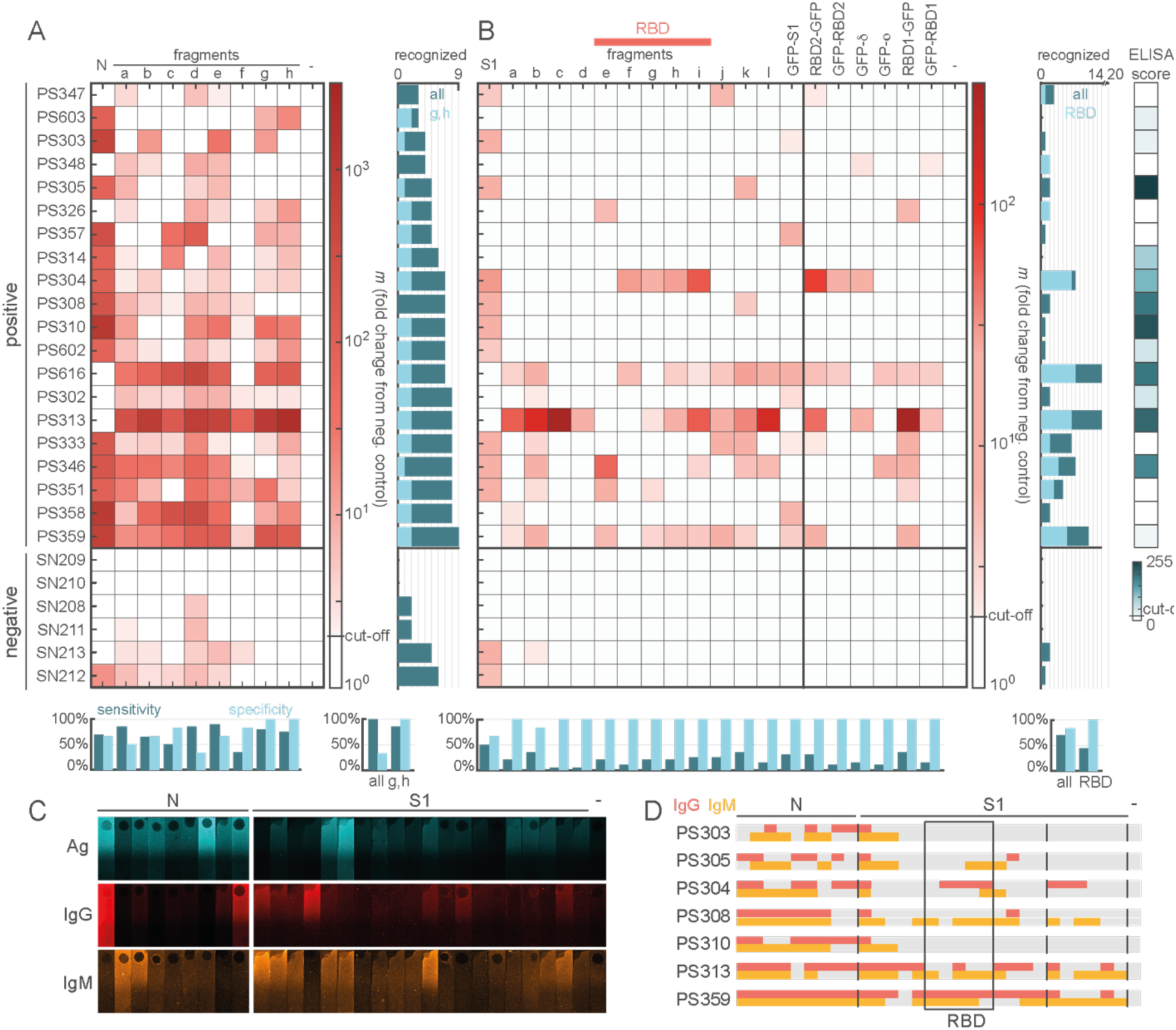
Detection of polyclonal antibodies in human sera. Binding of human sera was assessed against on-chip displayed antigens derived from N (A: N and fragments), S1 and RBD (B: S1 and fragments, RBD and variants). A,B: Fitted slope of antibody-antigen binding, represented as the slope fold change from a negative control (His) on a log scale (shades of red). Data plotted for twenty positive (PS) samples and six negative (SN) samples. Cut-off is set as *m*_*Ag*_ > 2*m*_*neg.control*_. In both A and B, lower bar graph: sensitivity (dark blue) and specificity (light blue) for each antigen; bottom right bar graph: sensitivity and specificity of N (A) and S1/RBD (B) recognition (samples are considered positives if two or more antigens are above cut-off); right hand bar graph: number of antigens recognized in each sample, dark blue: all antigens, light blue: subset of antigens (A: Ng,h, B: all antigens overlapping with RBD). B, right hand box panel: scores of IgG anti-RBD2 from a commercial ELISA test are indicated in shades of blue on a scale from 0 to 255 a.u., with a cut-off at 15 a.u.. ELISA scores for negative samples were not provided. C: Representative microscopy images of sample PS313 antigen synthesis (Ag, blue, top), IgG binding (IgG, red, center), IgM binding (IgM, orange, bottom) for all antigens (as in A and B, N indicates N and fragments, S1 indicates S1, fragments, and RBD variants). Scale for microscopy images: the long axis of one compartment is 750 µm. D: IgG and IgM scores for 7 positive samples. Red and orange indicate above background binding of IgG and IgM respectively. Above background binding is determined by *m*_*Ag*_ > 2*m*_*neg.control*_. Grey indicates below background binding for either antibody. Antigens are as in C. The fragments overlapping with the RBD region are highlighted with a black outline box.

To extract information from our analysis, we calculated the sensitivity and specificity for each antigen individually. Overall sensitivity and specificity across a set of antigens were calculated by considering a sample positive if at least 2 antigens are recognized. While the sensitivity for each N antigen was found to vary between 35% and 90%, the overall sensitivity across N antigens was 100% (Fig. 3A, Supplementary Table 3). The specificity of individual antigens varied between 33% and 100% while the overall N specificity was only 33%. However, the specificity calculated for only the C-ter fragments g and h, was 100%, with an 85% sensitivity. This result complies well with the fact that the N protein is highly homologous among coronaviruses except for its C-ter region^42,43^. The recognition of the full N and its N-ter fragments in pre-pandemic negative samples could therefore indicate an immunity against a previous infection with a coronavirus other than SARS-CoV-2. In comparison to the 100% sensitivity of fragments g and h only, a lateral flow assay using the full N antigen had only a 30% sensitivity for the detection of anti-N IgG for the same set of samples (Methods, specificity could not be measured as negative samples were not tested). This comparison highlights the increased reliability of an extended antigen fragmentation scan compared to a single antigen test.

Almost all S1/RBD antigens had 100% specificity but low sensitivity that varied from 5-50%. The overall sensitivity increased to 70% when calculated based on the recognition of at least two S1/RBD, with an overall specificity of 83% (Fig. 3B, Supplementary Table 3), and comparable to the 70% sensitivity measured by a standard ELISA assay performed on these samples (Methods). Interestingly, we found that our detection of anti-S1/RBD Abs was not correlated with the ELISA scores (Fig. 3B, Supplementary Fig. 14). For example, samples PS347, PS348, PS326, PS333, PS351 and PS358 were below detection with ELISA but showed an immune response to more than two, and up to seven S1/RBD antigens in our assay. Conversely, samples PS305 and PS310 that had the highest ELISA score, showed binding to only two and one antigens, respectively. By presenting a panel of antigens, rather than a single antigen in a standard ELISA, our assay offers a strong validation of immune response by simultaneously detecting several Abs binding to a variety of antigen targets. We further note that the samples presented in Fig. 4A,B were classified according to their response to N antigens (Fig. 4A) demonstrating that samples with a strong immune response to N (Fig. 4A) tended to have a strong response to S1/RBD antigens (Fig. 4B).

In addition, we further analyzed seven of the positive samples with orthogonal anti-IgM secondary Abs (Fig. 4C, Supplementary Fig. 15). IgM Abs are predominant during the early immune response, in the first seven days after the onset of infection, while IgG builds up as a delayed and more sustained immune response^44^. We found no general correlation between IgG and IgM signals (Fig. 4D, Supplementary Fig. 15), and instead observed individual signatures for each sample and each type of Ab. This suggests that despite the presence of both types of Abs in the serum samples, one type does not mask the binding of the other type to the antigens. Rather our assay could reveal a complex and rich picture of patient-specific immune profiles, that goes beyond a simple diagnostic test.

### Reconstitution of Human receptor - viral antigen interaction on chip

The infection of SARS-CoV-2 is initiated by the binding of the Spike’s RBD to the ACE2 receptor displayed on human epithelial cells. The on-chip compartments have been shown previously to provide favorable conditions for reconstitution of complex protein assembly^32^ and we therefore attempted to reconstitute the interaction between RBD2 and ACE2, both expressed from their corresponding genes in the same on-chip compartment. We first verified the correct expression of ACE2 (aa 18-740) in our platform by immobilizing genes coding for ACE2-HA in circular compartments. The surface-captured protein was detected specifically by an anti-ACE2 mAb that has been reported to bind native ACE2 on human cells^45^ (Supplementary Fig. 16) suggesting correct folding of ACE2. We observed colocalization of RBD2 and ACE2 after co-synthesis in circular compartments (Supplementary Fig. 17), however the level of non-specific signals was not negligible, perhaps due to enhanced interactions in the vicinity of a brush coding for two proteins^32^.

To make a more quantitative characterization of the ACE2-RBD interaction we then shifted to elongated compartments and immobilized two DNA brushes, one encoding ACE2-HA, the other RBD2-GFP without an HA tag. The surface of compartments in this set of experiments was fully exposed to UV light (without a pre-patterned protein trap gradient) to maximize capture of all nascent proteins in the compartments. The elongated compartment provided the means to separate the synthesis of the two proteins spatially thereby minimizing non-specific binding. We found that non-specific binding was minimal when the two DNA brushes were immobilized on either side of the compartments (Supplementary Fig. 18, Fig. 5A-C). In this scenario, RBD2 binding occurred on surface captured ACE2 spontaneously forming a surface gradient by diffusion from the DNA brush source (Fig. 5B).

**Figure 5.**
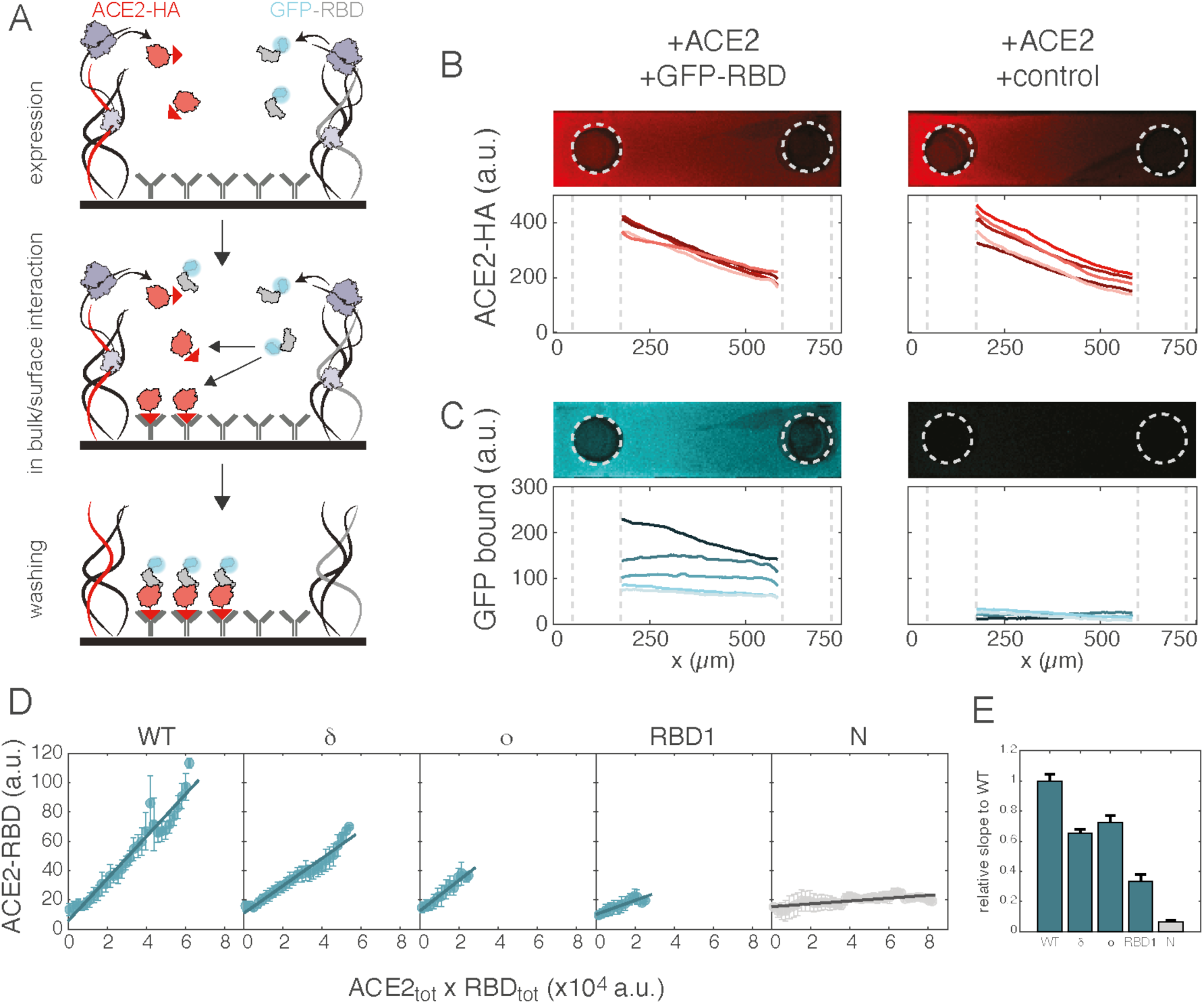
ACE2-RBD interaction. A: DNA brushes coding for ACE2-HA (red) and GFP-RBD (blue) are attached to the edges of a compartment. Cell-free expression, bulk interaction and surface interaction are simultaneous. B: ACE2-HA surface gradient measured by a labeled anti-ACE2 antibody. Microscopy images show one representative compartment, graph shows five compartments spotted with: ACE2-HA and GFP-RBD2 (left panel), ACE2-HA and a control N-GFP (right panel). All microscopy images scale: length of compartment is 750 µm. C: GFP fluorescence after washing of 5 compartments. Microscopy image show one representative GFP-RBD2 compartment. D: Bound complex ACE2-RBD (GFP fluorescence after washing) as a function of local surface available ACE2 (measured by antibody fluorescence) normalized by total RBD synthesis (measured by GFP before washing), for four variants of GFP-RBD and a negative control. Data is fitted with a linear fit, represented by a thick blue line. Data from five separate chips are binned in 50 bins of ACE2-HA antibody fluorescence normalized by total GFP synthesis, dots and error bars represent mean and standard deviation of the bins in the x and y axis. E: Slopes of the linear fit in E normalized by slope of WT.

We quantitatively compared ACE2 binding to the WT RBD2 and to three RBD2 variants: Delta, Omicron and RBD1 on one chip. We first determined the amount of total RBD variants synthesized in each compartment by imaging the GFP signal of the chip before washing (Methods). After chip washing, we determined the amount of bound RBD variants by the GFP signal remaining on the surface and that of nascent ACE2-HA by staining with anti-ACE2 Abs. Under the assumption that the in-situ concentration of all nascent proteins is greater than that of the bound complex, the function of bound ACE2-RBD against total ACE2 normalized by total RBD should be linear with a slope proportional to 1/*K*_*d*_(see Supplementary Information). Using this analysis we indeed obtained a linear function for all RBD variants synthesized on chip (Fig. 5D). We compared the slope obtained for each variant from the above analysis to that of the RBD2 WT (Fig. 5E). The relative slope is proportional to *K*_*d*,*WT*_/*K*_*d*,*var*:*ant*_ and therefore reflects the relative affinity of ACE2 to each variant. We found that in our platform, the affinity of ACE2 to RBD2 Delta, Omicron and RBD1 was respectively ∼0.6, ∼0.7 and ∼0.3 times its affinity for RBD2 WT. Several studies have measured the affinity of ACE2 to RBD and its variants using surface plasmon resonance (SPR) or ELISA, albeit with discrepancies in the relative affinities. One study^46^ found that the *K*_*d*,*app*_ of ACE2 was 1.5 times lower for Delta and 1.4 times lower for Omicron, while another study^47^ found Delta to have 4 times lower *K*_*d*,*app*_ compared to WT, while Omicron was not statistically different from WT. Several studies^48,49^ show no significant difference in the affinity of ACE2 to RBD2 WT, Delta or Omicron. The affinity of ACE2 to RBD2 WT was found to be significantly greater than to RBD1 across studies, but the ratio of affinities varied from 2 to 10^50,51^.

## Discussion

We established a quantitative cell-free platform for the rapid, safe and multiplexed characterization of protein-protein interactions. Specifically, in this study we demonstrated epitope mapping of mAbs, profiling of Abs in human sera, and interaction of a viral protein with a human receptor, both synthesized from their corresponding genes on chip. The mutational analysis with single-point mutation sensitivity, and sub-fragmentation of antigens, allowed us to characterize the amino acid recognition pattern of 5 mAbs whose epitopes have not been established previously, and to profile patient-specific complex immune responses using a broad panel of antigens.

The approach is quantitative using patterned surface gradients of nascent proteins that are formed autonomously and yield dozens of independent binding curves on one chip, bypassing the need for tedious protein purification steps. At a known concentration of mAb, the affinity constant could be deduced from the slope 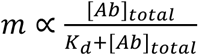 of every mAb-antigen binding response curve in each compartment and mAb titration. For a polyclonal serum sample, Abs differ in their target specificity, affinity, and concentration. Therefore, the slope cannot distinguish between e.g. a high affinity yet dilute Ab to a low affinity yet concentrated Ab. These parameters could be resolved by including an internal and orthogonal reference antigen-Ab pair on the same chip, but the assay remains to be developed.

We used a high-yield *E. coli* CFE system to synthesize the coronavirus antigens and the human ACE2 receptor, which are all mammalian proteins. Despite the lack of mammalian post-translational modifications (PTMs) in the *E. coli* CFE system, we found conditions to reconstitute the antigen-Ab and viral protein-human receptor interactions, and our surface-displayed antigens could be recognized by human serum Abs. *E. coli* CFE systems that support oxidative conditions^24^ required for disulfide bond formation (Supplementary Fig. 8) and specific glycosylation pathways^52^ are being developed, and could be used for human proteins whose structure and function rely on PTMs.

The antigen-presenting biochip is highly relevant for studying emerging pathogens with a very short response time^53^. Since our chips are genetically programmed, a biochip can be engineered to characterize an emerging pathogen based solely on genome sequences as they appear. Our chips could detect pAbs within a few weeks after the onset of an epidemic, allowing to monitor populations and track the spread of infections and mutation evolution, identify the most immunogenic epitopes, and provide information to advance vaccine development and drug leads.

## Materials and Methods

### Chip preparation

The process of chip preparation was previously described in ^32^. Briefly, a Si wafer (University Wafer, USA) was etched in three steps. The patterns were created using AutoCAD software (AutoDesk) in dxf format and converted to CIF format using KLayout (www.klayout.de). Photolithography on the wafer was done using a Microwriter ML3 (Durham Magneto Optics, UK) and etched with an inductively couple plasma (ICP) machine (LPX ICP, SPTS Technologies, UK). For the first step a ∼0.5 µm thick layer of S1805 resist (MicroChemicals, Germany) was spin-coated on the wafer, which was then etched using 10 cycles (circular compartments) or 20 cycles (elongated compartments) of an SF6 etch alternating with C4F8 polymer deposition (Bosch process). This resulted in ∼ 5 µm deep (circular compartments) or ∼10 µm deep (elongated compartments) chambers. Similarly, in the second step a ∼3 µm thick layer of S1818\S1828 resist (MicroChemicals, Germany) was spin-coated on the wafer that was etched in the first step. Following 100 cycles of Bosch process on the ICP the ∼ 50 µm deep separation channels were formed. On the third step a ∼3-9 µm thick layer of S1828\AZ4562 resist (MicroChemicals, Germany) was spin-coated on the back side of the wafer. Following 600-700 cycles of Bosch process etching the outer rim of the chip wafer was divided into separate chips. All heights were measured using a stylus profiler (DktakXT, Dektak/Bruker, USA).

The silicon chips are then coated with a photosensitive and biocompatible monolayer^54^. The monolayer consists of a polymer formed by a polyethylene glycol backbone with a protected amine at one end and a triethoxysilyl group at the other end. The chips are incubated with a 1 mg.ml^−1^ solution of the polymer in dried toluene (244511, Sigma-Aldrich, USA), for 15 min, then rinsed with toluene (Bio-Lab, Israel) and dried. For the elongated compartments, pre-treatment of the surface was done to reduce noise. Chips were incubated for 10 minutes in m-dPEG 4-NHS ester (QBD10211 Sigma-Aldrich) diluted 1:100 v/v in 0.2 M borate-buffered solution pH 8.6 (Thermo Fisher Scientific), rinsed with water and dried. Deprotection of the surface amines is then performed with UV exposition using the MicroWriter ML3. For the circular compartments and full surface exposure of the elongated compartments, the patterns were created using AutoCAD software in dxf format and converted to CIF format using KLayout. For elongated compartments with surface density gradient, a 24 bit RGB PNG was created using MATLAB. The gradient was coded on the Blue channel. The chips were immediately incubated with 0.5 mg/ml biotin 3-sulfo-*N*-hydroxysuccinimide ester (EZ-link NHS biotin, 20217, Thermo Fisher Scientific, USA) in 0.2 M borate-buffered solution pH 8.6 (Thermo Fisher Scientific) for 30 min, then rinsed with water and dried.

### DNA preparation

All coding sequences are placed under a T7 promoter, a strong ribosome binding site (RBS) and are optimized for expression in *E. coli* (see Supplementary Table 1). All genes were ordered as gBlocks from IDT, USA and cloned into a pIVEX2.5 plasmid using Gibson Assembly^55^ (NEBuilder HiFi Assembly Master Mix, E2621, NEB, USA). For ACE2/RBD binding experiments, ACE2 was tagged with a HA peptide tag, and RBD was tagged with a N-ters eGFP. For Ab binding experiments, genes coding for the antigens were placed in our expression cassette: T7 promoter-RBS-[Ag]-eGFP-HA-T7 terminator. All Nucleocapsid (N) related constructs, all Spike 1 (S1) fragments and all control constructs (His, FLAG) were cloned with eGFP at the C-ter of the antigen. All RBD2 variants and single point mutants were cloned with eGFP at the N-ter of the antigen. S1, RBD2 WT and RBD1 constructs were cloned with both positions, eGFP at the N- or at the C-ter. Exact DNA sequences are given in Supplementary Table 1.

Plasmids were transformed into *E. coli* DH5a, purified using Wizard SV-Gel Miniprep (Promega, USA) and DNA concentrations were determined using a NanoDrop (NanoPhotometer, Implen, USA). Linear double-stranded DNA fragments were then amplified with Polymerase Chain Reaction (PCR) with KAPA HotStart ready mix (07958935001, Roche, Switzerland) using a reverse primer conjugated to biotin as previously described^56^, and purified with the Wizard SV-Gel and PCR Clean-Up System (Promega, USA). DNA was then mixed with streptavidin (S0677 or S4762, Sigma-Aldrich) at a 1.4:1 streptavidin:DNA ratio in 1x phosphate-buffered saline (PBS, 02-023-5A, Sartorius, Germany) and 5-7% glycerol (Bio-Lab, Israel), forming a DNA-streptavidin conjugate. The correct amplification of DNA fragments and conjugation with streptavidin were verified with 1% agarose gel electrophoresis.

DNA brushes contained various percentages of the genes of interest, typically 1-50% of the total brush. The brush density was normalized to 150 nM total concentration of DNA–streptavidin conjugates with a non-coding DNA fragment from an unrelated gene amplified without a promoter. Exact brush compositions in all experiments are specified in Supplementary Table 2.

### Chip patterning, expression and staining

#### DNA deposition

The biotinylated compartments of the chips were patterned with DNA–streptavidin conjugate mixes using a sciFLEXARRAYER S3 spotter with a non-coated PDC60 nozzle (Scienion, Germany). For elongated compartments, smaller drops were generated using sciPU Vario (Scienion, Germany). Several compartments (typically ∼10 for circular compartments and ∼3 for elongated ones) were patterned with the same DNA brush mix, and the position of the DNA species across the chip was randomized to average out noise caused by chip inhomogeneities. Microdroplets were incubated for 2 h to overnight at room temperature and 50% humidity to limit evaporation. The chips were then washed with 1x PBS.

#### Immobilization of capture Abs

High affinity biotinylated anti-HA Abs (50 µg/ml, ∼500 nM, 12158167001, Roche, Sigma-Aldrich) were mixed with streptavidin at a concentration ratio between 1.5:1 and 2:1 streptavidin:Abs in 1x PBS and incubated for 30min at 4 °C. The mix was then diluted to 25-50 nM in 1x PBS. 75-100 µl of this solution was incubated on the surface of the chip for 1h at 4 °C, and then washed with 1x PBS. For ACE2/RBD co-expression experiments in circular experiments, the surface was incubated for 30 min at room temperature with 100 µl of a 3% (w/v) BSA (Bovine Serum Albumin, A7030, Sigma-Aldrich), 0.1% (v/v) Tween-20 (P1379, Sigma-Aldrich) solution in 1x PBS (BSA PBS-T) and washed with 1x PBS prior to incubation with anti-HA Abs.

#### Cell-free protein synthesis

All protein synthesis was carried out in an *E. coli* CFE system prepared according to published protocols^57^. The CFE reactions were supplemented with 5 µM 6xHis-GamS and 100 nM 6xHis-T7 RNA Polymerase, purified following published protocols (GamS^58^,T7 RNAP^59^). The chip was rinsed with 1x PBS and excess solution was carefully blotted using paper (Whatman grade 1, 1001-070, Cytiva, USA) without drying the chambers. The chip was rinsed then incubated with 2×50 µl of CFE, excess solution was removed and a 1-2 mm thick polydimethylsiloxane (PDMS) slab (SYLGARD 184 silicone elastomer kit, Dow Corning, USA) previously incubated in a plasma cleaner (PDC-32G-2, Harrick Plasma, USA) for 5 min was applied to seal the chambers. Expression was carried out at temperatures ranging from 16 °C to 30 °C and for times ranging from 30 min to 8 h, in a PCR machine (Labcycler 48, SensoQuest, Germany) fitted with a slide hybridization adapter. After expression, the PDMS slab was removed, and the chip was washed with 1× PBS. For ACE2/RBD experiments in elongated compartments, the total GFP synthesis in each compartment was measured by imaging chips through the PDMS before opening them. As a negative control for synthesis, a non-fluorescent, HA-tagged protein was synthesized in selected compartments (NC-HA): the coding sequence corresponds to the tail tubular protein of the T4 phage, gp11^32^, and is unrelated to SARS-CoV.

#### Ab staining

Chips were blocked following expression by overnight incubation at 4 °C in 1-2 ml BSA PBS-T. All Ab incubation steps were conducted for 1h at room temperature in 2 ml (1 ml for elongated compartments) of a 1 µg/ml Ab dilution in BSA PBS-T or as stated in Supplementary Table 2. Between incubation of the primary and the secondary Abs, the chips were washed 3 times for 20 min in 2 ml PBS-T (0.1% (v/v) Tween-20 in 1x PBS). After staining with the secondary Ab, the chips were washed in PBS-T and imaged. Anti-N mAb 1A6 titration (Fig. 1C, Supplementary Fig. 4) was conducted by diluting the primary mAb in BSA PBS-T, fetal bovine serum (FBS), or human serum (S1-100 mL, Sigma-Aldrich, Germany). Titrations with anti-RBD1 mAb CR3022 (Fig. 1H) and anti-N mAb 6H3 (Supplementary Fig. 11) were conducted by dilutions the primary mAb in BSA PBS-T. Primary and secondary mAbs were purchased from Abcam (UK) and Jackson ImmunoResearch (USA), and are listed below:

Primary Abs: monoclonal Mouse anti-6x His tag DyLight650 (ab117504), monoclonal Rabbit anti-FLAG Alexa647 (ab245893), monoclonal Chimeric anti-SARS-CoV-2 Nucleocapsid 1A6 (ab272852), monoclonal Mouse anti-SARS Nucleocapsid 6H3 (ab273434), monoclonal Human anti-SARS-CoV-2 Spike RBD CV30 (ab277513), monoclonal Mouse anti-SARS-CoV-2 Spike RBD 5G8 (ab277628), monoclonal Human anti-SARS-CoV-1 Spike Glycoprotein S1 CR3022 (ab273073), monoclonal Rabbit anti-SARS-CoV-2 Spike RBD HL1003 (ab281303), monoclonal Rabbit anti-SARS-CoV-2 Spike RBD CA11 (ab284651), monoclonal Rabbit anti-ACE2 (ab108252).

Secondary Abs, used accordingly to primary Abs: Donkey anti-Rabbit Alexa647 (ab150075), Goat anti-Rabbit Alexa594 (ab150080), Goat anti-Human DyLight650 (ab97006), Goat anti-Human DyLight550 (ab97004), Goat anti-Mouse DyLight650 (ab97018) and AffiniPure Goat Anti-Human IgM Cy3 (109-165-129).

#### Negative GFP staining

On-chip protein synthesis of non-fluorescently labeled proteins (ACE2-HA, NC-HA, Supplementary Fig. 16) was quantified using crude *E. coli* lysate with overexpressed recombinant GFP-HA gene, prepared as follows: A GFP-HA gene under a T7 promoter was transformed in BL21 (DE3) *E. coli* cells. A single colony was used to inoculate an overnight culture in LB medium with the appropriate antibiotic resistance. A large volume of LB medium (> 1 L) was then inoculated with the overnight culture, and protein synthesis was induced with 0.1 mM IPTG once OD reached 0.8. Cells were further grown at 37 °C for 3 h. Cells were then centrifuged, resuspended in a PBS buffer and lysed with sonication. After centrifugation of the cell debris, the supernatant (GFP crude lysate) was aliquoted and stored at −20 °C. GFP concentration in the crude lysate was estimated with absorbance at 495 nm. Chips were incubated in 2 mL of 50 nM GFP diluted in PBS buffer at room temperature for 10 min, then washed and imaged. The GFP-HA binds to surface anti-HA Abs that were left unbound during the CFE reaction, thereby providing a negative image of the occupied sites.

### Human samples

Serum samples were obtained from RayBiotech, CoV-PosSet-S1 (Georgia, USA). Anti-N IgG and IgM were detected by the company with lateral flow assay, anti-RBD2 IgG and IgM were detected and quantified by the company with ELISA. All positive samples but one were tested positive with PCR, one was tested positive with Abs. Negative samples were taken pre-pandemic and were not otherwise tested. The study was conducted according to the guidelines of the Declaration of Helsinki and collection and use of clinical samples was approved by IRB (ID# 8291-BZhang). Written informed consent was obtained from all subjects involved in the study.

For on chip measurement, 4 µL of serum sample were mixed with 8 µL of BSA PBS-T buffer and 4 µL of crude NC-HA lysate, prepared as described above for GFP crude lysate. The serum and NC-HA mixture was incubated for 10 min at room temperature. The goal of this step was to capture anti-HA antibodies present in the serum and prevent them from forming a background binding to surface captured HA tags. The HA tag is derived from the human influenza hemagglutinin protein, and therefore human sera are likely to present antibodies recognizing this peptide. Although different serum samples showed different background binding to HA tags, spiking the serum with NC-HA alleviated this binding and reduced false positive detection of antigens (Supplementary Fig. 12). Following this incubation, the serum was diluted to 100 µL total volume with BSA PBS-T, and was incubated for 2 h at room temperature on the chip. The chips were then washed with PBS-T and incubated with a secondary antibody as described above.

### Fluorescent microscopy imaging

Fluorescent images were obtained using two microscopes:

AxioObserver Z1 inverted microscope with a motorized stage (Zeiss) and Plan-Apochromat 20×/0.8 M27 (Olympus) objective. Illumination was performed using a Colibri2 LED illumination system equipped with a 470-nm, 555-nm, and 625-nm LED module (Zeiss) and filter set 38 HE (Zeiss; excitation 470/40nm, dichroic mirror 495nm, emission 525/50nm), filter set 43 HE (Zeiss; excitation 550/25nm, dichroic mirror 570nm, emission 605/70nm) and filter set 50 (Zeiss; excitation 640/30nm, dichroic mirror 660nm, emission 690/50nm) respectively. Images were captured using an iXon Ultra CCD camera (Andor Technology, Belfast, UK). Chip alignment and multi-image acquisition was performed using the Zeiss ZEN 2012 software.

AxioZoom V16 stereo zoom microscope with a motorized stage (Zeiss, Germany) and ApoZ 1.5× 10.37 FWD 30mm (Zeiss) objective. Illumination was performed using a Zeiss Illuminator HXP 200C equipped with filter set 38 (Zeiss; excitation 470/40nm, dichroic mirror 495nm, emission 525/50nm), filter set 43 HE (Zeiss; excitation 550/25nm, dichroic mirror 570nm, emission 605/70nm) and filter set 50 (Zeiss; excitation 640/30nm, dichroic mirror 660nm, emission 690/50nm) respectively. Images were captured using a Zeiss Axiocam 712 monochrome camera. Chip alignment and multi-image acquisition was performed using the Zeiss ZEN 3.4 pro software.

### Data analysis

For both circular and elongated compartments, fluorescence intensity data was extracted with an in-house script that cross-correlated the compartment’s image to a reference image to identify the circular or linear region of above-background fluorescence.

For circular compartments, the average intensity of this circular region was subtracted by its local background, the average intensity of the area outside the circle. Fluorescence intensity was normalized by exposure time and gain. All compartments containing a given DNA specie were averaged, and the average of the relevant negative control (gp11 for GFP synthesis, a non-binding antigen for Ab binding) was subtracted from all fluorescence intensity values.

For elongated compartments, the fluorescence intensity was summed on the short axis to extract a one-dimensional vector along the long axis. The fluorescence intensity of a negative control (gp11 for both GFP synthesis and Ab binding) was averaged over the long axis and over all compartments. This was taken as background and all fluorescence intensities were subtracted by this value.

### Bulk characterization

Expression was characterized in bulk in three types of experiment (Supplementary Fig. 6). Expression in *E. coli* CFE system at 30 °C was conducted in a ClarioStar plate reader (BMG Labtech, Germany). 1 nM of Streptavidin conjugated DNA was added to the CFE reaction supplemented with 5 µM 6xHis-GamS and 100 nM 6xHis-T7 RNA Polymerase. Volumes of 10 µL were pipetted in a black flat-bottomed 384-well plate and spun down at 1,000 rcf for 2 min. GFP fluorescence was measured with excitation filter 470/15 nm, dichroic filter 491 nm, and emission filter 515-20 nm.

Expression in human cell extract at 30 °C was conducted in a ClarioStar plate reader (BMG Labtech, Germany). 40 ng/µL of plasmid DNA was added to a 1-Step Human Coupled IVT Kit (catalog number: 88881, Thermo Fisher Scientific, USA) according to the kit specifications. Volumes of 10 µL were pipetted in a black flat-bottomed 384-well plate and spun down at 1,000 rcf for 2 min. GFP fluorescence was measured with excitation filter 470/15nm, dichroic filter 491 nm, and emission filter 515-20 nm.

*E. coli* CFE at temperatures varying from 16 °C to 30 °C was conducted in a StepOnePlus (Applied Biosystems, USA) real-time PCR. 1 nM of Streptavidin conjugated DNA was added to the CFE reactions supplemented with 5 µM 6xHis-GamS and 100 nM 6xHis-T7 RNA Polymerase. Volumes of 10 µL were pipetted in a white v-bottomed 96-well plate. GFP fluorescence was measured with the FAM fluorescent channel (excitation: 493 nm, emission: 517 nm).

### Protein synthesis and solubility characterization

Synthesis and solubility of proteins was assessed using SDS-Poly Acrylamide-Gel-Electrophoresis (PAGE) (Supplementary Fig. 6). Proteins were synthesized in an *E. coli* CFE system as described above. A sample of the whole CFE reaction was collected, and the rest was centrifuged at 30,000×g for 20 min at 4 °C. The supernatant and the pellet were collected separately. All samples were diluted or resuspended (for the pellet) in 1x sample buffer (4x sample buffer: 40% glycerol v/v, 8% SDS w/v, 400 mM DTT, 200 mM Tris pH 6.8, 0.1% bromo-phenol blue). A 4-20% PAGE-gel (Gene Bio-Application, Israel) was pre-run at 160V for 12 min. Samples were loaded unto the gel together with a protein marker (PM2700, SMOBIO, Taiwan) and ran at 160V for 70 min. The gel was imaged with a Typhoon FLA 9500 laser scanner (GE Healthcare, USA) in the GFP channel (excitation: 473 nm, filter LPB) and the far-red channel (excitation: 635 nm, filter LPR).

### Disulfide bond formation

To enable the formation of disulfide bonds, iodoacetamide (I1149, Sigma-Aldrich, USA) was added to the CFE reaction (naturally reducing) to a final concentration of 0.1 to 0.5 mM (Supplementary Fig. 8). Correct formation of disulfide bonds in this environment was assessed with synthesis of Gaussia luciferase and luminescence measurement with a Pierce Gaussia Luciferase Glow Assay Kit (16160, Thermo Fisher Scientific, USA). The sequence of RBD2 used throughout this work contains seven Cysteine. To enable an additional disulfide bond to form, an elongated sequence of RBD2 with an extra Cysteine codon was cloned (Supplementary Fig. 8, Supplementary Table 1).

### Affinity measurement with biolayer interferometry

On-chip measured affinity of antigen-Ab pairs was confirmed with biolayer interferometry (Supplementary Fig. 5) using an Octet RED96e (Sartorius, Germany). Samples were dispensed into 96-well microtiter plates (655209 Greiner) at a volume of 160 μL per well. Operating temperature was maintained at 30 °C. Streptavidin-coated biosensor tips (18-009 Sartorius, Germany) were dipped into assay buffer (1x PBS, 0.1% BSA, 0.02% Tween-20) for 60 s to establish a baseline. All solutions were diluted in assay buffer, and all steps were carried oud while agitating at 1000 rpm. The tips were then loaded with the antigen by serial dipping into the following solutions: 2.5 µg/mL biotinylated anti-HA Abs 240 s, assay buffer 60 s, 100 nM antigen-HA (synthesized overnight at 16 °C in cell-extract with 1 nM plasmid, quantified by GFP fluorescence) 240 s, assay buffer 60 s. Ab association to the antigen was then measured by dipping tips in an Ab solution, ranging from 0 nM to 400 nM Ab for 100 s. Ab dissociation was measured by dipping the tips in assay buffer for 180 s. Tips were recycled by 3 cycles of regeneration buffer (glycine HCl 10 mM pH 1.7) and assay buffer, and were reused until anti-HA attachment could not be detected. Negative controls validating the absence of Ab association to non-cognitive antigens, tips not coated with anti-HA or tips not coated with antigens were conducted. Data were generated automatically by the Octet User Software (version 3.1) and were subsequently plotted with Matlab_R2020a. Kinetics and affinity fits were performed in the software.

## Acknowledgments

The authors thank Moshe Goldsmith for his help with the Octet experiments and Yoav Barak for support with biochemical characterization. A.D. acknowledges funding from the EMBO postdoctoral fellowship, award number: ALTF 131-2020. This study was partly supported by the generous donation from Miel de Botton. We acknowledge funding from the Israel Science Foundation (R.B.Z. and S.S.D., grant no. 3435/24), the United States-Israel Binational Science Foundation (R.B.Z., S.S.D. and V.N. grant no. 2022385), the United States Department of Defense (R.B.Z and S.S.D, Agency Ref. No. W911NF2010119), the Isak Ferdinand and Dwosia Artmann Research Fund for Biological Physics and the Weizmann Institute office for Technology Transfer (R.B.Z. and S.S.D.).

## Authors contributions

A.D. designed and conducted all experiments, data analysis and data visualization, cloned constructs, prepared DNA and silicon chips. O.V. designed and conducted experiments, conceptualized and developed the continuous surface density gradient, prepared DNA and silicon chips, prepared DNA and supported experiments. V.A. conducted experiments on IgG and IgM detection in human sera, and supported DNA preparation. M.L. developed data extraction code for elongated compartments, Y.D. developed data extraction code for circular compartments. N.A., A.D. and S.S.D cloned constructs. S.P. produced elongated silicon chips. S.T. and V.N. provided the CFE system, as well as Gaussia luciferase plasmid. A.D., O.V., S.S.D. and R. B.-Z. conceptualized the project. A.D. and S.S.D. co-wrote the paper. All authors reviewed the manuscript.

## References

1. Yuan, J. et al. Novel technologies and emerging biomarkers for personalized cancer immunotherapy. J Immunother Cancer 4, (2016).

2. Connors, J. et al. Using the power of innate immunoprofiling to understand vaccine design, infection, and immunity. Hum Vaccin Immunother 19, (2023).

3. Ogunniyi, A. O. et al. Profiling human antibody responses by integrated single-cell analysis. Vaccine 32, 2866–2873 (2014).

4. Stork, E. M. et al. Antigen-specific Fab profiling achieves molecular-resolution analysis of human autoantibody repertoires in rheumatoid arthritis. Nature Communications 2024 15:1 15, 1–12 (2024).

5. Lu, R. M. et al. Development of therapeutic antibodies for the treatment of diseases. Journal of Biomedical Science 2020 27:1 27, 1–30 (2020).

6. Deshaies, R. J. Multispecific drugs herald a new era of biopharmaceutical innovation. Nature 580, 329–338 (2020).

7. Zhong, X., D’antona, A. M., Karagiannis, S. & White, A. Recent Advances in the Molecular Design and Applications of Multispecific Biotherapeutics. Antibodies 2021, Vol. 10, Page 13 10, 13 (2021).

8. Weiner, G. J. Building better monoclonal antibody-based therapeutics. Nature Reviews Cancer 2015 15:6 15, 361–370 (2015).

9. Gérard, A. et al. High-throughput single-cell activity-based screening and sequencing of antibodies using droplet microfluidics. Nature Biotechnology 2020 38:6 38, 715–721 (2020).

10. Muruato, A. E. et al. A high-throughput neutralizing antibody assay for COVID-19 diagnosis and vaccine evaluation. Nature Communications 2020 11:1 11, 1–6 (2020).

11. Tickle, S. et al. High-Throughput Screening for High Affinity Antibodies. J Lab Autom 14, 303–307 (2009).

12. Schofield, D. J. et al. Application of phage display to high throughput antibody generation and characterization. Genome Biol 8, 1–18 (2007).

13. Cubillos-Ruiz, A. et al. Engineering living therapeutics with synthetic biology. Nat Rev Drug Discov 20, 941–960 (2021).

14. Gu, Y. et al. Cell-free protein synthesis system for bioanalysis: Advances in methods and applications. TrAC Trends in Analytical Chemistry 161, 117015 (2023).

15. Slomovic, S., Pardee, K. & Collins, J. J. Synthetic biology devices for in vitro and in vivo diagnostics. Proc Natl Acad Sci U S A 112, 14429–14435 (2015).

16. Murray, C. J. & Baliga, R. Cell-free translation of peptides and proteins:from high throughput screening to clinical production. Curr Opin Chem Biol 17, 420–426 (2013).

17. Ramm, F. et al. Mammalian cell-free protein expression promotes the functional characterization of the tripartite non-hemolytic enterotoxin from Bacillus cereus. Scientific Reports 2020 10:1 10, 1–12 (2020).

18. Thoring, L., et al. Cell-Free Systems Based on CHO Cell Lysates: Optimization Strategies, Synthesis of “Difficult-to-Express” Proteins and Future Perspectives. PLoS One 11, e0163670 (2016).

19. Garenne, D., Bowden, S. & Noireaux, V. Cell-free expression and synthesis of viruses and bacteriophages: applications to medicine and nanotechnology. Curr Opin Syst Biol 28, 100373 (2021).

20. Sullivan, C. J. et al. A cell-free expression and purification process for rapid production of protein biologics. Biotechnol J 11, 238–248 (2016).

21. Pardee, K. et al. Rapid, Low-Cost Detection of Zika Virus Using Programmable Biomolecular Components. Cell 165, 1255–1266 (2016).

22. Gootenberg, J. S. et al. Nucleic acid detection with CRISPR-Cas13a/C2c2. Science (1979) 356, 438–442 (2017).

23. van Dongen, J. E. et al. Point-of-care CRISPR/Cas nucleic acid detection: Recent advances, challenges and opportunities. Biosens Bioelectron 166, 112445 (2020).

24. Hunt, A. C. et al. A rapid cell-free expression and screening platform for antibody discovery. Nature Communications 2023 14:1 14, 1–14 (2023).

25. Ojima-Kato, T., Nagai, S. & Nakano, H. Ecobody technology: rapid monoclonal antibody screening method from single B cells using cell-free protein synthesis for antigen-binding fragment formation OPEN. doi:10.1038/s41598-017-14277-0.

26. Stech, M. & Kubick, S. Cell-Free Synthesis Meets Antibody Production: A Review. Antibodies 4, 12–33 (2015).

27. Norouzi, M. et al. Cell-Free Dot Blot: an Ultra-Low-Cost and Practical Immunoassay Platform for Detection of Anti-SARS-CoV-2 Antibodies in Human and Animal Sera. Microbiol Spectr 11, (2023).

28. Hufnagel, K. et al. In situ, Cell-free Protein Expression on Microarrays and Their Use for the Detection of Immune Responses. Bio Protoc 9, (2019).

29. Goshima, N. et al. Human protein factory for converting the transcriptome into an in vitro-expressed proteome. Nat Methods 5, 1011–1017 (2008).

30. Morishita, R. et al. CF-PA2Vtech: a cell-free human protein array technology for antibody validation against human proteins. Scientific Reports 2019 9:1 9, 1–10 (2019).

31. Levy, M., Falkovich, R., Daube, S. S. & Bar-Ziv, R. H. Autonomous synthesis and assembly of a ribosomal subunit on a chip. Sci Adv 6, eaaz6020 (2020).

32. Vonshak, O. et al. Programming multi-protein assembly by gene-brush patterns and two-dimensional compartment geometry. Nat Nanotechnol 1–9 (2020) doi:10.1038/s41565-020-0720-7.

33. Caschera, F. & Noireaux, V. Synthesis of 2.3 mg/ml of protein with an all Escherichia coli cell-free transcription-translation system. Biochimie 99, 162–8 (2014).

34. Buxboim, A., Daube, S. S. & Bar-Ziv, R. Synthetic gene brushes: a structure-function relationship. Mol Syst Biol 4, 181 (2008).

35. Abcam. Recombinant Anti-SARS-CoV-2 nucleocapsid protein antibody [1A6] - Chimeric (ab272852). https://www.abcam.com/products/primary-antibodies/sars-cov-2-nucleocapsid-protein-antibody-1a6-chimeric-ab272852.html?productWallTab=ShowAll.

36. Klausberger, M. et al. A comprehensive antigen production and characterisation study for easy-to-implement, specific and quantitative SARS-CoV-2 serotests. EBioMedicine 67, (2021).

37. Zhang, S., Garcia-D’Angeli, A., Brennan, J. P. & Huo, Q. Predicting detection limits of enzyme-linked immunosorbent assay (ELISA) and bioanalytical techniques in general. Analyst 139, 439–445 (2013).

38. Telenti, A., Hodcroft, E. B. & Robertson, D. L. The Evolution and Biology of SARS-CoV-2 Variants. Cold Spring Harbor Perspectives in Medecine 12, (2022).

39. Gong, Y., Qin, S. & Dai, L. The glycosylation in SARS-CoV-2 and its receptor ACE2. Signal Transduct Target Ther 6, 396 (2021).

40. Hurlburt, N. K. et al. Structural basis for potent neutralization of SARS-CoV-2 and role of antibody affinity maturation. Nat Commun (2020) doi:10.1038/s41467-020-19231-9.

41. Huang, M. et al. Atlas of currently available human neutralizing antibodies against SARS-CoV-2 and escape. Immunity 1501–1514 (2022) doi:10.1016/j.immuni.2022.06.005.

42. Lee, C. H. et al. Potential CD8+ T Cell Cross-Reactivity Against SARS-CoV-2 Conferred by Other Coronavirus Strains. Front Immunol 11, 1–10 (2020).

43. Wu, W., Cheng, Y., Zhou, H., Sun, C. & Zhang, S. The SARS-CoV-2 nucleocapsid protein: its role in the viral life cycle, structure and functions, and use as a potential target in the development of vaccines and diagnostics. Virol J 20, 1–16 (2023).

44. Movsisyan, M. et al. Tracking the evolution of anti-SARS-CoV-2 antibodies and long-term humoral immunity within 2 years after COVID-19 infection. Sci Rep 14, 1–13 (2024).

45. Abcam. Recombinant Anti-ACE2 antibody [EPR4435(2)] (ab108252). https://www.abcam.com/products/primary-antibodies/ace2-antibody-epr44352-ab108252.html#lb.

46. Mannar, D. et al. SARS-CoV-2 Omicron variant: Antibody evasion and cryo-EM structure of spike protein – ACE2 complex. Science (1979) 764, 760–764 (2022).

47. Wu, L. et al. SARS-CoV-2 Omicron RBD shows weaker binding affinity than the currently dominant Delta variant to human ACE2. Signal Transduct Target Ther 7, 8 (2022).

48. Han, P. et al. Receptor binding and complex structures of human ACE2 to spike RBD from omicron and delta SARS-CoV-2. Cell 185, 630–640.e10 (2022).

49. Li, L. et al. Structural basis of human ACE2 higher binding affinity to currently circulating Omicron SARS-CoV-2 sub-variants BA.2 and BA.1.1. Cell 185, 2952–2960.e10 (2022).

50. Nguyen, H. L. et al. Does SARS-CoV-2 Bind to Human ACE2 More Strongly Than Does SARS-CoV? J Phys Chem B 124, 7336–7347 (2020).

51. Starr, T. N. et al. Deep Mutational Scanning of SARS-CoV-2 Receptor Binding Domain Reveals Constraints on Folding and ACE2 Binding. Cell 182, 1295–1310.e20 (2020).

52. Kightlinger, W. et al. A cell-free biosynthesis platform for modular construction of protein glycosylation pathways. Nat Commun 10, (2019).

53. Marani, M., Katul, G. G., Pan, W. K. & Parolari, A. J. Intensity and frequency of extreme novel epidemics. PNAS 118, (2021).

54. Buxboim, A. et al. A Single-Step Photolithographic Interface for Cell-Free Gene Expression and Active Biochips. Small 3, 500–510 (2007).

55. Gibson, D. G. et al. Enzymatic assembly of DNA molecules up to several hundred kilobases. Nat Methods 6, 343–345 (2009).

56. Buxboim, A., Daube, S. S. & Bar-Ziv, R. Ultradense Synthetic Gene Brushes on a Chip. Nano Lett 9, 909–913 (2009).

57. Caschera, F. & Noireaux, V. Synthesis of 2.3 mg/ml of protein with an all Escherichia coli cell-free transcription-translation system. Biochimie 99, 162–168 (2014).

58. Garamella, J., Marshall, R., Rustad, M. & Noireaux, V. The All E. coli TX-TL Toolbox 2.0: A Platform for Cell-Free Synthetic Biology. ACS Synth Biol 5, 344–355 (2016).

59. He, B. et al. Rapid Mutagenesis and Purification of Phage RNA Polymerases. Protein Expr Purif 9, 142–151 (1997).

