## Supplementary Information for "Cell-free immuno-profiling on a genetically programmed biochip"

This Supplementary Information contains:

- Supplementary text describing models and calculations used in the main text.
- 18 Supplementary figures.
- 3 Supplementary tables (1 is separately attached).

### Calculation of effective DNA concentration in a compartment

The DNA brush has a density of ~ 1000 molecules/μm$^2$ [1], and the radius of the brush is ~ 50 μm. The number of DNA molecules in the compartment is

$$1000 \frac{molecules}{\mu m^2} \times \pi \times (50)^2 \mu m^2 \approx 1.3 \times 10^{-17}\, mol$$

The compartment has a radius of 150 μm and a height of 5 μm. Its volume is

$$\pi \times 150^2 \times 5 \mu m^3 \approx 0.4\, nL$$

The effective DNA concentration is therefore

$$C = \frac{n}{V} \approx 37\, nM$$

Our genes are on average 2000 base pairs long, so the molecular weight of a DNA molecule is approximately 1,300 kDa. The effective DNA concentration of ~ 37 nM corresponds to a mass concentration of ~ 0.05 mg/ml, which is 200 times more dilute than DNA in either eukaryotic[2] or prokaryotic[3] cells.

To calculate the local effective concentration of DNA in the brush, we consider that a 2000 base pairs long linear double stranded DNA has a length of around 170 nm in the CFE conditions[4], and that the DNA brush is roughly cylindrical. The volume of the DNA brush is

$$\pi \times 150^2 \times 0.17 \mu m^3 \approx 12 pL$$

So that the local DNA concentration is ~1 µM.

### Diffusion and binding of proteins in a compartment

The characteristic diffusion time $\tau$ of a protein of diffusion coefficient $D$ in a compartment depends on its typical dimension $L$:

$$\tau \sim \frac{L^2}{D}$$

Assuming a typical protein weight of ~ 50 kDa, we consider an approximate diffusion coefficient in the CFE reaction of $D \approx$ 50 µm$^2$/sec [5].

In the circular compartments of radius $r$=150 µm and height $z$=5 µm, the characteristic diffusion times of a protein is minutes along the radial dimension and seconds along the vertical dimension.
In the elongated compartments of height $z$=10 µm, width $y$= 200 µm and length $x$=750 µm, the characteristic diffusion times of a protein seconds along the vertical dimension, minutes along the width dimension, and hours along the length dimension.

The association rate of a HA-tagged protein binding to a surface-bound anti-HA antibody is in the order of $k_{on}$ ≈$10^5$ /sec/M [6]. Similarly, the binding of RBD2 to ACE2 was measured to have an association rate of $k_{on}$ ≈$10^5$-$10^6$ /sec/M [7].

### Derivation of the antigen-antibody binding chemical equilibrium

We consider the following chemical equilibrium:

$$A + B \rightleftharpoons AB$$

with $A$ the antigen, $B$ the antibody, $AB$ the complex for which the dissociation constant is:

$$K_d = \frac{[A]_{eq} \times [B]_{eq}}{[AB]_{eq}}$$

with $eq$ denoting equilibrium, when the reaction is complete.

We are varying the total amount of antigen available and measuring the bound antibody-antigen, assuming equilibrium. Therefore, we set:

$$[A]_{tot} = [A]_{eq} + [AB]_{eq}$$
$$[B]_{tot} = [B]_{eq} + [AB]_{eq}$$
$$x = [A]_{tot}, y = [AB]_{eq}$$

The general dependency between these two values is:

$$y = \frac{1}{2}\left(x + [B]_{tot} + K_d - \sqrt{(x + [B]_{tot} + K_d)^2 - 4 \times x \times [B]_{tot}}\right)$$

We consider four limit cases:

1. $x, [B]_{tot} \gg K_d$: the reaction is complete
   a. $x \gg [B]_{tot}$

$B$ is the limiting reagent. At equilibrium, $[A]_{eq} \approx [A]_{tot} = x$, $[B]_{eq} \approx 0$ and $[AB]_{eq} = y \approx [B]_{tot}$.

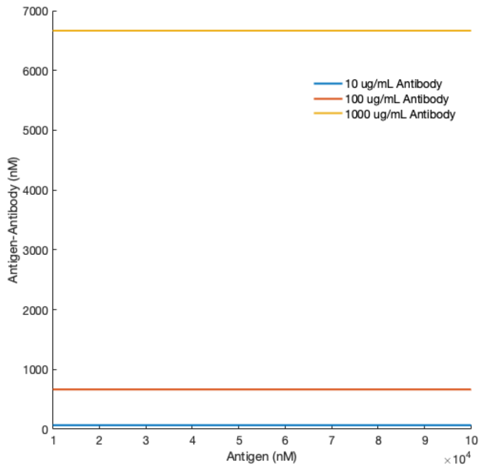


Modeling with $K_d$=1 nM, 100 nM ≤ $x$ ≤ 1 μM, $[B]_{tot}$= 10, 100, 1000 μg/mL ≈ 70, 700, 7000 nM.

   b. $[B]_{tot} \gg x$

$A$ is the limiting reagent. At equilibrium, $[A]_{eq} \approx 0$, $[B]_{eq} \approx [B]_{tot}$ and $[AB]_{eq} = y \approx [A]_{tot} = x$.

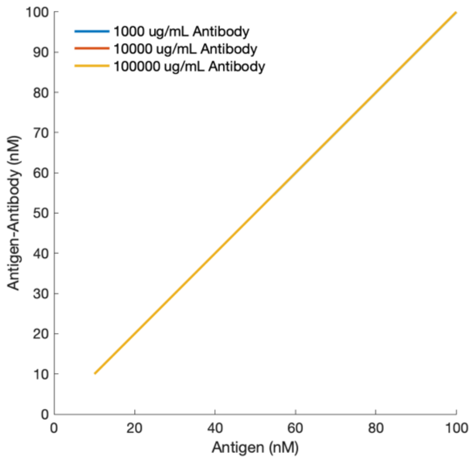


Modeling with $K_d$=1 nM, 10 nM ≤ $x$ ≤ 100 nM, $[B]_{tot}$= 1, 10, 100 mg/mL ≈ 7, 70, 700 μM.

2. $x, [B]_{tot} <$ or $\sim K_d$: the reaction is not complete
   c. $x \gg [B]_{tot}$

At equilibrium, $[A]_{eq} \approx [A]_{tot} = x$, $[B]_{eq} = [B]_{tot} - y$ and $[AB]_{eq} = y$.

The dissociation constant becomes:

$$K_d \approx \frac{x \times ([B]_{tot} - y)}{y}$$

so that:

$$y \approx [B]_{tot} \times \frac{x}{K_d + x}$$

$y$ is a Hill function of $x$ with cooperativity 1.

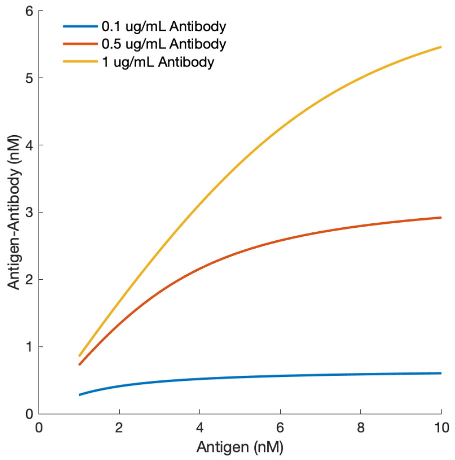


Modeling with $K_d$=1 nM, 1 nM $\leq x \leq$ 10 nM, $[B]_{tot}$= 0.1, 0.5, 1 µg/mL ≈ 0.7, 3, 7 nM.

d. $[B]_{tot} \gg x$

At equilibrium, $[A]_{eq} = x - y$, $[B]_{eq} \approx [B]_{tot}$ and $[AB]_{eq} = y$.

The dissociation constant becomes:

$$K_d \approx \frac{(x - y) \times [B]_{tot}}{y}$$

so that:

$$y \approx \frac{[B]_{tot}}{K_d + [B]_{tot}} \times x$$

$y$ is linearly correlated to $x$, and the slope is a Hill function of $[B]_{tot}$ with cooperativity 1, the total concentration of antibodies incubated.

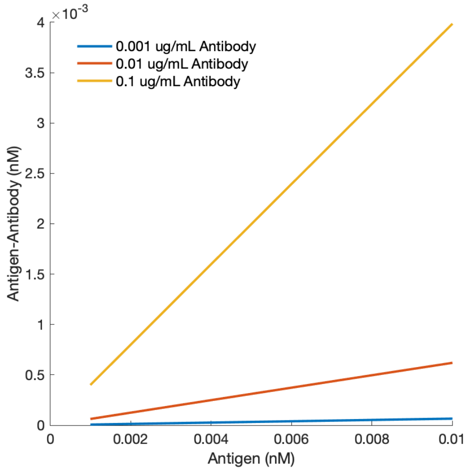


Modeling with $K_d$=1 nM, 1 pM ≤ $x$ ≤ 10 pM, $[B]_{tot}$= 0.001, 0.01, 0.1 μg/mL ≈ 7, 70, 700 pM.

The difference between two slopes can be due to an increase in $[B]_{tot}$ or a decrease in $K_d$.

In the case of ACE2-HA and GFP-RBD binding (Figure 5), we consider A = ACE2-HA, B = GFP-RBD, and using the proportional behavior of DNA concentration to gene synthesis of CFE we tuned the conditions to be as in case d above, so that:

$$[AB]_{eq} \approx \frac{[B]_{tot}}{K_d + [B]_{tot}} \times [A]_{tot}$$

Tuning the DNA brush composition so that $[B]_{tot} \ll K_d$ during the experiment, we can further approximate

$$[AB]_{eq} \approx \frac{1}{K_d} \times [A]_{tot} \times [B]_{tot}$$

(All simulations done with MATLAB R2024a).

### Langmuir isotherm model

The binding of ACE2-HA to GFP-RBD or antigen-antibody binding can be modelled similarly to a Langmuir isotherm as follows.

The surface is covered with A-HA sites (A=ACE2) that bind a second molecule B (GFP-RBD), with adsorption rate $r_{ad} = k_{on} \times A \times B$ and desorption rate $r_d = k_{off} \times (AB)$.

At equilibrium, these rates are equal $r_{ad} = r_d$, so that:

$$\frac{k_{off}}{k_{on}} = K_d = \frac{[A]_{eq} \times [B]_{eq}}{[AB]_{eq}}$$

The total number of A-HA sites is:

$$A_{tot} = A_{eq} + [AB)]_{eq} = \frac{K_d \times [AB])_{eq}}{[B]_{eq}} + [AB)]_{eq}$$

The ratio of bound sites is

$$\theta = \frac{[AB]_{eq}}{[A]_{tot}} = \frac{[B]_{eq}}{K_d + [B]_{eq}}$$

At equilibrium $[B]_{eq} = [B]_{tot} - [AB]_{eq}$

If B is in excess so that $B_{eq} \approx B_{tot}$, then

$$\theta = \frac{B_{tot}}{K_d + B_{tot}}$$

#### Estimation of synthesized antigen

Our data shows a linear correlation between the total antigen concentration and the bound antibody concentration, which varies with total antibody concentration. This suggests we are in limit case *d*, where $[B]_{tot} \gg x$ and $x$ is in concentrations lower or similar to $K_d$ so that the reaction is not total. As the affinity of antibodies for their specific antigens is in the order of magnitude of $K_d \sim 1$ nM, the effective concentration of antigens captured on the surface is in the tens of nM or lower.

#### Fluorescence intensity quantification

Fluorescence intensity (FI) of the bound antibodies relates to the equilibrium antigen-antibody complex as follows:

$$FI = a[AB]_{eq} + b$$

In limit case *d*,

$$FI = a \times \frac{[B]_{tot}}{K_d + [B]_{tot}} \times [A]_{tot} + b$$

With the slope $m$

$$m = a \times \frac{[B]_{tot}}{K_d + [B]_{tot}}$$

#### Relative affinity calculation

We consider two antigens $A_1$ and $A_2$ that are synthesized in similar concentration ranges and bind to the same antibody $B$ with affinities $K_{d,1}$ and $K_{d,2}$ respectively. Both antigens are incubated with the same antibody in identical conditions. As derived above, in our experimental conditions, the antigen-antibody complex concentration $[AB]_{eq}$ depends linearly on the total antigen concentration $[A]_{tot}$ with the slope $m$ :

$$[AB]_{eq} \approx \frac{[B]_{tot}}{K_d + [B]_{tot}} \times [A]_{tot} = m \times [A]_{tot}$$

The ratio of the slopes of the antigens exposed to the same $[B]_{tot}$ is therefore:

$$\frac{m_1}{m_2} = \left(\frac{[B]_{tot}}{K_{d,1} + [B]_{tot}}\right) \times \left(\frac{K_{d,2} + [B]_{tot}}{[B]_{tot}}\right) = \frac{K_{d,2} + [B]_{tot}}{K_{d,1} + [B]_{tot}}$$

So that

$$K_{d,2} = \frac{m_1}{m_2} \times \left(K_{d,1} + [B]_{tot}\right) - [B]_{tot}$$

The affinities of RBD2 mutants to the CV30 antibody were calculated assuming $K_{d,1}$ = 8.4 nM[8] and $[B]_{tot}$ = 1 µg/mL = 6.7 nM assuming an antibody molecular weight of 150 kDa.

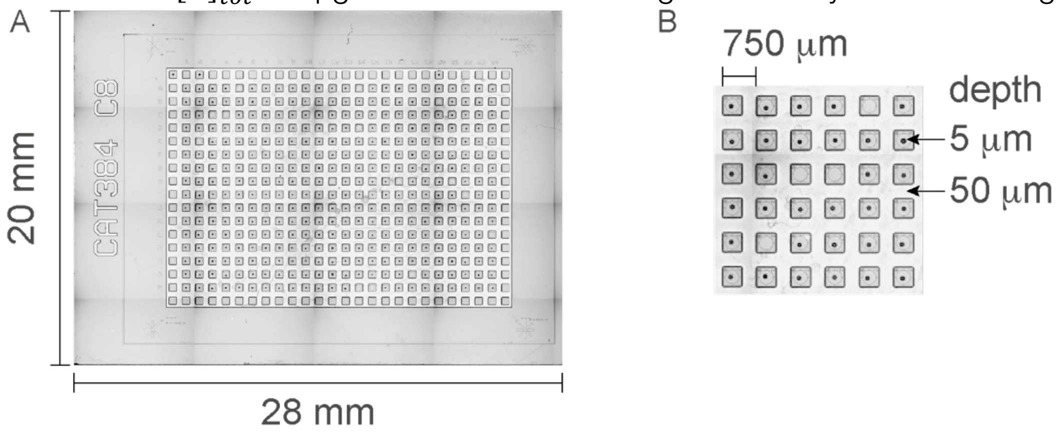


**Supplementary Figure 1** | Chip fabrication and dimensions. **A.** The whole 384-compartments chip is 20x28 mm, with 16 rows and 24 columns. **B.** Pedestals are 750 µm apart. The circular compartment carved in each pedestal to form the compartment is 5 µm deep. The channels in between the pedestals are 50 µm deep. Although a plasma-cleaned PDMS slab seals the chips and isolates each compartment, these deep channels act as a safety measure to guarantee that a gene overexpressing and overflowing from one compartment is diluted and does not contaminate a neighbor compartment.

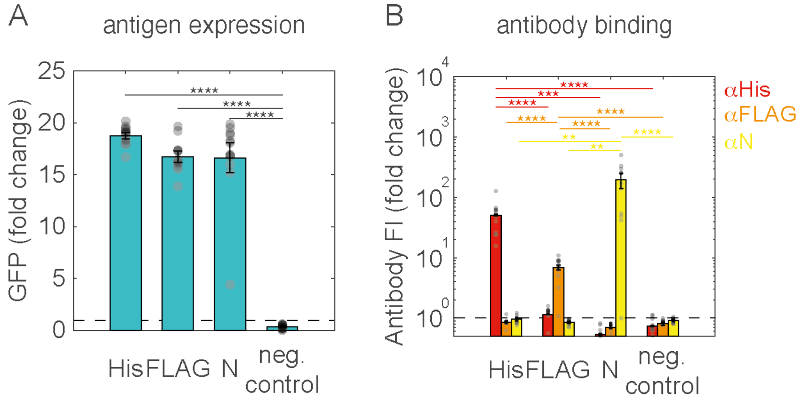


**Supplementary Figure 2** | Orthogonality of antigen recognition. **A.** Synthesis of antigens. Bars and error bars represent mean and s.e.m. of 10 compartments, dots represent individual data points. Black line represents a threshold of synthesis defined by three standard deviations above the average of the negative control. Stars represent *p*-values from a two-tailed Student's t-test: ** *p*<0.01, *** *p*<0.001, **** *p*<0.0001. **B**. Binding of antibodies (left: anti-His, center: anti-FLAG, right: anti-N). Antibodies are either directly labeled (His, FLAG) or a labeled secondary antibody is applied (N). Bars, error bars, dots and stars as in A. Black line represents a threshold of synthesis defined by one standard deviation above the average of the negative control.

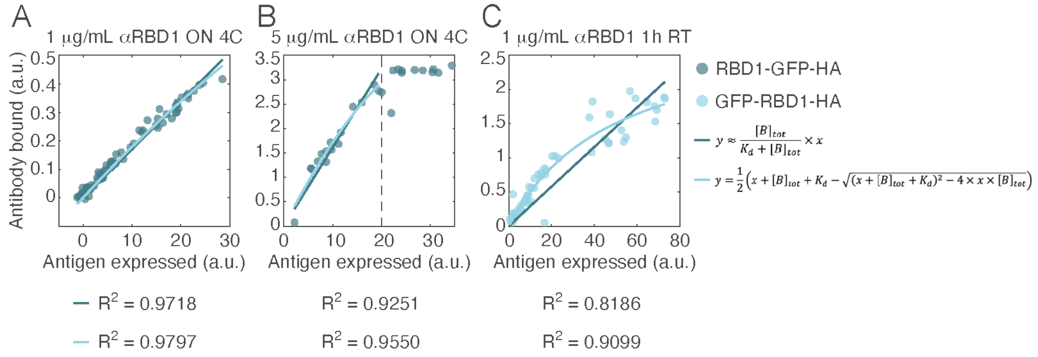


**Supplementary Figure 3** | Antibody-antigen bi-molecular interaction and fitting. 3 experiments demonstrated the effect of varying antibody concentration and incubation time on antibody-antigen binding. Each panel displays bound antibody fluorescence against synthesized antigen fluorescence for at least 55 compartments expressing the same specie (circles). This data is fitted with the simplified binding equation (dark blue) or the full equation (light blue) as described above. The quality of the fit is indicated by the $R^2$ parameter. **A.** 1 μg/mL anti-SARS-CoV-1 RBD (RBD1) mAb is incubated overnight at 4 °C on a chip having synthesized RBD1-GFP-HA antigen. The antibody-antigen data fits well with a linear approximation of the binding curve. **B.** An excess 5 μg/mL anti-RBD1 mAb is incubated overnight at 4 °C on a chip having synthesized RBD1-GFP-HA antigen. Above a certain level of antigen synthesis (20 arbitrary units a.u.), saturation of antibody binding is reached and the bound antibodies level plateau. Below that saturation, a linear fit is still a good approximation. **C.** 1 μg/mL anti-RBD1 mAb is incubated for a limited time (1h at room temperature RT) on a chip having synthesized GFP-RBD1-HA antigen. The antibody-antigen data no longer correctly fitted by a linear approximation of the binding equation. Instead, the full binding equilibrium equation describes better the bi-molecular interaction.

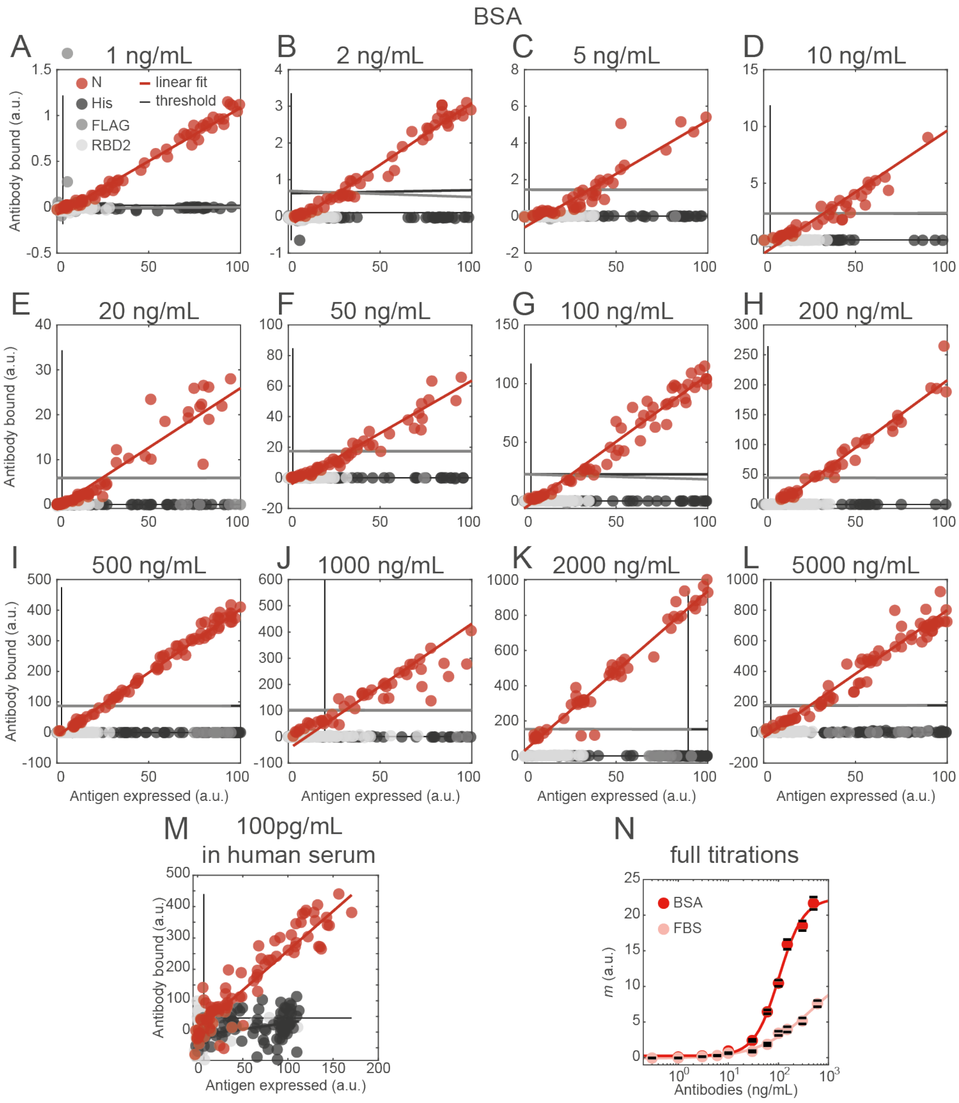


**Supplementary Figure 4** | Titration of anti-N mAb 1A6 binding to surface-displayed antigens. **A-M**. The bound antibody fluorescence is displayed against synthesized antigen fluorescence for at least 84 compartments expressing the same specie (circles). N antigens are displayed as red circles. Control antigens are: His (dark gray circles), FLAG (median gray circles), GFP-RBD2-HA (light gray circles). Linear fits are displayed as a solid line of the corresponding colour. The vertical and horizontal gray lines represent the thresholds for detection of an antigen (vertical) or antibody (horizontal) fluorescence signal. These thresholds are defined by the average of a negative control plus 3 standard deviations (for the antigen threshold, control is NC-HA), and as the average of a negative control plus 1 standard deviation (for the antibody threshold, control is His-GFP-HA). **A-L**. 12 chips spotted and synthesized identically are incubated with 12 different concentrations of anti-N mAb diluted in BSA PBS-T buffer: **A**, 1 ng/mL, **B**, 2 ng/mL, **C**, 5 ng/mL, **D**, 10 ng/mL, **E**, 20 ng/mL, **F**, 50 ng/mL, **G**, 100 ng/mL, **H**, 200 ng/mL, **I**, 500 ng/mL, **J**, 1000 ng/mL, **K**, 2000 ng/mL, **L**, 5000 ng/mL. **M**. A chip is incubated with 100 pg/mL anti-N mAb diluted in human serum. **N**. Titration of anti-N mAb diluted in 2 different media are compared: dark red, BSA PBS-T buffer, light red, Fetal Bovine Serum. At least 250 compartments synthesizing N antigens are fitted with a linear fit for each antibody concentration. Circles and error bars represent the fitted slope value and a 95% confidence interval of the fit. The thick line represents a fit with a Hill function of cooperativity 1.

N and Chimeric anti-N 1A6 Ab

A

**On-chip affinity measurement**

| | 2-steps fit | | | | 1-step fit | | | |
|---|---|---|---|---|---|---|---|---|
| | $K_d$ fit mean (nM) | $K_d$ fit stdev (nM) | $K_d$ mean (nM) | $K_d$ stdev (nM) | $K_d$ fit mean (nM) | $K_d$ fit stdev (nM) | $K_d$ mean (nM) | $K_d$ stdev (nM) |
| BSA | 4.1 | 0.5 | 3.64 | 1.32 | 3.62 | 0.19 | 3.72 | 1.31 |
| | 1.15 | 0.02 | | | 1.51 | 0.03 | | |
| | 5.65 | 0.01 | | | 6.0 | 0.2 | | |
| FBS | 1.51 | 0.01 | 2.64 | 1.05 | 0.87 | 0.01 | 1.14 | 0.36 |
| | 4.7 | 0.4 | | | 1.84 | 0.01 | | |
| | 1.66 | 0.01 | | | 0.70 | 0.04 | | |

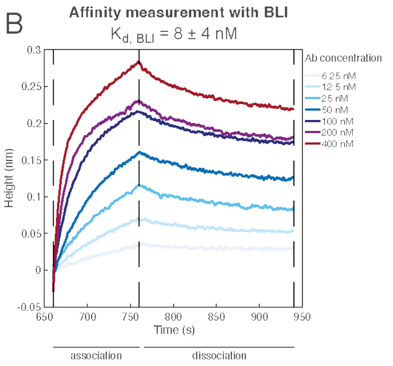


RBD WT and Rabbit anti-RBD 5G8 Ab

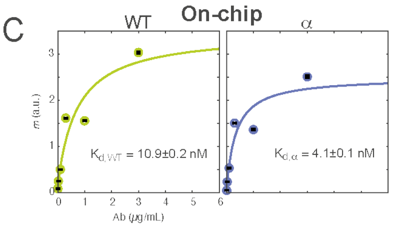


D

**BLI**

$K_{d,\ WT}$ = 5 ± 3 nM

$K_{d,\ \alpha}$ = 1 ± 2 nM

**Supplementary Figure 5** | Affinity measurement through antibody titration. **A,B.** Affinity measurement for N antigen/anti-N mAb 1A6. **A.** On-chip affinity measurements with 1A6 antibody diluted in BSA PBS-T buffer or FBS. The average and standard deviation of the fitted $K_d$ value is indicated for 3 different titration experiments, and then averaged over these 3 biological repeats. 2-steps fit: for each total antibody concentration, the slope $m$ of antibody-antigen binding is fitted with $FI_{Ab,bound} = FI_{Ag,total} \times m$. The values of $m$ are then fitted against antibody concentration with $m = \frac{[Ab]_{total}}{K_d + [Ab]_{total}} \times C$, with $K_d$ and $C$ as fit parameters. 1-step fit: all data for all antibody concentrations are simultaneously fitted with $FI_{Ab,bound} = FI_{Ag,total} \times \frac{[Ab]_{total}}{K_d + [Ab]_{total}} \times C$, with $[Ab]_{total}$ fixed for each antibody concentration condition, and $K_d$ and $C$ as fit parameters. Both fitting methods lead consistent values of $K_d$. **B.** Biolayer interferometry (BLI) affinity measurement. CFE synthesized N-GFP-HA is bound to the streptavidin-coated biolayer via biotinylated anti-HA antibodies. Titrated concentrations of 1A6 are then associated to N and dissociated in buffer. Association constant $k_{on}$ and dissociation constant $k_{off}$ are fitted for each $[Ab]_{total}$, $K_d$ is then calculated for each condition and averaged over all $[Ab]_{total}$. **C,D.** Affinity measurement for RBD2 WT or Alpha antigens/Rabbit anti-RBD2 mAb 5G8. **C.** Fitted $K_d$ values. Bars and error bars represent fitted $K_d$ value and 95% confidence interval of the fit, line represents fit of all slopes at once. **D.** BLI measured values of affinity for WT variant and α variant.

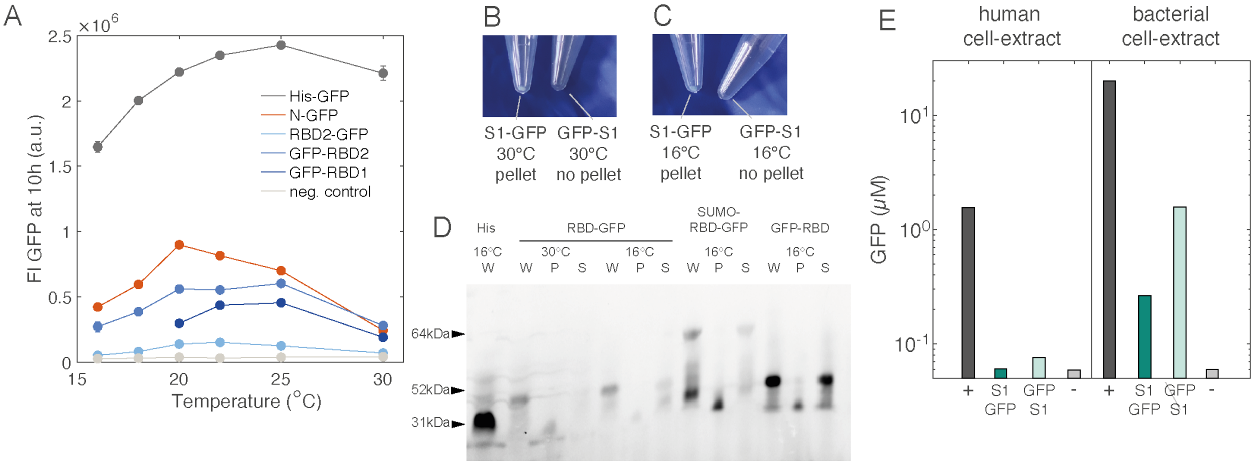


**Supplementary Figure 6** | Synthesis and solubility of Spike1 (S1) and RBD2 in bacterial CFE and insufficient synthesis in human CFE. **A.** GFP levels at 10 h of CFE of various constructs (colors) at a range of temperatures. Bars and error bars represent mean and standard deviation of three technical repeats. Negative control is synthesis of ACE2-HA (light grey line). **B-C.** Solubility of Spike protein synthesized by CFE. Photos show 20 µl of CFE reaction after 12 hours expression at the indicated temperature of a linear Streptavidin conjugated DNA fragment coding for the protein mentioned in the corresponding images. Aggregation of proteins is visualized by the formation of a pellet after centrifugation. **B.** Expression of constructs at 30 °C, followed by spinning and removing of supernatant. The pellet is green fluorescent because constructs are GFP labeled. Left, S1-GFP, visible pellet. Right, GFP-S1, no visible pellet. **C.** Expression of constructs at 16 °C, followed by spinning and removing of supernatant. Left, S1-GFP, small pellet. Right, GFP-S1, no pellet. **D.** SDS PAGE analysis of cell-free synthesized His (at 30 ºC), RBD2-GFP (at 30 ºC and 16 ºC) and GFP-RBD2 (at 16 ºC). Molecular weights of proteins indicated with arrow: His-GFP-HA 31kDa and RBD2-GFP-HA/GFP-RBD2-HA 52 kDa. W: whole CFE reaction after expression. S: soluble fraction, supernatant after centrifugation of the CFE reaction. **E.** HeLa cell-extract (left panel) is used to synthesize a positive control (dark gray, pCFE-GFP), S1-GFP-HA (dark blue) or GFP-S1-HA (light blue). *E. coli* bacterial CFE reaction (right panel) is used to synthesize a positive control (dark gray, His-GFP-HA), S1-GFP-HA (dark blue) or GFP-S1-HA (light blue). Negative controls are indicated in light gray. All DNA constructs are optimized for expression in human or bacterial cells respectively. End point (t = 10h) GFP fluorescence measurements are shown as bars. A positive control in human cell-extract synthesizes ~1 µM GFP compared to ~10 µM GFP in bacterial CFE reaction, demonstrating the higher total expression capacities in bacterial CFE reaction. S1-GFP-HA synthesizes in >100 nM equivalent units of GFP and GFP-S1-HA in > 1 µM in bacterial CFE reaction. In comparison, both S1-GFP-HA and GFP-S1-HA were synthesized below 100 nM equivalent units of GFP in human cell-extract.

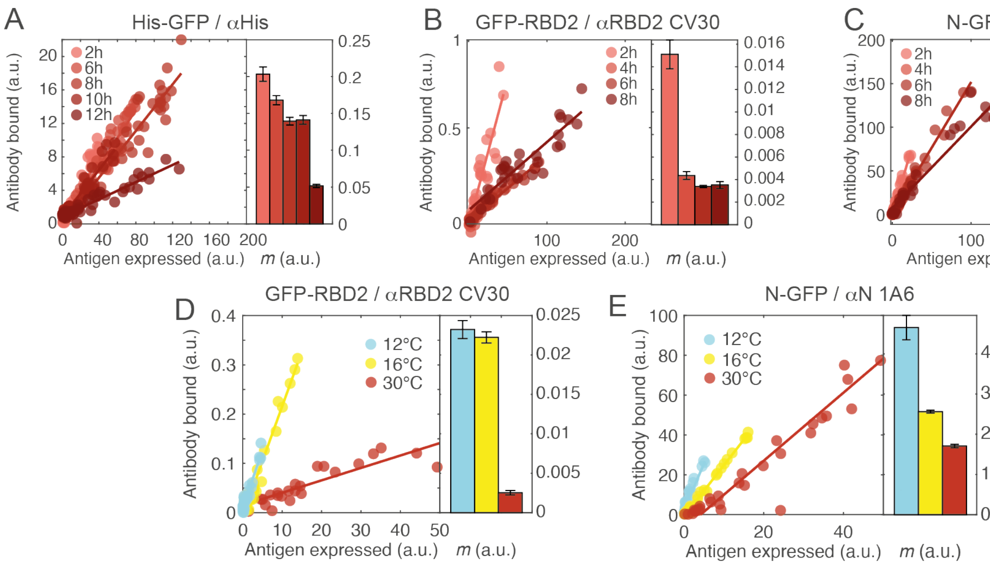


**Supplementary Figure 7** | Time and temperature effect on on-chip antigen synthesis and recognition by antibodies. **A-C**. Effect of time of synthesis on the concentration-dependent linear binding of antibodies to antigens. As the same antibody solution is used for all chips, the change in the slope reflects a change in the quality of the antigens displayed on the surface. Each left panel displays bound antibody fluorescence against synthesized antigen fluorescence for at least 36 compartments expressing the same specie (circles). The specie and antibody are indicated in the figure legend. The data is fitted with a linear fit, and the right panel displays the fitted slope value and a 95% confidence interval of the fit as bars and error bars. Time of expression (from 2 h to 8-12 h) is represented by shades of red from lighter (2h) to darker (8 h to 12 h). For several antigens, the data shows that the slope of binding gets lower for higher expression time, suggesting less well-recognized antigens displayed on the surface. **A**. His-GFP-HA. **B**. GFP-RBD2-HA. **C**. N-GFP-HA. **D-E**. Effect of temperature of expression on slope of antibody-antigen binding. Left panel displays bound antibody fluorescence against synthesized antigen fluorescence for at least 40 compartments synthesizing the Ag (circles). The chip is stained with CV30 (**D**) or 1A6 (**E**). The data is fitted with a linear fit, and the right panel displays the fitted slope value and a 95% confidence interval of the fit as bars and error bars. Temperature of expression is represented by light blue (12 °C), yellow (16 °C) and red (30 °C). **D**. GFP-RBD2-HA. **E**. N-GFP-HA.

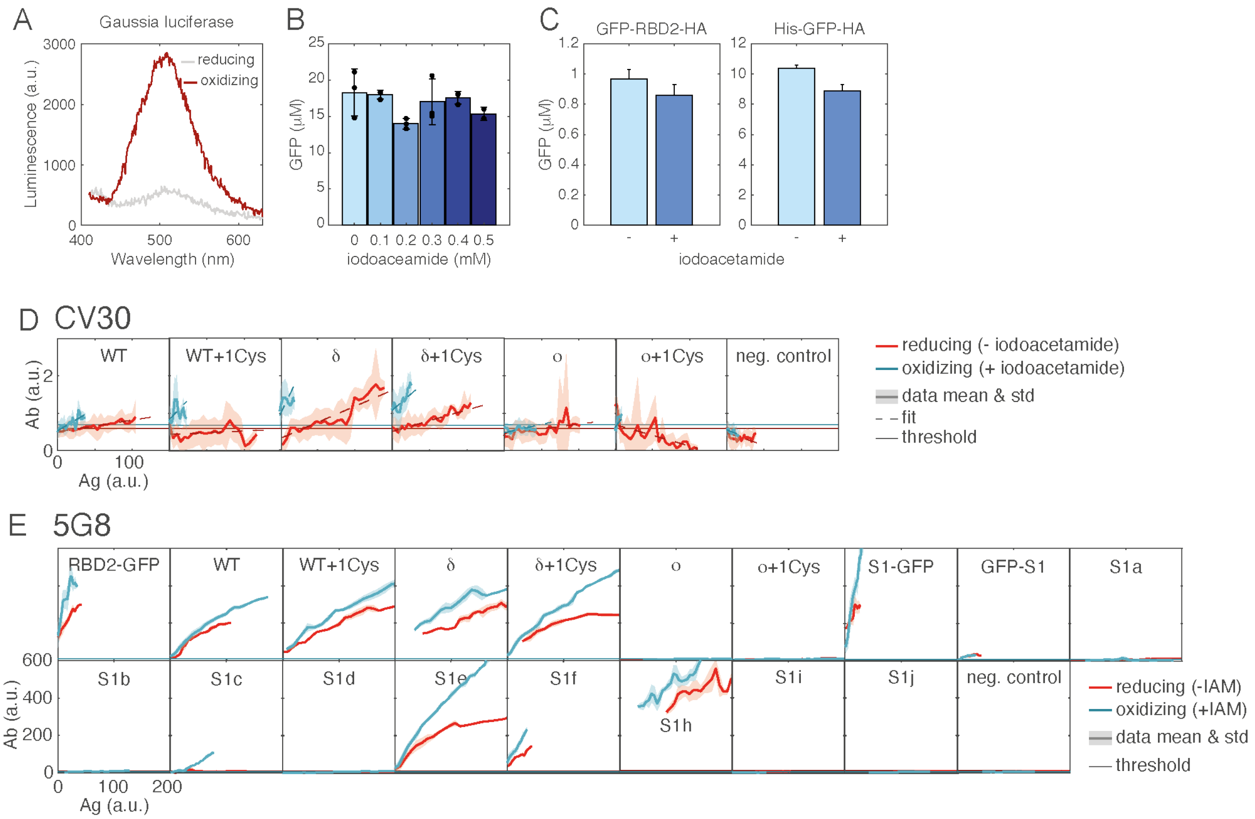


**Supplementary Figure 8** | Disulfide bond formation in oxidizing conditions does not improve antibody recognition. **A.** Luminescence of Gaussia luciferase synthesized in *E. coli* CFE reaction in native (reducing, grey) conditions or oxidizing conditions (red, with 0.5 mM iodoacetamide). Correct folding of Gaussia luciferase depends on the formation of disulfide bonds. Luminescence was measured from a 10 times diluted CFE reaction sample with the Pierce™ Gaussia Luciferase Glow Assay Kit (16160, Thermo Fisher Scientific, USA) in a Clariostar plate reader with a white 96-compartment plate. **B.** Synthesis of a His-GFP reporter is decreased (by ~15%) in presence of iodoacetamide ranging from 0.1 to 0.5 mM iodoacetamide. **C.** Synthesis of GFP-RBD2-HA is not improved in the presence of 0.5 mM iodoacetamide. It's decreased in the same proportions as a positive synthesis control His-GFP-HA. **D.** On chip binding of anti-RBD2 mAb CV30 to antigen in reducing and oxidizing environments. +1Cys: GFP-RBD2-HA construct cloned with 4 additional amino acids (CGPK), adding an additional cysteine to the sequence (from 7 to 8), and permitting an additional disulfide bond (see Supplementary Table 1). Red: native reducing environment of the CFE reaction. Blue: oxidizing environment (+ 0.4 mM IAM). Thick line: mean, transparent area: standard deviation, dotted line: fit, thin line: threshold defined by the average of negative control mean + 1 standard deviation. The CV30, has significant batch-to-batch variability and is unstable through long-term storage, giving a noisy signal in this specific experiment. **E.** On-chip binding of anti-RBD2 mAb 5G8 to antigen in reducing and oxidizing environment. Legend as in **D**.

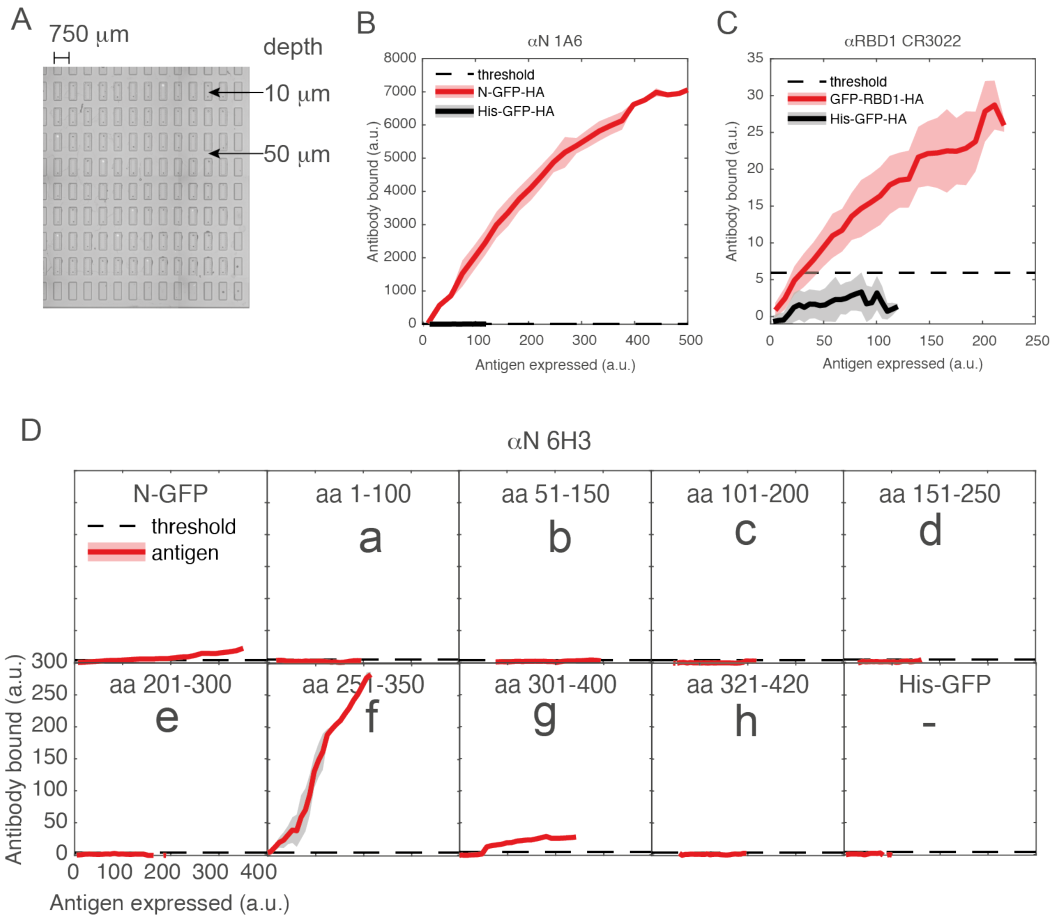


**Supplementary Figure 9** | Elongated compartments. A. Brightfield image of a 96 compartment-chip with dimensions. B-D. Binding of monoclonal antibodies to antigens in elongated compartments. Antigen brushes are spotted in at least 4 different compartments. Bound monoclonal antibody and antigen synthesized are measured with fluorescence. Each compartment consists of 320 individual data points along the long axis of the compartment. Antigen species are plotted in red and negative control (His-GFP-HA) in grey. Lines and shaded area correspond to mean and standard deviation. Dotted line is threshold, defined by the maximum average plus 1 standard deviation of the negative control. B. anti-N mAb 1A6 against N antigen. C. anti-RBD1 mAb CR3022 against RBD1 antigen. D. anti-N mAb 6H3 against N and fragments of N antigens. Results obtained from circular compartments with titrated antigen synthesis (Figure 3 and Supplementary Figure 11) are recapitulated here.

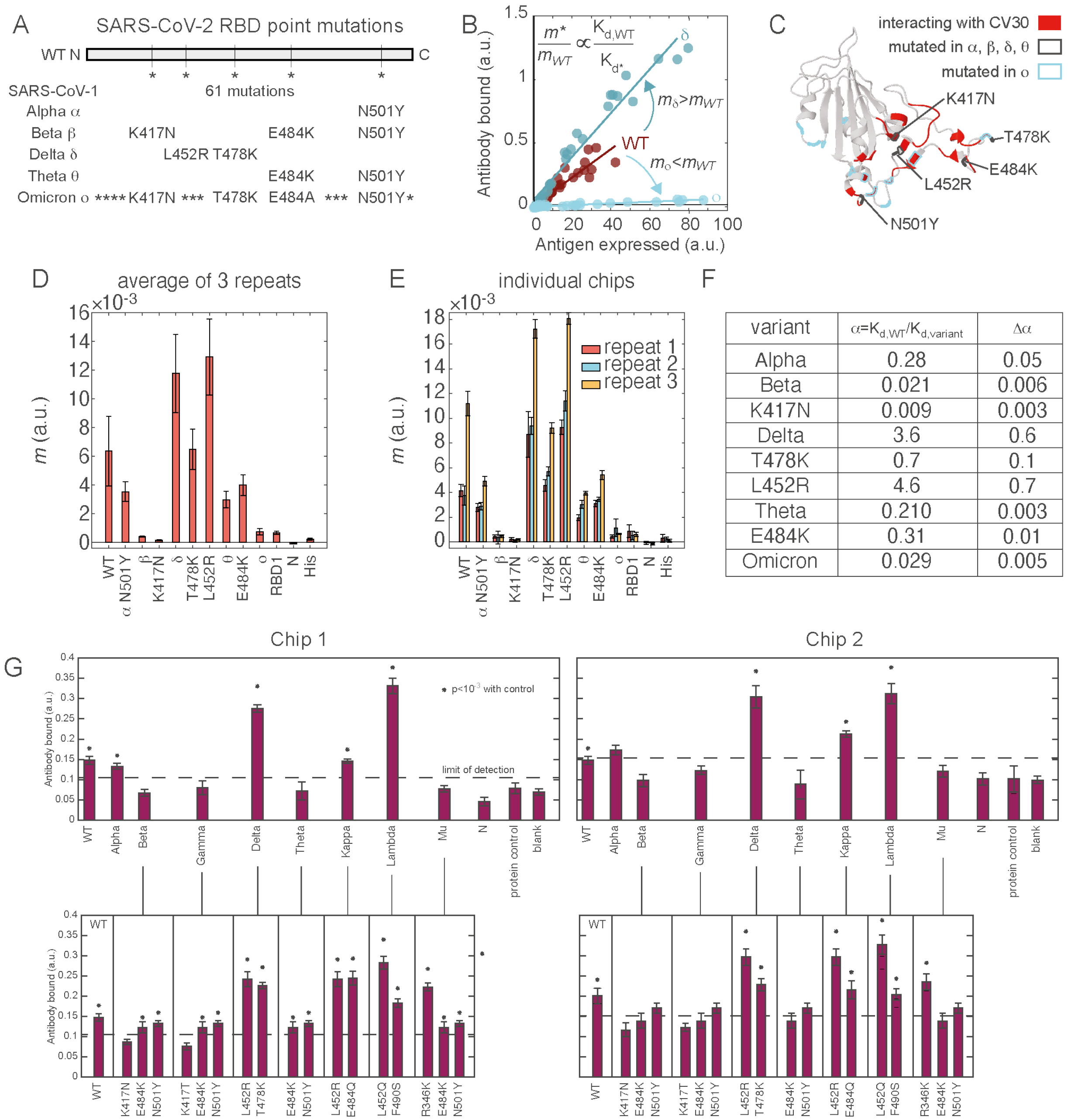


F

| variant | $\alpha=K_{d,WT}/K_{d,variant}$ | $\Delta\alpha$ |
|---|---|---|
| Alpha | 0.28 | 0.05 |
| Beta | 0.021 | 0.006 |
| K417N | 0.009 | 0.003 |
| Delta | 3.6 | 0.6 |
| T478K | 0.7 | 0.1 |
| L452R | 4.6 | 0.7 |
| Theta | 0.210 | 0.003 |
| E484K | 0.31 | 0.01 |
| Omicron | 0.029 | 0.005 |

**Supplementary Figure 10** | Epitope recognition of anti-RBD2 mAb CV30. **A-G.** Three different chips are incubated with CV30. At least 28 compartments synthesizing RBD2 and variants antigens are fitted with a linear fit. Three control antigens are also displayed on the chip: RBD1-GFP-HA, N-GFP-HA and His-GFP-HA. **A.** Sequence differences between SARS-CoV-2 RBD wild-type (RBD2 WT) variant, SARS-CoV-1 RBD (RBD1), and SARS-CoV-2 variants of concern. A star indicates a point mutation. **B.** CV30 mAb bound (measured by labeled secondary antibody fluorescence) against antigen synthesized (measured by GFP fluorescence) for three variants (dark blue: δ, light blue: ο) compared to WT (dark red). Dots represent individual compartment data and lines represent linear fit. **C.** 3D structure of RBD2 with mutated amino acids indicated in black (variants α, β, δ, θ) and blue (ο), and amino acids interacting with CV30 antibody in red, as characterized in Hurlburt *et al.*[9]. **D**. Bars and error bars represent the mean and standard deviation of the fitted slope values averaged between the 3 different chips. **E**. Bars and error bars represent the fitted slope value and a 95% confidence interval of the fit in each chip (red, blue, yellow). **F**. The relative affinity of each variant compared to WT is calculated by comparing the relative slopes as described above. The calculated relative affinity and its error based on the 95% confidence interval of the fit is shown for repeat 3. **G**. Two chips presenting RBD2 WT, 8 variants, 11-point mutants and 3 controls are assessed for antibody recognition of these antigens. Bars and error bars represent the mean and s.e.m. of 9 compartments spotted with the same brush. Here, antigen concentration is not titrated, all compartments are spotted with the same brush composition of different sequences. The limit of detection is taken as the protein control (His-GFP-HA) mean + 1 standard deviation. Stars represent p-values from a two-tailed Student's t-test: * $p<0.001$.

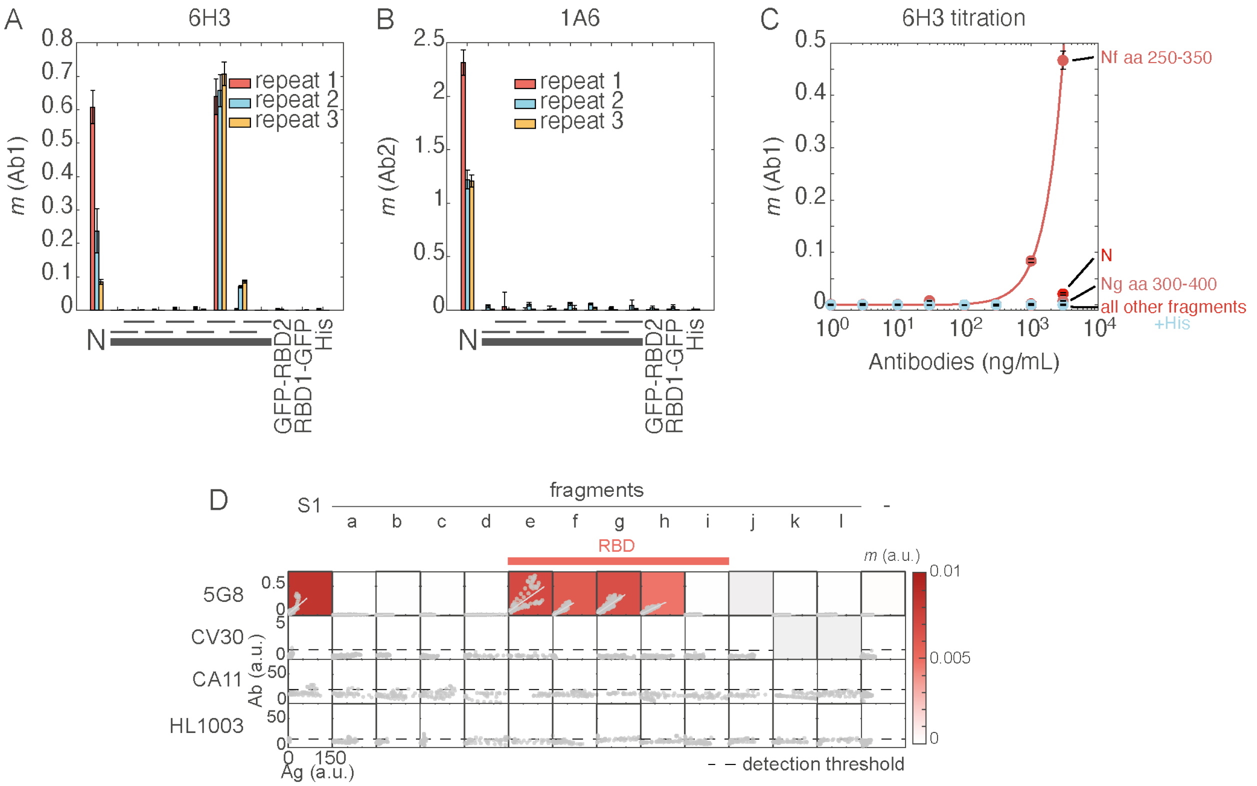


**Supplementary Figure 11** | Epitope recognition of two different anti-N antibodies. Three different chips are incubated with anti-N mAbs 6H3 (**A**) or 1A6 (**B**). At least 32 compartments synthesizing N and N-fragments antigens are fitted with a linear fit for each antibody. Bars and error bars represent the fitted slope value and a 95% confidence interval of the fit in each chip (red, blue, yellow). N-fragments are 100 amino acids (aa) long and cover the full sequence of N as follows. Three control antigens are also displayed: GFP-RBD2-HA, RBD1-GFP-HA and His-GFP-HA. **C**. Titration of 6H3 diluted in BSA PBS-T buffer. 20 compartments synthesizing each N fragment are fitted with a linear fit for each antibody concentration. Circles and error bars represent the fitted slope value and a 95% confidence interval of the fit. The thick line represents a two-step fit. Different antigen species are indicated in the right hand-side legend. **D**. Antibody binding for anti-RBD2 mAbs 5G8, CV30,

CA11 and HL1003 against antigens S1, fragments (a-l) and negative control (His-GFP-HA). Dots: data from 1 chip, lines: fitted slope averaged over all data points. Background color: slopes values, $m$, each slope fitted from at least 960 data points.

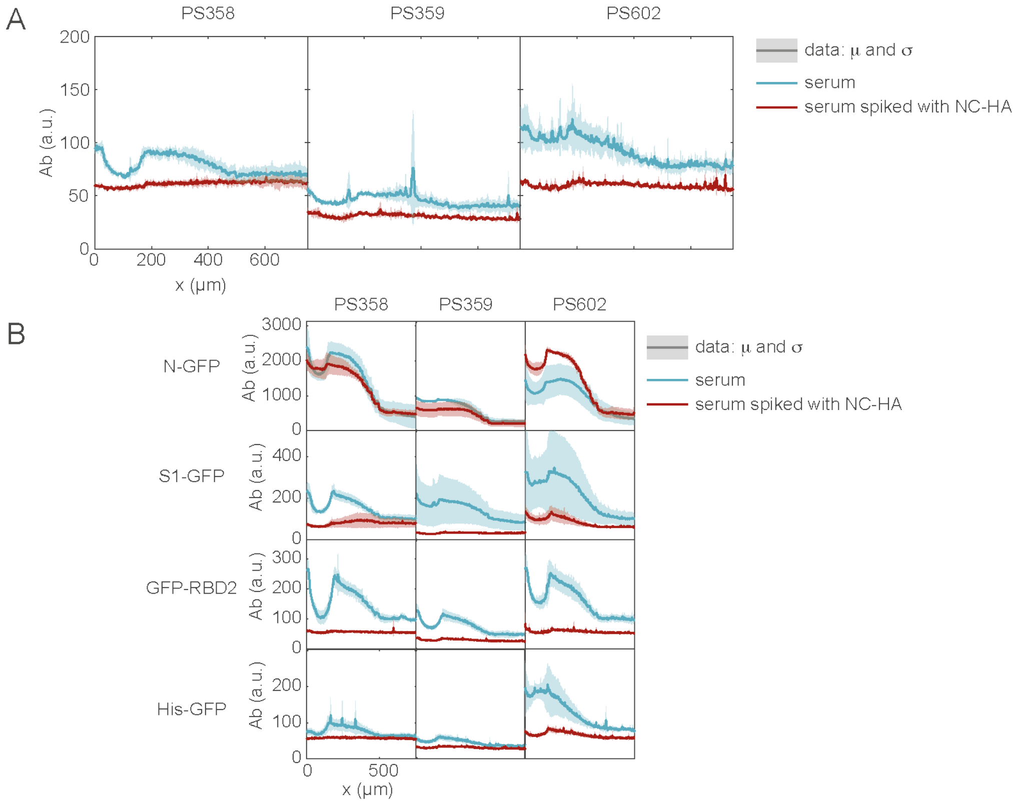


**Supplementary Figure 12** | Serum spiking and antibodies level. Sera were spiked with NC-HA synthesized in crude *E. coli* bacterial CFE to capture all anti-HA antibodies that may be present in human sera. Spiked sera (red) and negative control not spiked sera (blue) were incubated for 10 min with identical volumes of NC-HA and BSA PBS-T respectively (see Methods). All graphs show antibody binding along the gradient of the compartment. **A.** Secondary anti-Human IgG Ab binding to negative control NC-HA, indicating a non-specific adsorption of serum antibodies or binding of anti-HA antibodies. Sera samples (left: PS358, center: PS359, right: PS602) that were not spiked show a strong and location-specific adsorption of antibodies, which was minimized in the spiked samples. **B.** Antibody binding to various antigen targets in spiked and non-spiked serum samples (left: PS358, center: PS359, right: PS602). Spiking decreased antigen non-specific binding and allowed to detect true positive antigen hits. Top: N, center-top: S1-GFP, center-bottom: GFP-RBD2, bottom: negative control His.

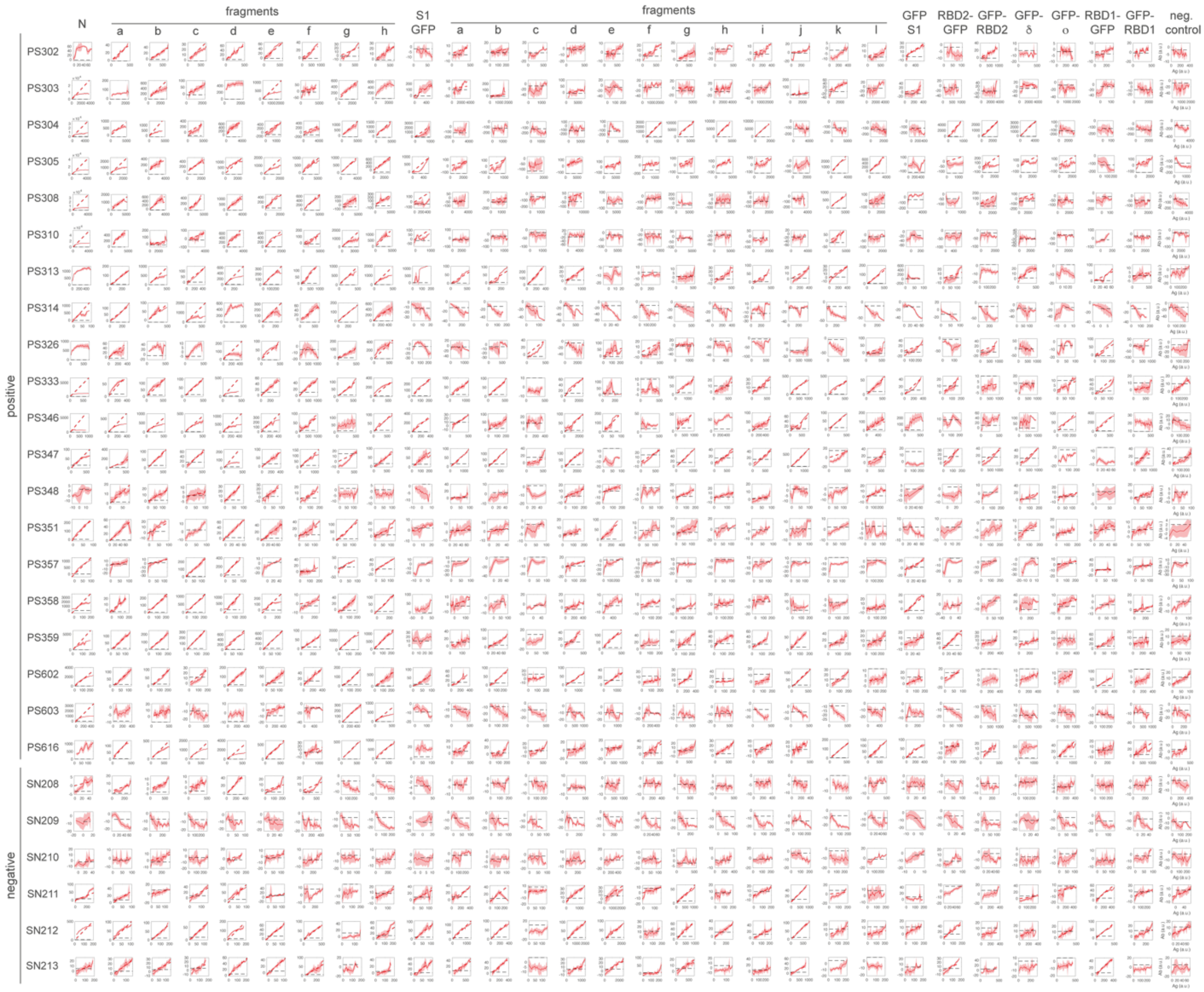


**Supplementary Figure 13** | All data from 20 positive and 6 negative human sera samples tested against 30 antigens. Binding curves of serum antibodies from a single human sample (each line) to antigens (columns). Thick line and shaded area represent the mean and standard deviation of 3 compartments. Dotted red line represents the fit. Dotted grey line represents the threshold for detection (maximum of negative control His average plus one standard deviation).

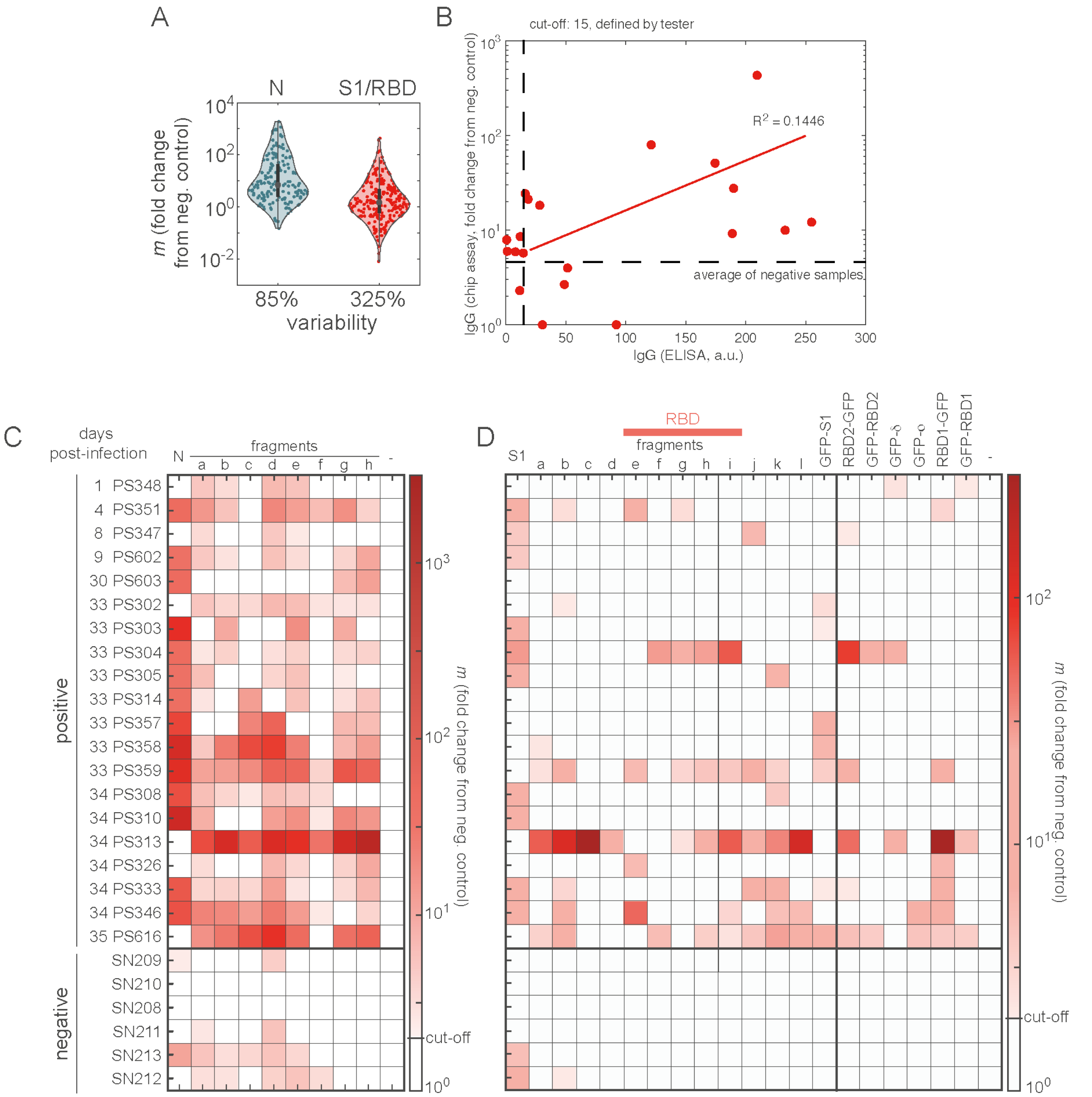


**Supplementary Figure 14** | Human sera response. **A**. Distribution of slopes across all positive samples and all antigens: left, N, right S1/RBD. Distributions envelopes are shown as violin plots, grey circle: median, dark thick line: first and third quartiles, dots: individual data points. Coefficient of variation is calculated as standard deviation over mean in percentage. **B**. On-chip IgG score (maximum IgG slope across all antigens, calculated as fold change from negative control) against ELISA IgG score for all 20 positive samples, indicated as red dots. Cut-off indicated as dashed black line: for ELISA, defined by the tester as 15 a.u., for on-chip IgG, defined as the average score of 6 negative samples. Linear fit indicated as line. Values for the two tests are uncorrelated ($R^2$ = 0.1446), but both tests show 14 (ELISA)/15 (on-chip) samples above threshold of detection. **C,D**. IgG scores for all 30 antigens sorted by days post-infection. Data represented as in Fig.5f,j.

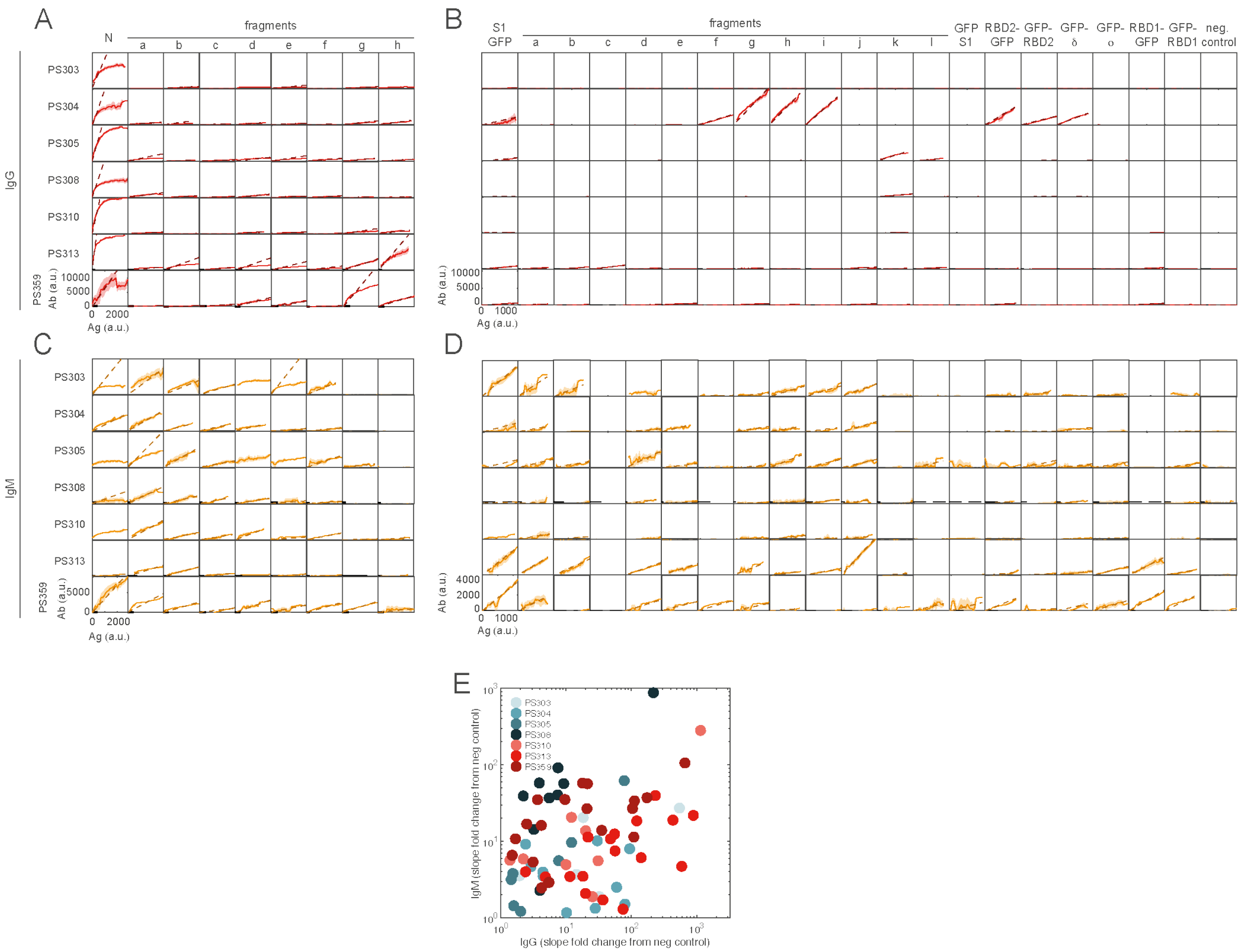


**Supplementary Figure 15** | IgG and IgM data from 7 positive human sera samples tested against 30 antigens. Binding curves of serum antibodies from a single human sample (each line) to antigens (columns). Thick line and shaded area represent the mean and standard deviation of 3 compartments. Dotted colored line represents the fit. Dotted grey line represents the threshold for detection (maximum of negative control His average plus one standard deviation). **A,B**. Red: IgG. **C,D**. orange: IgM. **A,C**. N and fragments. **B,D**. S1 and fragments, RBD and variants. **E**. Correlation between IgG and IgM for 7 samples tested: IgG and IgM scores (fold change of the slope above negative control) are plotted as colored dots for all antigens that had above background response in at least one antibody. No strong correlation within samples between antigens, and across all samples.

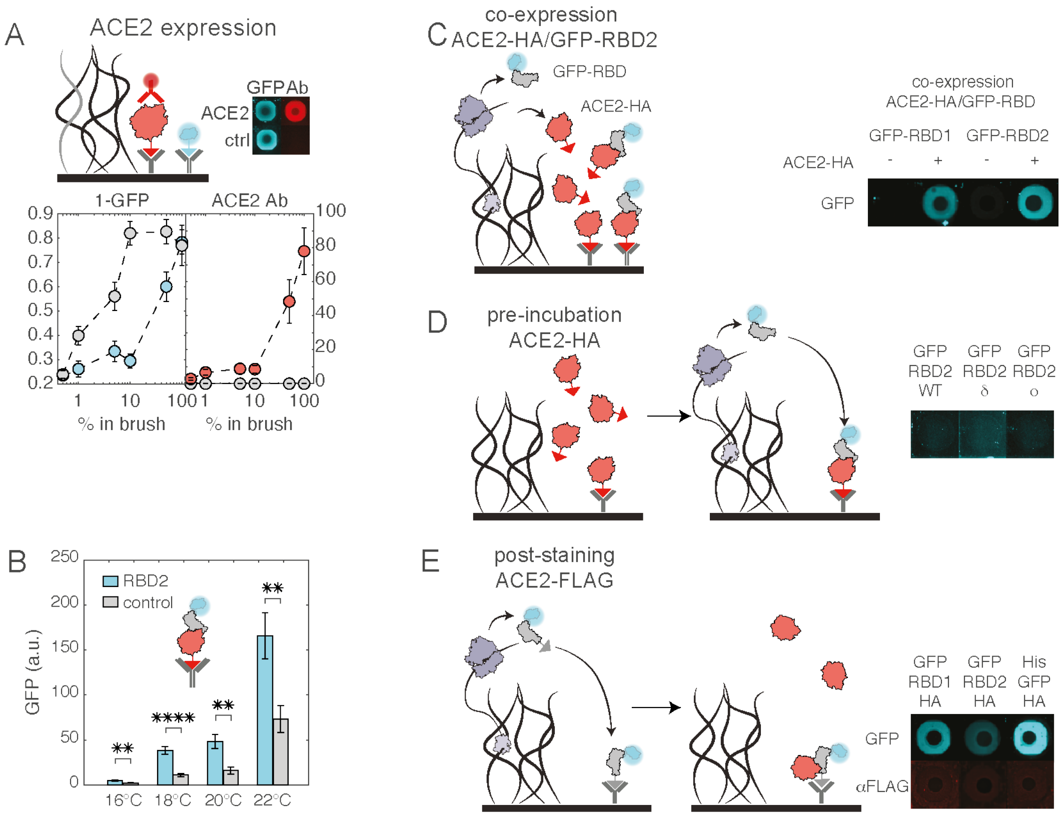


**Supplementary Figure 16** | Synthesis and binding of RBD2 and ACE2. **A**. On-chip synthesis of human receptor ACE2 and binding of monoclonal corresponding antibody. Different percentages of the coding gene in the brush (blue/red: ACE2-HA, grey: control NC-HA) are localized in different compartments. Synthesis (left panel) is indirectly measured by incubating the compartments with GFP-HA which covers any surface free from ACE2-HA or NC-HA (1-GFP, negative staining). Anti-ACE2 Ab and fluorescently labeled secondary antibody (right panel) shows ACE2 bound to the surface. Circles and bars represent mean and s.e.m. of 20 compartments. Microscopy images show single compartments GFP-HA negative staining (blue) and antibody (red). Compartment image width: 400 mm. **B**. Effect of temperature on assembly of ACE2-RBD complex. GFP fluorescence of compartments co-synthesis GFP-tagged SARS-CoV-2 RBD (RBD2, blue) and ACE2-HA are compared to compartments co-synthesizing a control protein (N-GFP-FLAG, control, grey) and ACE2-HA. GFP fluorescence is measured in chips synthesized at 4 different temperatures (as indicated in x-axis legend). Bars and error bars represent the mean and standard error of the mean of at least 12 compartments. Stars represent *p*-values from a two-tailed Student's t-test: ** $p<0.01$, **** $p<0.0001$ comparing the fluorescence of RBD2 compartments and control compartments. Significant and specific ACE2-RBD assembly could be observed in all 4 temperature conditions, and 18 °C was chosen for further experiments. **C-E**. ACE2-RBD interaction is only measured in co-expression conditions. ACE2-RBD interaction is tested in one-pot co-synthesis (**C**), by pre-incubation of pre-synthesized ACE2-HA before expression of GFP-RBD2 from the DNA brush (**D**), and by expression of GFP-RBD2-HA from the DNA brush followed by post-staining with pre-synthesized ACE2-FLAG (**E**). Left side, schemes of ACE2-RBD assembly in all 3 conditions. Right side, microscopy image of representative compartments. **C**. From left to right, synthesis of GFP-RBD1 alone, co-synthesis of GFP-RBD1 with ACE2-HA, synthesis of GFP-RBD2 alone, co-synthesis of GFP-RBD2 with ACE2-HA. **D**. From left to right, synthesis of GFP-RBD2-WT, Delta and Omicron on a surface pre-incubated with ACE2-HA. No significant fluorescent signal can be seen in the compartments, indicating lack of assembly of the protein-protein complex. **E**. From left to right, synthesis of GFP-RBD1-HA, GFP-RBD2-HA and a control His-GFP-HA. Top: synthesis measured by GFP fluorescence (blue), bottom: ACE2-FLAG binding measured by labeled anti-FLAG antibody fluorescence (red). Compartment image width: 400 μm.

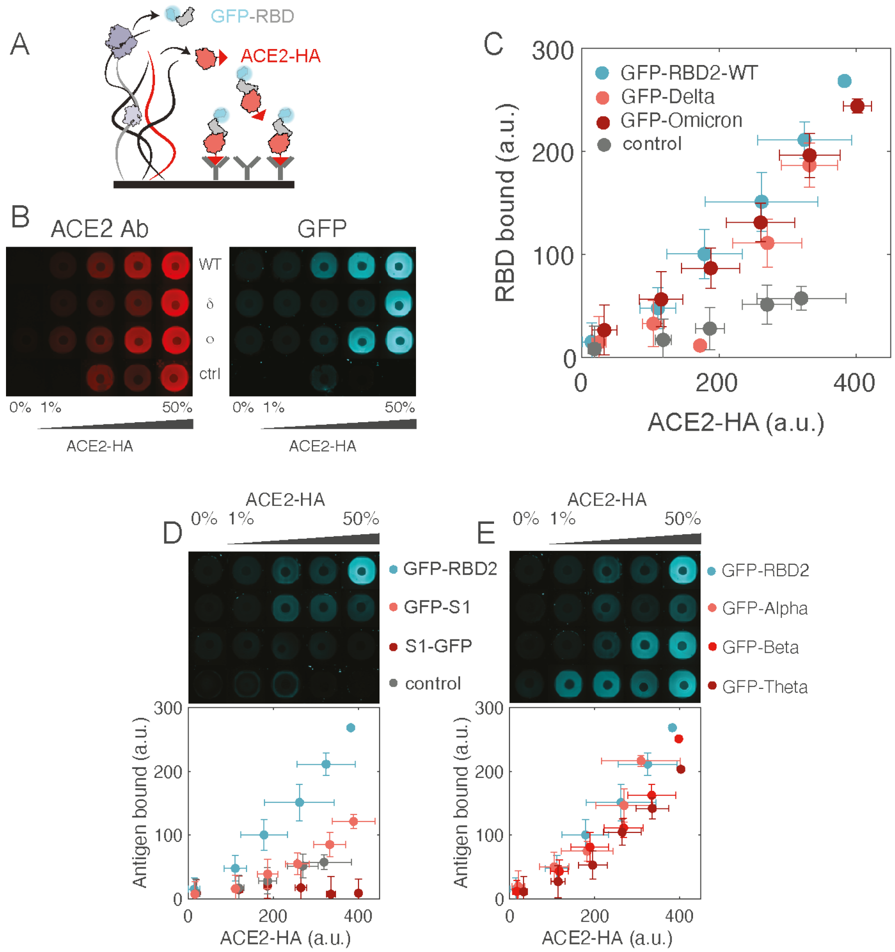


**Supplementary Figure 17** | Binding of ACE2-RBD complex. **A**. Binding of ACE2 and RBD during co-synthesis on-chip. **B**. Fluorescence microscopy images of compartments co-synthesizing GFP-RBD and ACE2-HA at different brush percentages. Surface bound ACE2-HA (stained with labeled antibody, red, left) interacts with GFP-RBD (blue, right image). RBD sequence is wild-type (WT) or variants: delta (δ) and omicron (o). Negative control is N-GFP-FLAG. Compartment image width: 400 μm. **C-E**. Binding of various RBD-related constructs to ACE2. Top panels: fluorescence microscopy images of compartments co-synthesizing an RBD-related or control construct and ACE2-HA at different brush percentages as indicated by the legend. Compartment image width: 400 μm. Bottom panels: fluorescence of GFP-RBD bound to surface-captured ACE2-HA as a function of ACE2-HA concentration (measured with a labeled antibody). 3 separate chips (60 compartments per RBD-related construct) are binned in 6 bins of ACE2-HA antibody fluorescence, dots and error bars represent mean and standard deviation. RBD-related construct is, from top row to bottom row of fluorescence images: **C**. GFP-RBD2-WT (blue), GFP-$\delta$ (light red), GFP-$o$ (dark red) and a control N-GFP-FLAG (grey). **D**. GFP-S1 (light red), S1-GFP (dark red) and a control N-GFP-FLAG (grey). **E**. GFP-RBD2-WT (blue), GFP-$\alpha$ (light red), GFP-$\beta$ (median red), GFP-$\theta$ (dark red). The assay does not detect differences in RBD variants binding to ACE2. The weaker binding of GFP-S1 and S1-GFP can be attributed to their weaker synthesis as shown in Supplementary Figure 6.

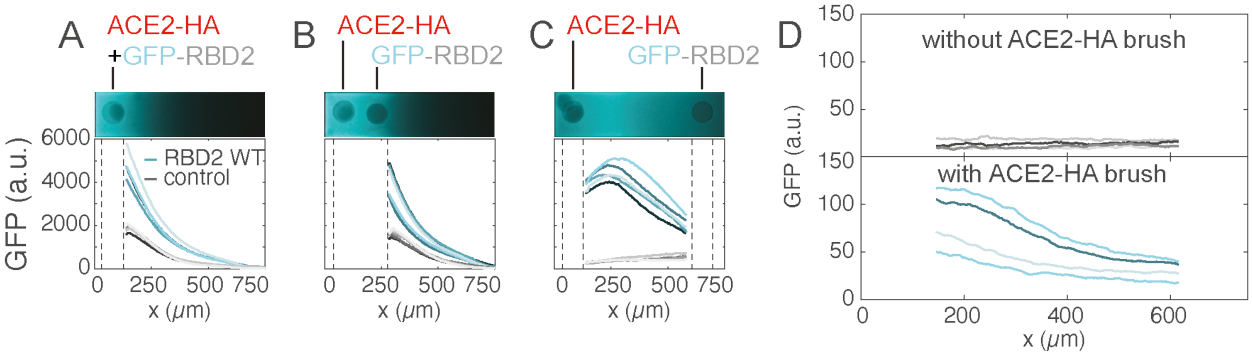


**Supplementary Figure 18** | Controls of ACE2-RBD2 protein-protein interaction. **A-C**. ACE2-RBD2 interaction with a single brush co-synthesizing both proteins (**A**) as in the circular compartment geometry, with two brushes close to one another (**B**) and two brushes far from each other (**C**). GFP-tagged protein is RBD2 (shades of blue) or a control protein not interacting with ACE2 (shades of grey). Microscopy images show one representative compartment for each condition. Graphs show GFP fluorescence after washing of at least 4 compartments. Above background specific protein-protein interaction is seen for all three conditions. The far apart brushes are the condition that minimizes non-specific interaction (low fluorescence in the control) and limits resource competition. **D**. GFP-RBD2 surface binding in the presence (bottom) or absence (top) of a ACE2-HA brush.

**Supplementary Table 1** | DNA sequences
In the following sequences, the different regions are marked as: T7 promoter, GFP, **protein of interest**, HA tag, T7 terminator
Notes: Links Sequences represented here are inserts, from T7 promoter to T7 terminator. Full plasmid maps (backbone pIVEX2.5) are linked. All sequences without HA simply have the HA tag sequence removed. The sequences of point mutants derive from their reference sequence as described in the main text. For the fragments of larger proteins, only DNA sequences of the fragments are indicated below.

His-GFP-HA
https://benchling.com/s/seq-revGTox6TSgrAPCEtszw?m=slm-N4MjAt28ffF70eVTcJFY
TAATACGACTCACTATAGGGAGACCACAACGGTTTCCCTCTAGAAATAATTTTGTTTAACT
TTAAGAAGGAGATATACC**ATGACCAGCCACCATCACCATCACCAT**ATCGAAGGCCGCG
GCCGCTTAATTAAACATATGACCATGGTGAGCAAGGGCGAGGAGCTGTTCACCGGGGT
GGTGCCCATCCTGGTCGAGCTGGACGGCGACGTAAACGGCCACAAGTTCAGCGTGTC
CGGCGAGGGCGAGGGCGATGCCACCTACGGCAAGCTGACCCTGAAGTTCATCTGCAC
CACCGGCAAGCTGCCCGTGCCCTGGCCCACCCTCGTGACCACCCTGACCTACGGCGT
GCAGTGCTTCAGCCGCTACCCCGACCACATGAAGCAGCACGACTTCTTCAAGTCCGCC
ATGCCCGAAGGCTACGTCCAGGAGCGCACCATCTTCTTCAAGGACGACGGCAACTACA
AGACCCGCGCCGAGGTGAAGTTCGAGGGCGACACCCTGGTGAACCGCATCGAGCTGA
AGGGCATCGACTTCAAGGAGGACGGCAACATCCTGGGGCACAAGCTGGAGTACAACTA
CAACAGCCACAACGTCTATATCATGGCCGACAAGCAGAAGAACGGCATCAAGGTGAAC
TTCAAGATCCGCCACAACATCGAGGACGGCAGCGTGCAGCTCGCCGACCACTACCAG
CAGAACACCCCCATCGGCGACGGCCCCGTGCTGCTGCCCGACAACCACTACCTGAGC
ACCCAGTCCGCCCTGAGCAAAGACCCCAACGAGAAGCGCGATCACATGGTCCTGCTG
GAGTTCGTGACCGCCGCCGGGATCACTCTCGGCATGGACGAGCTGTACAAGAGAAGA
GCAAAGAAGGAAGCCTACCCATACGATGTTCCAGATTACGCTTAATAAAAGGGCGAATT
CCAGCACACTGGCGGCCGTTACTAGTGGATCCGGCTGCTAACAAAGCCCGAAAGGAA
GCTGAGTTGGCTGCTGCCACCGCTGAGCAATAACTAGCATAACCCCTTGGGGCCTCTA
AACGGGTCTTGAGGGGTTTTTTG

FLAG-GFP-HA
https://benchling.com/s/seq-qRRpBhSGxXV6rdD0fNyw?m=slm-rOA54LyCMVLewuzoGWwN
TAATACGACTCACTATAGGGAGACCACAACGGTTTCCCTCTAGAAATAATTTTGTTTAACT
TTAAGAAGGAGATATACC**ATGACCAGCGACTACAAAGACGATGACGACAAG**ATCGAAG
GCCGCGGCCGCTTAATTAAACATATGACCATGGTGAGCAAGGGCGAGGAGCTGTTCAC
CGGGGTGGTGCCCATCCTGGTCGAGCTGGACGGCGACGTAAACGGCCACAAGTTCAG
CGTGTCCGGCGAGGGCGAGGGCGATGCCACCTACGGCAAGCTGACCCTGAAGTTCAT
CTGCACCACCGGCAAGCTGCCCGTGCCCTGGCCCACCCTCGTGACCACCCTGACCTA
CGGCGTGCAGTGCTTCAGCCGCTACCCCGACCACATGAAGCAGCACGACTTCTTCAAG
TCCGCCATGCCCGAAGGCTACGTCCAGGAGCGCACCATCTTCTTCAAGGACGACGGC
AACTACAAGACCCGCGCCGAGGTGAAGTTCGAGGGCGACACCCTGGTGAACCGCATC

GAGCTGAAGGGCATCGACTTCAAGGAGGACGGCAACATCCTGGGGCACAAGCTGGAG
TACAACTACAACAGCCACAACGTCTATATCATGGCCGACAAGCAGAAGAACGGCATCAA
GGTGAACTTCAAGATCCGCCACAACATCGAGGACGGCAGCGTGCAGCTCGCCGACCA
CTACCAGCAGAACACCCCCATCGGCGACGGCCCCGTGCTGCTGCCCGACAACCACTA
CCTGAGCACCCAGTCCGCCCTGAGCAAAGACCCCAACGAGAAGCGCGATCACATGGT
CCTGCTGGAGTTCGTGACCGCCGCCGGGATCACTCTCGGCATGGACGAGCTGTACAA
GAGAAGAGCAAAGAAGGAAGCCTACCCATACGATGTTCCAGATTACGCTTAATAAAAGG
GCGAATTCCAGCACACTGGCGGCCGTTACTAGTGGATCCGGCTGCTAACAAAGCCCGA
AAGGAAGCTGAGTTGGCTGCTGCCACCGCTGAGCAATAACTAGCATAACCCCTTGGGG
CCTCTAAACGGGTCTTGAGGGGTTTTTTG

N-GFP-HA
https://benchling.com/s/seq-IDTgRFpLakjMEHYZkLDP?m=slm-a0sm9YFjfFXrzvxBvcXp
TAATACGACTCACTATAGGGAGACCACAACGGTTTCCCTCTAGAAATAATTTTGTTTAACT
TTAAGAAGGAGATATACAT**ATGAGCGATAACGGACCCCAGAACCAGCGTAATGCCCC**
**TCGCATTACTTTCGGAGGACCCAGCGATTCTACCGGATCTAATCAAAACGGGGAGCG**
**CTCCGGGGCGCGTTCCAAACAACGCCGTCCGCAAGGGCTTCCCAACAACACTGCAT**
**CGTGGTTTACAGCATTGACGCAACATGGGAAAGAAGACTTGAAGTTCCCACGCGGC**
**CAGGGAGTGCCCATTAATACAAATTCTTCGCCCGACGACCAGATCGGATATTATCGT**
**CGCGCAACTCGCCGCATCCGCGGAGGTGATGGTAAAATGAAGGATTTAAGCCCGCG**
**TTGGTATTTCTACTACTTGGGAACGGGTCCCGAAGCGGGATTACCATATGGAGCAAA**
**CAAAGATGGGATTATCTGGGTTGCTACTGAGGGAGCGTTGAATACGCCGAAGGATCA**
**CATTGGTACGCGTAATCCAGCTAACAATGCTGCCATCGTTTTGCAACTTCCACAGGG**
**GACGACTCTGCCCAAGGGTTTCTACGCTGAAGGTTCGCGTGGGGGTCTCAGGCAT**
**CTTCCCGTAGTTCGTCTCGTTCGCGCAACTCATCCCGTAATTCCACACCCGGCTCCT**
**CACGCGGCACGAGTCCGGCACGCATGGCAGGGAATGGGGGAGATGCTGCCCTGG**
**CTTTATTACTGCTTGATCGTTTGAACCAATTGGAATCCAAGATGTCTGGAAAAGGACA**
**ACAGCAACAAGGTCAGACTGTGACTAAGAAATCAGCAGCAGAGGCGTCCAAGAAAC**
**CACGTCAAAAACGCACAGCTACGAAGGCGTATAACGTGACACAGGCATTTGGTCGTC**
**GCGGCCCGGAACAAACCCAAGGTAATTTCGGCGACCAAGAACTGATCCGTCAAGGG**
**ACGGACTATAAGCATTGGCCTCAGATTGCCCAGTTCGCTCCCTCGGCATCGGCATTC**
**TTCGGCATGTCACGCATCGGAATGGAAGTGACACCTAGTGGGACCTGGCTGACTTAT**
**ACAGGCGCAATCAAGCTTGATGACAAAGACCCTAACTTCAAGGATCAAGTAATTCTGT**
**TAAATAAGCACATTGATGCGTATAAGACATTCCCGCCGACCGAGCCAAAGAAAGATA**
**AGAAAAAGAAAGCCGACGAAACTCAAGCCTTGCCGCAGCGTCAAAAAAAGCAGCAG**
**ACTGTCACTCTTTTGCCAGCAGCAGATTTAGATGATTTTTCTAAACAACTTCAACAGTC**
**GATGTCCTCAGCGGACAGCACACAGGCC**AAGCGAGCTCCCGGGACCAGCATGGTGA
GCAAGGGCGAGGAGCTGTTCACCGGGGTGGTGCCCATCCTGGTCGAGCTGGACGGC
GACGTAAACGGCCACAAGTTCAGCGTGTCCGGCGAGGGCGAGGGCGATGCCACCTAC
GGCAAGCTGACCCTGAAGTTCATCTGCACCACCGGCAAGCTGCCCGTGCCCTGGCCC
ACCCTCGTGACCACCCTGACCTACGGCGTGCAGTGCTTCAGCCGCTACCCCGACCAC
ATGAAGCAGCACGACTTCTTCAAGTCCGCCATGCCCGAAGGCTACGTCCAGGAGCGCA
CCATCTTCTTCAAGGACGACGGCAACTACAAGACCCGCGCCGAGGTGAAGTTCGAGG
GCGACACCCTGGTGAACCGCATCGAGCTGAAGGGCATCGACTTCAAGGAGGACGGCA

ACATCCTGGGGCACAAGCTGGAGTACAACTACAACAGCCACAACGTCTATATCATGGC
CGACAAGCAGAAGAACGGCATCAAGGTGAACTTCAAGATCCGCCACAACATCGAGGAC
GGCAGCGTGCAGCTCGCCGACCACTACCAGCAGAACACCCCCATCGGCGACGGCCC
CGTGCTGCTGCCCGACAACCACTACCTGAGCACCCAGTCCGCCCTGAGCAAAGACCC
CAACGAGAAGCGCGATCACATGGTCCTGCTGGAGTTCGTGACCGCCGCCGGGATCAC
TCTCGGCATGGACGAGCTGTACAAGAGAAGAGCAAAGAAGGAAGccTACCCATACGAT
GTTCCAGATTACGCTTAATAAAAGGGCGAATTCCAGCACACTGGCGGCCGTTACTAGTG
GATCCGGCTGCTAACAAAGCCCGAAAGGAAGCTGAGTTGGCTGCTGCCACCGCTGAG
CAATAACTAGCATAACCCCTTGGGGCCTCTAAACGGGTCTTGAGGGGTTTTTTG

N fragments

a (aa 1-100):
AGCGATAACGGACCCCAGAACCAGCGTAATGCCCCTCGCATTACTTTCGGAGGACCCA
GCGATTCTACCGGATCTAATCAAAACGGGGAGCGCTCCGGGGCGCGTTCCAAACAAC
GCCGTCCGCAAGGGCTTCCCAACAACACTGCATCGTGGTTTACAGCATTGACGCAACA
TGGGAAAGAAGACTTGAAGTTCCCACGCGGCCAGGGAGTGCCCATTAATACAAATTCTT
CGCCCGACGACCAGATCGGATATTATCGTCGCGCAACTCGCCGCATCCGCGGAGGTG
ATGGTAAAATG

b (aa 51-150):
TGGTTTACAGCATTGACGCAACATGGGAAAGAAGACTTGAAGTTCCCACGCGGCCAGG
GAGTGCCCATTAATACAAATTCTTCGCCCGACGACCAGATCGGATATTATCGTCGCGCA
ACTCGCCGCATCCGCGGAGGTGATGGTAAAATGAAGGATTTAAGCCCGCGTTGGTATTT
CTACTACTTGGGAACGGGTCCCGAAGCGGGATTACCATATGGAGCAAACAAAGATGGG
ATTATCTGGGTTGCTACTGAGGGAGCGTTGAATACGCCGAAGGATCACATTGGTACGCG
TAATCCA

c (aa 101-200):
AAGGATTTAAGCCCGCGTTGGTATTTCTACTACTTGGGAACGGGTCCCGAAGCGGGATT
ACCATATGGAGCAAACAAAGATGGGATTATCTGGGTTGCTACTGAGGGAGCGTTGAATA
CGCCGAAGGATCACATTGGTACGCGTAATCCAGCTAACAATGCTGCCATCGTTTTGCAA
CTTCCACAGGGGACGACTCTGCCCAAGGGTTTCTACGCTGAAGGTTCGCGTGGGGGG
TCTCAGGCATCTTCCCGTAGTTCGTCTCGTTCGCGCAACTCATCCCGTAATTCCACACC
CGGCTCC

d (aa 151-250):
GCTAACAATGCTGCCATCGTTTTGCAACTTCCACAGGGGACGACTCTGCCCAAGGGTTT
CTACGCTGAAGGTTCGCGTGGGGGGTCTCAGGCATCTTCCCGTAGTTCGTCTCGTTCG
CGCAACTCATCCCGTAATTCCACACCCGGCTCCTCACGCGGCACGAGTCCGGCACGC
ATGGCAGGGAATGGGGGAGATGCTGCCCTGGCTTTATTACTGCTTGATCGTTTGAACCA
ATTGGAATCCAAGATGTCTGGAAAAGGACAACAGCAACAAGGTCAGACTGTGACTAAGA
AATCAGCA

e (aa 201-300):
TCACGCGGCACGAGTCCGGCACGCATGGCAGGGAATGGGGGAGATGCTGCCCTGGC
TTTATTACTGCTTGATCGTTTGAACCAATTGGAATCCAAGATGTCTGGAAAAGGACAACA
GCAACAAGGTCAGACTGTGACTAAGAAATCAGCAGCAGAGGCGTCCAAGAAACCACGT
CAAAAACGCACAGCTACGAAGGCGTATAACGTGACACAGGCATTTGGTCGTCGCGGCC
CGGAACAAACCCAAGGTAATTTCGGCGACCAAGAACTGATCCGTCAAGGGACGGACTA
TAAGCATTGG

f (aa 251-350):
GCAGAGGCGTCCAAGAAACCACGTCAAAAACGCACAGCTACGAAGGCGTATAACGTGA
CACAGGCATTTGGTCGTCGCGGCCCGGAACAAACCCAAGGTAATTTCGGCGACCAAGA
ACTGATCCGTCAAGGGACGGACTATAAGCATTGGCCTCAGATTGCCCAGTTCGCTCCC
TCGGCATCGGCATTCTTCGGCATGTCACGCATCGGAATGGAAGTGACACCTAGTGGGA
CCTGGCTGACTTATACAGGCGCAATCAAGCTTGATGACAAAGACCCTAACTTCAAGGAT
CAAGTAATT

g (aa 301-400):
CCTCAGATTGCCCAGTTCGCTCCCTCGGCATCGGCATTCTTCGGCATGTCACGCATCG
GAATGGAAGTGACACCTAGTGGGACCTGGCTGACTTATACAGGCGCAATCAAGCTTGAT
GACAAAGACCCTAACTTCAAGGATCAAGTAATTCTGTTAAATAAGCACATTGATGCGTAT
AAGACATTCCCGCCGACCGAGCCAAAGAAAGATAAGAAAAAGAAAGCCGACGAAACTC
AAGCCTTGCCGCAGCGTCAAAAAAAGCAGCAGACTGTCACTCTTTTGCCAGCAGCAGA
TTTAGAT

h (aa 319-418):
ATCGGAATGGAAGTGACACCTAGTGGGACCTGGCTGACTTATACAGGCGCAATCAAGC
TTGATGACAAAGACCCTAACTTCAAGGATCAAGTAATTCTGTTAAATAAGCACATTGATG
CGTATAAGACATTCCCGCCGACCGAGCCAAAGAAAGATAAGAAAAAGAAAGCCGACGA
AACTCAAGCCTTGCCGCAGCGTCAAAAAAAGCAGCAGACTGTCACTCTTTTGCCAGCA
GCAGATTTAGATGATTTTTCTAAACAACTTCAACAGTCGATGTCCTCAGCGGACAGCACA
CAGGCC

N-GFP-FLAG
https://benchling.com/s/seq-swuTnQAuUYxI0qrOHRQY?m=slm-eGQVTLOgqc7tZPMSvFbt
TAATACGACTCACTATAGGGAGACCACAACGGTTTCCCTCTAGAAATAATTTTGTTTAACT
TTAAGAAGGAGATATACAT**ATGAGCGATAACGGACCCCAGAACCAGCGTAATGCCCC**
**TCGCATTACTTTCGGAGGACCCAGCGATTCTACCGGATCTAATCAAAACGGGGAGCG**
**CTCCGGGGCGCGTTCCAAACAACGCCGTCCGCAAGGGCTTCCCAACAACACTGCAT**
**CGTGGTTTACAGCATTGACGCAACATGGGAAAGAAGACTTGAAGTTCCCACGCGGC**
**CAGGGAGTGCCCATTAATACAAATTCTTCGCCCGACGACCAGATCGGATATTATCGT**
**CGCGCAACTCGCCGCATCCGCGGAGGTGATGGTAAAATGAAGGATTTAAGCCCGCG**
**TTGGTATTTCTACTACTTGGGAACGGGTCCCGAAGCGGGATTACCATATGGAGCAAA**
**CAAAGATGGGATTATCTGGGTTGCTACTGAGGGAGCGTTGAATACGCCGAAGGATCA**

CATTGGTACGCGTAATCCAGCTAACAATGCTGCCATCGTTTTGCAACTTCCACAGGG
GACGACTCTGCCCAAGGGTTTCTACGCTGAAGGTTCGCGTGGGGGGTCTCAGGCAT
CTTCCCGTAGTTCGTCTCGTTCGCGCAACTCATCCCGTAATTCCACACCCGGCTCCT
CACGCGGCACGAGTCCGGCACGCATGGCAGGGAATGGGGGAGATGCTGCCCTGG
CTTTATTACTGCTTGATCGTTTGAACCAATTGGAATCCAAGATGTCTGGAAAAGGACA
ACAGCAACAAGGTCAGACTGTGACTAAGAAATCAGCAGCAGAGGCGTCCAAGAAAC
CACGTCAAAAACGCACAGCTACGAAGGCGTATAACGTGACACAGGCATTTGGTCGTC
GCGGCCCGGAACAAACCCAAGGTAATTTCGGCGACCAAGAACTGATCCGTCAAGGG
ACGGACTATAAGCATTGGCCTCAGATTGCCCAGTTCGCTCCCTCGGCATCGGCATTC
TTCGGCATGTCACGCATCGGAATGGAAGTGACACCTAGTGGGACCTGGCTGACTTAT
ACAGGCGCAATCAAGCTTGATGACAAAGACCCTAACTTCAAGGATCAAGTAATTCTGT
TAAATAAGCACATTGATGCGTATAAGACATTCCCGCCGACCGAGCCAAAGAAAGATA
AGAAAAAGAAAGCCGACGAAACTCAAGCCTTGCCGCAGCGTCAAAAAAAGCAGCAG
ACTGTCACTCTTTTGCCAGCAGCAGATTTAGATGATTTTTCTAAACAACTTCAACAGTC
GATGTCCTCAGCGGACAGCACACAGGCCAAGCGAGCTCCCGGGACCAGCatggTGAG
CAAGGGCGAGGAGCTGTTCACCGGGGTGGTGCCCATCCTGGTCGAGCTGGACGGCG
ACGTAAACGGCCACAAGTTCAGCGTGTCCGGCGAGGGCGAGGGCGATGCCACCTACG
GCAAGCTGACCCTGAAGTTCATCTGCACCACCGGCAAGCTGCCCGTGCCCTGGCCCA
CCCTCGTGACCACCCTGACCTACGGCGTGCAGTGCTTCAGCCGCTACCCCGACCACA
TGAAGCAGCACGACTTCTTCAAGTCCGCCATGCCCGAAGGCTACGTCCAGGAGCGCA
CCATCTTCTTCAAGGACGACGGCAACTACAAGACCCGCGCCGAGGTGAAGTTCGAGG
GCGACACCCTGGTGAACCGCATCGAGCTGAAGGGCATCGACTTCAAGGAGGACGGCA
ACATCCTGGGGCACAAGCTGGAGTACAACTACAACAGCCACAACGTCTATATCATGGC
CGACAAGCAGAAGAACGGCATCAAGGTGAACTTCAAGATCCGCCACAACATCGAGGAC
GGCAGCGTGCAGCTCGCCGACCACTACCAGCAGAACACCCCCATCGGCGACGGCCC
CGTGCTGCTGCCCGACAACCACTACCTGAGCACCCAGTCCGCCCTGAGCAAAGACCC
CAACGAGAAGCGCGATCACATGGTCCTGCTGGAGTTCGTGACCGCCGCCGGGATCAC
TCTCGGCATGGACGAGCTGTACAAGAGAAGAGCAAAGAAGGAAGccGACACAAAGACG
ATGACGACAAGTAATAAAAGGGCGAATTCCAGCACACTGGCGGCCGTTACTAGTGGAT
CCGGCTGCTAACAAAGCCCGAAAGGAAGCTGAGTTGGCTGCTGCCACCGCTGAGCAA
TAACTAGCATAACCCCTTGGGGCCTCTAAACGGGTCTTGAGGGGTTTTTTG

S1-GFP-HA
https://benchling.com/s/seq-1E26o3JywKLGF0WzDZpg?m=slm-R2849A4qFngsFHFtzwtL
TAATACGACTCACTATAGGGAGACCACAACGGTTTCCCTCTAGAAATAATTTTGTTTAACT
TTAAGAAGGAGATATACATATGTCACAATGTGTAAATTTAACAACTCGCACTCAATTGC
CACCAGCATACACAAATTCATTCACTCGTGGGGTGTATTATCCGGACAAGGTTTTCCG
TTCGTCCGTCCTTCACAGTACGCAAGACCTGTTTCTGCCTTTTTTTTCTAACGTCACTT
GGTTCCACGCGATCCACGTTAGTGGTACGAACGGGACAAAGCGTTTTGACAATCCAG
TACTGCCTTTCAATGACGGTGTCTACTTCGCCTCAACTGAAAAGTCGAACATTATCCG
TGGCTGGATTTTTGGTACTACGTTAGATTCTAAGACCCAAAGTTTGTTAATTGTGAATA
ACGCAACCAATGTCGTGATCAAAGTATGCGAGTTCCAGTTCTGTAACGACCCGTTTCT
GGGTGTGTACTATCACAAGAACAACAAAAGTTGGATGGAGAGTGAATTTCGCGTCTAT

TCCTCAGCGAATAACTGTACCTTCGAATACGTGTCCCAGCCGTTCTTGATGGACCTT
GAGGGAAAACAGGGTAACTTCAAGAATTTACGTGAGTTTGTGTTCAAAAATATTGACG
GTTATTTCAAGATCTACTCCAAGCATACTCCCATTAACCTTGTGCGTGACCTTCCCCA
AGGTTTCTCGGCCTTGGAGCCTCTTGTAGATCTTCCGATTGGAATTAATATCACACGC
TTCCAGACTTTACTGGCTTTACATCGTTCCTATCTGACGCCCGGTGATAGTTCCTCAG
GTTGGACCGCGGGAGCTGCAGCATACTACGTCGGATACCTTCAACCCCGTACTTTC
CTGTTAAAATATAATGAGAATGGCACTATTACCGATGCAGTCGACTGCGCGCTTGATC
CCTTATCAGAAACTAAGTGCACGTTAAAATCTTTCACAGTAGAAAAGGGGATTTACCA
GACCAGCAATTTTCGCGTCCAACCGACAGAAAGTATCGTGCGCTTCCCTAATATTAC
GAACTTGTGCCCTTTCGGGGAGGTTTTTAATGCTACTCGTTTTGCAAGCGTATACGCC
TGGAACCGCAAGCGTATCTCTAACTGTGTTGCGGACTATTCTGTATTGTACAATTCTG
CTTCGTTTTCAACGTTCAAGTGTTACGGTGTCTCGCCAACTAAACTTAACGATTTGTGT
TTTACCAATGTGTACGCTGACAGTTTCGTAATCCGTGGTGATGAGGTGCGTCAAATCG
CCCCAGGCCAGACTGGCAAAATCGCGGATTACAATTATAAACTTCCTGACGATTTTAC
TGGTTGTGTTATCGCATGGAATTCTAATAACCTGGATTCTAAAGTGGGCGGCAACTAT
AATTACTTATACCGCCTTTTTCGCAAGAGTAATCTTAAGCCTTTTGAACGTGATATTTC
AACTGAAATTTACCAAGCGGGGTCCACACCCTGTAACGGAGTGGAGGGGTTCAACT
GCTACTTTCCACTTCAGTCTTATGGTTTCCAACCCACTAACGGGGTCGGATATCAGCC
TTACCGCGTGGTCGTTCTTAGCTTCGAATTACTTCATGCACCAGCGACGGTGTGCGG
ACCTAAAAAGAGTACCAACCTTGTTAAAAACAAATGTGTGAATTTTAACTTCAATGGAC
TGACAGGCACCGGGGTACTTACTGAATCCAATAAAAAATTCCTTCCCTTTCAACAATT
CGGTCGCGACATCGCGGATACAACTGATGCGGTTCGTGATCCACAAACTCTGGAAAT
CTTGGATATCACTCCTTGTTCCTTCGGCGGTGTGTCGGTTATCACCCCTGGGACGAA
TACATCCAACCAGGTTGCCGTCTTATATCAAGACGTGAATTGCACCGAAGTTCCAGTA
GCGATTCATGCCGACCAGCTGACACCCACATGGCGCGTGTATAGTACCGGGAGTAA
CGTTTTTCAAACCCGTGCTGGTTGCTTAATTGGCGCAGAGCACGTAAACAACTCTTAT
GAATGTGATATCCCAATCGGAGCAGGAATCTGCGCTAGCTATCAGACCCAAACGAAT
TCGCCACGCCGCGCACGTAAGCGAGCTCCCGGGACCAGCatggTGAGCAAGGGCGAG
GAGCTGTTCACCGGGGTGGTGCCCATCCTGGTCGAGCTGGACGGCGACGTAAACGGC
CACAAGTTCAGCGTGTCCGGCGAGGGCGAGGGCGATGCCACCTACGGCAAGCTGACC
CTGAAGTTCATCTGCACCACCGGCAAGCTGCCCGTGCCCTGGCCCACCCTCGTGACC
ACCCTGACCTACGGCGTGCAGTGCTTCAGCCGCTACCCCGACCACATGAAGCAGCAC
GACTTCTTCAAGTCCGCCATGCCCGAAGGCTACGTCCAGGAGCGCACCATCTTCTTCA
AGGACGACGGCAACTACAAGACCCGCGCCGAGGTGAAGTTCGAGGGCGACACCCTG
GTGAACCGCATCGAGCTGAAGGGCATCGACTTCAAGGAGGACGGCAACATCCTGGGG
CACAAGCTGGAGTACAACTACAACAGCCACAACGTCTATATCATGGCCGACAAGCAGA
AGAACGGCATCAAGGTGAACTTCAAGATCCGCCACAACATCGAGGACGGCAGCGTGCA
GCTCGCCGACCACTACCAGCAGAACACCCCCATCGGCGACGGCCCCGTGCTGCTGC
CCGACAACCACTACCTGAGCACCCAGTCCGCCCTGAGCAAAGACCCCAACGAGAAGC
GCGATCACATGGTCCTGCTGGAGTTCGTGACCGCCGCCGGGATCACTCTCGGCATGG
ACGAGCTGTACAAGAGAAGAGCAAAGAAGGAAGccTACCCATACGATGTTCCAGATTAC
GCTTAATAAAAGGGCGAATTCCAGCACACTGGCGGCCGTTACTAGTGGATCCGGCTGC
TAACAAAGCCCGAAAGGAAGCTGAGTTGGCTGCTGCCACCGCTGAGCAATAACTAGCA
TAACCCCTTGGGGCCTCTAAACGGGTCTTGAGGGGTTTTTTG

S1 fragments
a (aa 1-100):
TCACAATGTGTAAATTTAACAACTCGCACTCAATTGCCACCAGCATACACAAATTCATTC
ACTCGTGGGGTGTATTATCCGGACAAGGTTTTCCGTTCGTCCGTCCTTCACAGTACGCA
AGACCTGTTTCTGCCTTTTTTTTCTAACGTCACTTGGTTCCACGCGATCCACGTTAGTGG
TACGAACGGGACAAAGCGTTTTGACAATCCAGTACTGCCTTTCAATGACGGTGTCTACT
TCGCCTCAACTGAAAAGTCGAACATTATCCGTGGCTGGATTTTTGGTACTACGTTAGATT
CT

b (aa 101-200):
AAGACCCAAAGTTTGTTAATTGTGAATAACGCAACCAATGTCGTGATCAAAGTATGCGAG
TTCCAGTTCTGTAACGACCCGTTTCTGGGTGTGTACTATCACAAGAACAACAAAAGTTGG
ATGGAGAGTGAATTTCGCGTCTATTCCTCAGCGAATAACTGTACCTTCGAATACGTGTCC
CAGCCGTTCTTGATGGACCTTGAGGGAAAACAGGGTAACTTCAAGAATTTACGTGAGTT
TGTGTTCAAAAATATTGACGGTTATTTCAAGATCTACTCCAAGCATACTCCCATTAACCTT

c (aa 201-300):
GTGCGTGACCTTCCCCAAGGTTTCTCGGCCTTGGAGCCTCTTGTAGATCTTCCGATTGG
AATTAATATCACACGCTTCCAGACTTTACTGGCTTTACATCGTTCCTATCTGACGCCCGG
TGATAGTTCCTCAGGTTGGACCGCGGGAGCTGCAGCATACTACGTCGGATACCTTCAA
CCCCGTACTTTCCTGTTAAAATATAATGAGAATGGCACTATTACCGATGCAGTCGACTGC
GCGCTTGATCCCTTATCAGAAACTAAGTGCACGTTAAAATCTTTCACAGTAGAAAAGGG
GATT

d (aa 301-400):
TACCAGACCAGCAATTTTCGCGTCCAACCGACAGAAAGTATCGTGCGCTTCCCTAATAT
TACGAACTTGTGCCCTTTCGGGGAGGTTTTTAATGCTACTCGTTTTGCAAGCGTATACGC
CTGGAACCGCAAGCGTATCTCTAACTGTGTTGCGGACTATTCTGTATTGTACAATTCTGC
TTCGTTTTCAACGTTCAAGTGTTACGGTGTCTCGCCAACTAAACTTAACGATTTGTGTTTT
ACCAATGTGTACGCTGACAGTTTCGTAATCCGTGGTGATGAGGTGCGTCAAATCGCCCC
A

e (aa 321-420):
ACGAACTTGTGCCCTTTCGGGGAGGTTTTTAATGCTACTCGTTTTGCAAGCGTATACGC
CTGGAACCGCAAGCGTATCTCTAACTGTGTTGCGGACTATTCTGTATTGTACAATTCTGC
TTCGTTTTCAACGTTCAAGTGTTACGGTGTCTCGCCAACTAAACTTAACGATTTGTGTTTT
ACCAATGTGTACGCTGACAGTTTCGTAATCCGTGGTGATGAGGTGCGTCAAATCGCCCC
AGGCCAGACTGGCAAAATCGCGGATTACAATTATAAACTTCCTGACGATTTTACTGGTTG
T

f (aa 351-450):
GCGGACTATTCTGTATTGTACAATTCTGCTTCGTTTTCAACGTTCAAGTGTTACGGTGTCT
CGCCAACTAAACTTAACGATTTGTGTTTTACCAATGTGTACGCTGACAGTTTCGTAATCC
GTGGTGATGAGGTGCGTCAAATCGCCCCAGGCCAGACTGGCAAAATCGCGGATTACAA

TTATAAACTTCCTGACGATTTTACTGGTTGTGTTATCGCATGGAATTCTAATAACCTGGAT
TCTAAAGTGGGCGGCAACTATAATTACTTATACCGCCTTTTTCGCAAGAGTAATCTTAAG

g (aa 381-480):
ACCAATGTGTACGCTGACAGTTTCGTAATCCGTGGTGATGAGGTGCGTCAAATCGCCCC
AGGCCAGACTGGCAAAATCGCGGATTACAATTATAAACTTCCTGACGATTTTACTGGTTG
TGTTATCGCATGGAATTCTAATAACCTGGATTCTAAAGTGGGCGGCAACTATAATTACTT
ATACCGCCTTTTTCGCAAGAGTAATCTTAAGCCTTTTGAACGTGATATTTCAACTGAAATT
TACCAAGCGGGGTCCACACCCTGTAACGGAGTGGAGGGGTTCAACTGCTACTTTCCAC
TT

h (aa 401-500):
GGCCAGACTGGCAAAATCGCGGATTACAATTATAAACTTCCTGACGATTTTACTGGTTGT
GTTATCGCATGGAATTCTAATAACCTGGATTCTAAAGTGGGCGGCAACTATAATTACTTA
TACCGCCTTTTTCGCAAGAGTAATCTTAAGCCTTTTGAACGTGATATTTCAACTGAAATTT
ACCAAGCGGGGTCCACACCCTGTAACGGAGTGGAGGGGTTCAACTGCTACTTTCCACT
TCAGTCTTATGGTTTCCAACCCACTAACGGGGTCGGATATCAGCCTTACCGCGTGGTCG
TT

i (aa 421-520):
GTTATCGCATGGAATTCTAATAACCTGGATTCTAAAGTGGGCGGCAACTATAATTACTTA
TACCGCCTTTTTCGCAAGAGTAATCTTAAGCCTTTTGAACGTGATATTTCAACTGAAATTT
ACCAAGCGGGGTCCACACCCTGTAACGGAGTGGAGGGGTTCAACTGCTACTTTCCACT
TCAGTCTTATGGTTTCCAACCCACTAACGGGGTCGGATATCAGCCTTACCGCGTGGTCG
TTCTTAGCTTCGAATTACTTCATGCACCAGCGACGGTGTGCGGACCTAAAAAGAGTACC
AAC

j (aa 451-550):
CCTTTTGAACGTGATATTTCAACTGAAATTTACCAAGCGGGGTCCACACCCTGTAACGG
AGTGGAGGGGTTCAACTGCTACTTTCCACTTCAGTCTTATGGTTTCCAACCCACTAACG
GGGTCGGATATCAGCCTTACCGCGTGGTCGTTCTTAGCTTCGAATTACTTCATGCACCA
GCGACGGTGTGCGGACCTAAAAAGAGTACCAACCTTGTTAAAAACAAATGTGTGAATTTT
AACTTCAATGGACTGACAGGCACCGGGGTACTTACTGAATCCAATAAAAAATTCCTTCC
CTTT

k (aa 501-600):
CTTAGCTTCGAATTACTTCATGCACCAGCGACGGTGTGCGGACCTAAAAAGAGTACCAA
CCTTGTTAAAAACAAATGTGTGAATTTTAACTTCAATGGACTGACAGGCACCGGGGTACT
TACTGAATCCAATAAAAAATTCCTTCCCTTTCAACAATTCGGTCGCGACATCGCGGATAC
AACTGATGCGGTTCGTGATCCACAAACTCTGGAAATCTTGGATATCACTCCTTGTTCCTT
CGGCGGTGTGTCGGTTATCACCCCTGGGACGAATACATCCAACCAGGTTGCCGTCTTA
TAT

I (aa 574-673):
GATATCACTCCTTGTTCCTTCGGCGGTGTGTCGGTTATCACCCCTGGGACGAATACATC
CAACCAGGTTGCCGTCTTATATCAAGACGTGAATTGCACCGAAGTTCCAGTAGCGATTC
ATGCCGACCAGCTGACACCCACATGGCGCGTGTATAGTACCGGGAGTAACGTTTTTCA
AACCCGTGCTGGTTGCTTAATTGGCGCAGAGCACGTAAACAACTCTTATGAATGTGATAT
CCCAATCGGAGCAGGAATCTGCGCTAGCTATCAGACCCAAACGAATTCGCCACGCCGC
GCACGT

GFP-S1-HA
https://benchling.com/s/seq-eOckhlPMa3hJ4Im5pktG?m=slm-OT7SIncH8qzNzymz16cp
TAATACGACTCACTATAGGGAGACCACAACGGTTTCCCTCTAGAAATAATTTTGTTTAACT
TTAAGAAGGAGATATACATatggTGAGCAAGGGCGAGGAGCTGTTCACCGGGGTGGTGC
CCATCCTGGTCGAGCTGGACGGCGACGTAAACGGCCACAAGTTCAGCGTGTCCGGCG
AGGGCGAGGGCGATGCCACCTACGGCAAGCTGACCCTGAAGTTCATCTGCACCACCG
GCAAGCTGCCCGTGCCCTGGCCCACCCTCGTGACCACCCTGACCTACGGCGTGCAGT
GCTTCAGCCGCTACCCCGACCACATGAAGCAGCACGACTTCTTCAAGTCCGCCATGCC
CGAAGGCTACGTCCAGGAGCGCACCATCTTCTTCAAGGACGACGGCAACTACAAGACC
CGCGCCGAGGTGAAGTTCGAGGGCGACACCCTGGTGAACCGCATCGAGCTGAAGGG
CATCGACTTCAAGGAGGACGGCAACATCCTGGGGCACAAGCTGGAGTACAACTACAAC
AGCCACAACGTCTATATCATGGCCGACAAGCAGAAGAACGGCATCAAGGTGAACTTCA
AGATCCGCCACAACATCGAGGACGGCAGCGTGCAGCTCGCCGACCACTACCAGCAGA
ACACCCCCATCGGCGACGGCCCCGTGCTGCTGCCCGACAACCACTACCTGAGCACCC
AGTCCGCCCTGAGCAAAGACCCCAACGAGAAGCGCGATCACATGGTCCTGCTGGAGTT
CGTGACCGCCGCCGGGATCACTCTCGGCATGGACGAGCTGTACAAGAAGCGAGCTCC
CGGGACCAGC**ATGTCACAATGTGTAAATTTAACAACTCGCACTCAATTGCCACCAGCA**
**TACACAAATTCATTCACTCGTGGGGTGTATTATCCGGACAAGGTTTTCCGTTCGTCCG**
**TCCTTCACAGTACGCAAGACCTGTTTCTGCCTTTTTTTTCTAACGTCACTTGGTTCCAC**
**GCGATCCACGTTAGTGGTACGAACGGGACAAAGCGTTTTGACAATCCAGTACTGCCT**
**TTCAATGACGGTGTCTACTTCGCCTCAACTGAAAAGTCGAACATTATCCGTGGCTGGA**
**TTTTTGGTACTACGTTAGATTCTAAGACCCAAAGTTTGTTAATTGTGAATAACGCAACC**
**AATGTCGTGATCAAAGTATGCGAGTTCCAGTTCTGTAACGACCCGTTTCTGGGTGTGT**
**ACTATCACAAGAACAACAAAAGTTGGATGGAGAGTGAATTTCGCGTCTATTCCTCAGC**
**GAATAACTGTACCTTCGAATACGTGTCCCAGCCGTTCTTGATGGACCTTGAGGGAAA**
**ACAGGGTAACTTCAAGAATTTACGTGAGTTTGTGTTCAAAAATATTGACGGTTATTTCA**
**AGATCTACTCCAAGCATACTCCCATTAACCTTGTGCGTGACCTTCCCCAAGGTTTCTC**
**GGCCTTGGAGCCTCTTGTAGATCTTCCGATTGGAATTAATATCACACGCTTCCAGACT**
**TTACTGGCTTTACATCGTTCCTATCTGACGCCCGGTGATAGTTCCTCAGGTTGGACC**
**GCGGGAGCTGCAGCATACTACGTCGGATACCTTCAACCCCGTACTTTCCTGTTAAAA**
**TATAATGAGAATGGCACTATTACCGATGCAGTCGACTGCGCGCTTGATCCCTTATCAG**
**AAACTAAGTGCACGTTAAAATCTTTCACAGTAGAAAAGGGGATTTACCAGACCAGCAA**
**TTTTCGCGTCCAACCGACAGAAAGTATCGTGCGCTTCCCTAATATTACGAACTTGTGC**
**CCTTTCGGGGAGGTTTTTAATGCTACTCGTTTTGCAAGCGTATACGCCTGGAACCGC**
**AAGCGTATCTCTAACTGTGTTGCGGACTATTCTGTATTGTACAATTCTGCTTCGTTTTC**
**AACGTTCAAGTGTTACGGTGTCTCGCCAACTAAACTTAACGATTTGTGTTTTACCAATG**

**TGTACGCTGACAGTTTCGTAATCCGTGGTGATGAGGTGCGTCAAATCGCCCCAGGC**
**CAGACTGGCAAAATCGCGGATTACAATTATAAACTTCCTGACGATTTTACTGGTTGTG**
**TTATCGCATGGAATTCTAATAACCTGGATTCTAAAGTGGGCGGCAACTATAATTACTTA**
**TACCGCCTTTTTCGCAAGAGTAATCTTAAGCCTTTTGAACGTGATATTTCAACTGAAAT**
**TTACCAAGCGGGGTCCACACCCTGTAACGGAGTGGAGGGGTTCAACTGCTACTTTC**
**CACTTCAGTCTTATGGTTTCCAACCCACTAACGGGGTCGGATATCAGCCTTACCGCG**
**TGGTCGTTCTTAGCTTCGAATTACTTCATGCACCAGCGACGGTGTGCGGACCTAAAA**
**AGAGTACCAACCTTGTTAAAAACAAATGTGTGAATTTTAACTTCAATGGACTGACAGG**
**CACCGGGGTACTTACTGAATCCAATAAAAAATTCCTTCCCTTTCAACAATTCGGTCGC**
**GACATCGCGGATACAACTGATGCGGTTCGTGATCCACAAACTCTGGAAATCTTGGAT**
**ATCACTCCTTGTTCCTTCGGCGGTGTGTCGGTTATCACCCCTGGGACGAATACATCC**
**AACCAGGTTGCCGTCTTATATCAAGACGTGAATTGCACCGAAGTTCCAGTAGCGATT**
**CATGCCGACCAGCTGACACCCACATGGCGCGTGTATAGTACCGGGAGTAACGTTTTT**
**CAAACCCGTGCTGGTTGCTTAATTGGCGCAGAGCACGTAAACAACTCTTATGAATGT**
**GATATCCCAATCGGAGCAGGAATCTGCGCTAGCTATCAGACCCAAACGAATTCGCCA**
**CGCCGCGCACGT**AGAAGAGCAAAGAAGGAAGccTACCCATACGATGTTCCAGATTACG
CTTAATAAAAGGGCGAATTCCAGCACACTGGCGGCCGTTACTAGTGGATCCGGCTGCT
AACAAAGCCCGAAAGGAAGCTGAGTTGGCTGCTGCCACCGCTGAGCAATAACTAGCAT
AACCCCTTGGGGCCTCTAAACGGGTCTTGAGGGGTTTTTTG

RBD2-GFP-HA
TAATACGACTCACTATAGGGAGACCACAACGGTTTCCCTCTAGAAATAATTTTGTTTAACT
TTAAGAAGGAGATATACAT**ATGAACATTACAAACTTGTGTCCATTTGGGGAAGTTTTTAA**
**TGCAACTCGCTTTGCCAGTGTGTATGCCTGGAATCGTAAGCGTATTAGTAACTGTGTT**
**GCTGACTATAGCGTACTTTACAATTCCGCCAGCTTCAGTACATTTAAGTGTTACGGTG**
**TGTCCCCGACTAAGTTAAATGACCTTTGCTTCACAAATGTATACGCAGACTCATTCGT**
**TATTCGTGGAGATGAGGTTCGCCAGATCGCACCCGGCCAAACCGGTAAGATCGCCG**
**ATTACAACTATAAGTTGCCTGACGATTTTACTGGTTGTGTTATTGCATGGAATTCGAAT**
**AATTTGGATAGTAAGGTTGGCGGTAACTACAACTATCTTTATCGCCTTTTTCGCAAATC**
**CAATCTGAAGCCCTTCGAACGCGACATCTCGACCGAGATTTACCAAGCAGGTTCAAC**
**GCCTTGCAATGGAGTAGAGGGTTTTAACTGCTATTTTCCATTACAGTCCTACGGATTT**
**CAGCCAACGAATGGTGTAGGTTACCAGCCGTACCGTGTCGTAGTGCTTAGTTTTGAG**
**CTGTTACATGCGCCCGCCACTGTT**AAGCGAGCTCCCGGGACCAGCatggTGAGCAAGG
GCGAGGAGCTGTTCACCGGGGTGGTGCCCATCCTGGTCGAGCTGGACGGCGACGTAA
ACGGCCACAAGTTCAGCGTGTCCGGCGAGGGCGAGGGCGATGCCACCTACGGCAAG
CTGACCCTGAAGTTCATCTGCACCACCGGCAAGCTGCCCGTGCCCTGGCCCACCCTC
GTGACCACCCTGACCTACGGCGTGCAGTGCTTCAGCCGCTACCCCGACCACATGAAG
CAGCACGACTTCTTCAAGTCCGCCATGCCCGAAGGCTACGTCCAGGAGCGCACCATCT
TCTTCAAGGACGACGGCAACTACAAGACCCGCGCCGAGGTGAAGTTCGAGGGCGACA
CCCTGGTGAACCGCATCGAGCTGAAGGGCATCGACTTCAAGGAGGACGGCAACATCC
TGGGGCACAAGCTGGAGTACAACTACAACAGCCACAACGTCTATATCATGGCCGACAA
GCAGAAGAACGGCATCAAGGTGAACTTCAAGATCCGCCACAACATCGAGGACGGCAG
CGTGCAGCTCGCCGACCACTACCAGCAGAACACCCCCATCGGCGACGGCCCCGTGCT
GCTGCCCGACAACCACTACCTGAGCACCCAGTCCGCCCTGAGCAAAGACCCCAACGA

GAAGCGCGATCACATGGTCCTGCTGGAGTTCGTGACCGCCGCCGGGATCACTCTCGG
CATGGACGAGCTGTACAAGAGAAGAGCAAAGAAGGAAGccTACCCATACGATGTTCCAG
ATTACGCTTAATAAAAGGGCGAATTCCAGCACACTGGCGGCCGTTACTAGTGGATCCG
GCTGCTAACAAAGCCCGAAAGGAAGCTGAGTTGGCTGCTGCCACCGCTGAGCAATAAC
TAGCATAACCCCTTGGGGCCTCTAAACGGGTCTTGAGGGGTTTTTTG

GFP-RBD2-HA
TAATACGACTCACTATAGGGAGACCACAACGGTTTCCCTCTAGAAATAATTTTGTTTAACT
TTAAGAAGGAGATATACATatggTGAGCAAGGGCGAGGAGCTGTTCACCGGGGTGGTGC
CCATCCTGGTCGAGCTGGACGGCGACGTAAACGGCCACAAGTTCAGCGTGTCCGGCG
AGGGCGAGGGCGATGCCACCTACGGCAAGCTGACCCTGAAGTTCATCTGCACCACCG
GCAAGCTGCCCGTGCCCTGGCCCACCCTCGTGACCACCCTGACCTACGGCGTGCAGT
GCTTCAGCCGCTACCCCGACCACATGAAGCAGCACGACTTCTTCAAGTCCGCCATGCC
CGAAGGCTACGTCCAGGAGCGCACCATCTTCTTCAAGGACGACGGCAACTACAAGACC
CGCGCCGAGGTGAAGTTCGAGGGCGACACCCTGGTGAACCGCATCGAGCTGAAGGG
CATCGACTTCAAGGAGGACGGCAACATCCTGGGGCACAAGCTGGAGTACAACTACAAC
AGCCACAACGTCTATATCATGGCCGACAAGCAGAAGAACGGCATCAAGGTGAACTTCA
AGATCCGCCACAACATCGAGGACGGCAGCGTGCAGCTCGCCGACCACTACCAGCAGA
ACACCCCCATCGGCGACGGCCCCGTGCTGCTGCCCGACAACCACTACCTGAGCACCC
AGTCCGCCCTGAGCAAAGACCCCAACGAGAAGCGCGATCACATGGTCCTGCTGGAGTT
CGTGACCGCCGCCGGGATCACTCTCGGCATGGACGAGCTGTACAAGAAGCGAGCTCC
CGGGACCAGC**ATGAACATTACAAACTTGTGTCCATTTGGGGAAGTTTTTAATGCAACT
CGCTTTGCCAGTGTGTATGCCTGGAATCGTAAGCGTATTAGTAACTGTGTTGCTGACT
ATAGCGTACTTTACAATTCCGCCAGCTTCAGTACATTTAAGTGTTACGGTGTGTCCCC
GACTAAGTTAAATGACCTTTGCTTCACAAATGTATACGCAGACTCATTCGTTATTCGTG
GAGATGAGGTTCGCCAGATCGCACCCGGCCAAACCGGTAAGATCGCCGATTACAAC
TATAAGTTGCCTGACGATTTTACTGGTTGTGTTATTGCATGGAATTCGAATAATTTGGA
TAGTAAGGTTGGCGGTAACTACAACTATCTTTATCGCCTTTTTCGCAAATCCAATCTGA
AGCCCTTCGAACGCGACATCTCGACCGAGATTTACCAAGCAGGTTCAACGCCTTGCA
ATGGAGTAGAGGGTTTTAACTGCTATTTTCCATTACAGTCCTACGGATTTCAGCCAAC
GAATGGTGTAGGTTACCAGCCGTACCGTGTCGTAGTGCTTAGTTTTGAGCTGTTACAT
GCGCCCGCCACTGTT**AGAAGAGCAAAGAAGGAAGccTACCCATACGATGTTCCAGATT
ACGCTTAATAAAAGGGCGAATTCCAGCACACTGGCGGCCGTTACTAGTGGATCCGGCT
GCTAACAAAGCCCGAAAGGAAGCTGAGTTGGCTGCTGCCACCGCTGAGCAATAACTAG
CATAACCCCTTGGGGCCTCTAAACGGGTCTTGAGGGGTTTTTTG

Sequence of additional 4 amino acids adding one Cys at the C-term of RBD:
TGCGGACCTAAA

GFP-Omicron-HA
TAATACGACTCACTATAGGGAGACCACAACGGTTTCCCTCTAGAAATAATTTTGTTTAACT
TTAAGAAGGAGATATACATatggTGAGCAAGGGCGAGGAGCTGTTCACCGGGGTGGTGC
CCATCCTGGTCGAGCTGGACGGCGACGTAAACGGCCACAAGTTCAGCGTGTCCGGCG
AGGGCGAGGGCGATGCCACCTACGGCAAGCTGACCCTGAAGTTCATCTGCACCACCG

GCAAGCTGCCCGTGCCCTGGCCCACCCTCGTGACCACCCTGACCTACGGCGTGCAGT
GCTTCAGCCGCTACCCCGACCACATGAAGCAGCACGACTTCTTCAAGTCCGCCATGCC
CGAAGGCTACGTCCAGGAGCGCACCATCTTCTTCAAGGACGACGGCAACTACAAGACC
CGCGCCGAGGTGAAGTTCGAGGGCGACACCCTGGTGAACCGCATCGAGCTGAAGGG
CATCGACTTCAAGGAGGACGGCAACATCCTGGGGCACAAGCTGGAGTACAACTACAAC
AGCCACAACGTCTATATCATGGCCGACAAGCAGAAGAACGGCATCAAGGTGAACTTCA
AGATCCGCCACAACATCGAGGACGGCAGCGTGCAGCTCGCCGACCACTACCAGCAGA
ACACCCCCATCGGCGACGGCCCCGTGCTGCTGCCCGACAACCACTACCTGAGCACCC
AGTCCGCCCTGAGCAAAGACCCCAACGAGAAGCGCGATCACATGGTCCTGCTGGAGTT
CGTGACCGCCGCCGGGATCACTCTCGGCATGGACGAGCTGTACAAGAAGCGAGCTCC
CGGGACCAGC**ATGAACATTACAAACTTGTGTCCATTTGATGAAGTTTTTAATGCAACTC**
**GCTTTGCCAGTGTGTATGCCTGGAATCGTAAGCGTATTAGTAACTGTGTTGCTGACTA**
**TAGCGTACTTTACAATCTCGCCCCATTCTTTACATTTAAGTGTTACGGTGTGTCCCCG**
**ACTAAGTTAAATGACCTTTGCTTCACAAATGTATACGCAGACTCATTCGTTATTCGTGG**
**AGATGAGGTTCGCCAGATCGCACCCGGCCAA**ACCGGTAATATCGCCGATTACAACTA
TAAGTTGCCTGACGATTTTACTGGTTGTGTTATTGCATGGAATTCGAATAAGTTGGATAGT
AAGGTTAGCGGTAACTACAACTATCTTTATCGCCTTTTTCGCAAATCCAATCTGAAGCCC
TTCGAACGCGACATCTCGACCGAGATTTACCAAGCAGGTAATAAGCCTTGCAATGGAGT
AGCAGGTTTTAACTGCTATTTTCCATTAAAGTCCTACTCGTTTCGTCCAACGTATGGTGTA
GGTCATCAGCCGTACCGTGTCGTAGTGCTTAGTTTTGAGCTGTTACATGCGCCCGCCAC
TGTTAGAAGAGCAAAGAAGGAAGccTACCCATACGATGTTCCAGATTACGCTTAATAAAA
GGGCGAATTCCAGCACACTGGCGGCCGTTACTAGTGGATCCGGCTGCTAACAAAGCC
CGAAAGGAAGCTGAGTTGGCTGCTGCCACCGCTGAGCAATAACTAGCATAACCCCTTG
GGGCCTCTAAACGGGTCTTGAGGGGTTTTTTG

RBD1-GFP-HA
https://benchling.com/s/seq-O5LRh9W8L1ETIzNW2aBj?m=slm-jm791mTR2HrJDX9nuvoE
TAATACGACTCACTATAGGGAGACCACAACGGTTTCCCTCTAGAAATAATTTTGTTTAACT
TTAAGAAGGAGATATACAT**ATGAACATCACCAACCTTTGCCCTTTTGGGGAAGTTTTTA**
**ACGCAACAAAATTCCCAAGCGTCTACGCCTGGGAACGCAAGAAAATCTCGAATTGCG**
**TGGCGGATTACTCCGTGCTTTATAATAGCACATTCTTTAGCACGTTCAAATGCTATGG**
**AGTTTCGGCAACGAAATTAAATGACTTATGTTTCTCGAACGTATATGCTGACTCCTTCG**
**TGGTAAAAGGTGACGATGTCCGCCAGATTGCTCCGGGTCAAACAGGGGTAATCGCT**
**GACTATAACTACAAGCTGCCGGACGATTTTATGGGTTGTGTTCTGGCGTGGAATACA**
**CGCAACATTGATGCCACATCAACGGGTAACTATAACTACAAATATCGCTACCTTCGTC**
**ACGGCAAATTACGTCCGTTTGAGCGCGACATTTCTAACGTACCCTTCTCACCGGATG**
**GCAAACCTTGTACACCTCCAGCTTTAAACTGCTACTGGCCTCTTAACGACTACGGGTT**
**CTATACAACAACCGGAATTGGATATCAGCCATACCGTGTTGTTGTATTGTCATTTGAA**
**CTGCTTAACGCGCCCGCTACAGTT**AAGCGAGCTCCCGGGACCAGCatggTGAGCAAGG
GCGAGGAGCTGTTCACCGGGGTGGTGCCCATCCTGGTCGAGCTGGACGGCGACGTAA
ACGGCCACAAGTTCAGCGTGTCCGGCGAGGGCGAGGGCGATGCCACCTACGGCAAG
CTGACCCTGAAGTTCATCTGCACCACCGGCAAGCTGCCCGTGCCCTGGCCCACCCTC
GTGACCACCCTGACCTACGGCGTGCAGTGCTTCAGCCGCTACCCCGACCACATGAAG

CAGCACGACTTCTTCAAGTCCGCCATGCCCGAAGGCTACGTCCAGGAGCGCACCATCT
TCTTCAAGGACGACGGCAACTACAAGACCCGCGCCGAGGTGAAGTTCGAGGGCGACA
CCCTGGTGAACCGCATCGAGCTGAAGGGCATCGACTTCAAGGAGGACGGCAACATCC
TGGGGCACAAGCTGGAGTACAACTACAACAGCCACAACGTCTATATCATGGCCGACAA
GCAGAAGAACGGCATCAAGGTGAACTTCAAGATCCGCCACAACATCGAGGACGGCAG
CGTGCAGCTCGCCGACCACTACCAGCAGAACACCCCCATCGGCGACGGCCCCGTGCT
GCTGCCCGACAACCACTACCTGAGCACCCAGTCCGCCCTGAGCAAAGACCCCAACGA
GAAGCGCGATCACATGGTCCTGCTGGAGTTCGTGACCGCCGCCGGGATCACTCTCGG
CATGGACGAGCTGTACAAGAGAAGAGCAAAGAAGGAAGCCTACCCATACGATGTTCCA
GATTACGCTTAATAAAAGGGCGAATTCCAGCACACTGGCGGCCGTTACTAGTGGATCC
GGCTGCTAACAAAGCCCGAAAGGAAGCTGAGTTGGCTGCTGCCACCGCTGAGCAATAA
CTAGCATAACCCCTTGGGGCCTCTAAACGGGTCTTGAGGGGTTTTTTG

GFP-RBD1-HA

TAATACGACTCACTATAGGGAGACCACAACGGTTTCCCTCTAGAAATAATTTTGTTTAACT
TTAAGAAGGAGATATACATatggTGAGCAAGGGCGAGGAGCTGTTCACCGGGGTGGTGC
CCATCCTGGTCGAGCTGGACGGCGACGTAAACGGCCACAAGTTCAGCGTGTCCGGCG
AGGGCGAGGGCGATGCCACCTACGGCAAGCTGACCCTGAAGTTCATCTGCACCACCG
GCAAGCTGCCCGTGCCCTGGCCCACCCTCGTGACCACCCTGACCTACGGCGTGCAGT
GCTTCAGCCGCTACCCCGACCACATGAAGCAGCACGACTTCTTCAAGTCCGCCATGCC
CGAAGGCTACGTCCAGGAGCGCACCATCTTCTTCAAGGACGACGGCAACTACAAGACC
CGCGCCGAGGTGAAGTTCGAGGGCGACACCCTGGTGAACCGCATCGAGCTGAAGGG
CATCGACTTCAAGGAGGACGGCAACATCCTGGGGCACAAGCTGGAGTACAACTACAAC
AGCCACAACGTCTATATCATGGCCGACAAGCAGAAGAACGGCATCAAGGTGAACTTCA
AGATCCGCCACAACATCGAGGACGGCAGCGTGCAGCTCGCCGACCACTACCAGCAGA
ACACCCCCATCGGCGACGGCCCCGTGCTGCTGCCCGACAACCACTACCTGAGCACCC
AGTCCGCCCTGAGCAAAGACCCCAACGAGAAGCGCGATCACATGGTCCTGCTGGAGTT
CGTGACCGCCGCCGGGATCACTCTCGGCATGGACGAGCTGTACAAGAAGCGAGCTCC
CGGGACCAGC**ATGAACATCACCAACCTTTGCCCTTTTGGGGAAGTTTTTAACGCAACA**
**AAATTCCCAAGCGTCTACGCCTGGGAACGCAAGAAAATCTCGAATTGCGTGGCGGAT**
**TACTCCGTGCTTTATAATAGCACATTCTTTAGCACGTTCAAATGCTATGGAGTTTCGGC**
**AACGAAATTAAATGACTTATGTTTCTCGAACGTATATGCTGACTCCTTCGTGGTAAAAG**
**GTGACGATGTCCGCCAGATTGCTCCGGGTCAAACAGGGGTAATCGCTGACTATAACT**
**ACAAGCTGCCGGACGATTTTATGGGTTGTGTTCTGGCGTGGAATACACGCAACATTG**
**ATGCCACATCAACGGGTAACTATAACTACAAATATCGCTACCTTCGTCACGGCAAATT**
**ACGTCCGTTTGAGCGCGACATTTCTAACGTACCCTTCTCACCGGATGGCAAACCTTG**
**TACACCTCCAGCTTTAAACTGCTACTGGCCTCTTAACGACTACGGGTTCTATACAACA**
**ACCGGAATTGGATATCAGCCATACCGTGTTGTTGTATTGTCATTTGAACTGCTTAACG**
**CGCCCGCTACAGTT**AGAAGAGCAAAGAAGGAAGccTACCCATACGATGTTCCAGATTAC
GCTTAATAAAAGGGCGAATTCCAGCACACTGGCGGCCGTTACTAGTGGATCCGGCTGC
TAACAAAGCCCGAAAGGAAGCTGAGTTGGCTGCTGCCACCGCTGAGCAATAACTAGCA
TAACCCCTTGGGGCCTCTAAACGGGTCTTGAGGGGTTTTTTG

ACE2-HA

https://benchling.com/s/seq-Js6y4qByzY6f0hRnmlC1?m=slm-LHt8GIsHhIoVK7Bnr3dB
TAATACGACTCACTATAGGGAGACCACAACGGTTTCCCTCTAGAAATAATTTTGTTTAACT
TTAAGAAGGAGATATACCATGCAGTCCACCATTGAGGAACAGGCCAAGACATTTTTGG
ACAAGTTTAACCACGAAGCCGAAGACCTGTTCTATCAAAGTTCACTTGCTTCTTGGAA
TTATAACACCAATATTACTGAAGAGAATGTCCAAAACATGAATAACGCTGGGGACAAA
TGGTCTGCCTTTTTAAAGGAACAGTCCACACTTGCCCAAATGTATCCACTACAAGAAA
TTCAGAATCTCACAGTCAAGCTTCAGCTGCAGGCTCTTCAGCAAAATGGGTCTTCAGT
GCTCTCAGAAGACAAGAGCAAACGGTTGAACACAATTCTAAATACAATGAGCACCAT
CTACAGTACTGGAAAAGTTTGTAACCCAGATAATCCACAAGAATGCTTATTACTTGAA
CCAGGTTTGAATGAAATAATGGCAAACAGTTTAGACTACAATGAGAGGCTCTGGGCTT
GGGAAAGCTGGAGATCTGAGGTCGGCAAGCAGCTGAGGCCATTATATGAAGAGTAT
GTGGTCTTGAAAAATGAGATGGCAAGAGCAAATCATTATGAGGACTATGGGGATTATT
GGAGAGGAGACTATGAAGTAAATGGGGTAGATGGCTATGACTACAGCCGCGGCCAG
TTGATTGAAGATGTGGAACATACCTTTGAAGAGATTAAACCATTATATGAACATCTTCA
TGCCTATGTGAGGGCAAAGTTGATGAATGCCTATCCTTCCTATATCAGTCCAATTGGA
TGCCTCCCTGCTCATTTGCTTGGTGATATGTGGGGTAGATTTTGGACAAATCTGTACT
CTTTGACAGTTCCCTTTGGACAGAAACCAAACATAGATGTTACTGATGCAATGGTGGA
CCAGGCCTGGGATGCACAGAGAATATTCAAGGAGGCCGAGAAGTTCTTTGTATCTGT
TGGTCTTCCTAATATGACTCAAGGATTCTGGGAAAATTCCATGCTAACGGACCCAGGA
AATGTTCAGAAAGCAGTCTGCCATCCCACAGCTTGGGACCTGGGGAAAGGCGACTT
CAGGATCCTTATGTGCACAAAGGTGACAATGGACGACTTCCTGACAGCTCATCATGA
GATGGGGCATATTCAGTATGATATGGCATATGCTGCACAACCTTTTCTGCTAAGAAAT
GGAGCTAATGAAGGATTCCATGAAGCTGTTGGGGAAATCATGTCACTTTCTGCAGCC
ACACCTAAGCATTTAAAATCCATTGGTCTTCTGTCACCCGATTTTCAAGAAGACAATG
AAACAGAAATAAACTTCCTGCTCAAACAAGCACTCACGATTGTTGGGACTCTGCCATT
TACTTACATGTTAGAGAAGTGGAGGTGGATGGTCTTTAAAGGGGAAATTCCCAAAGAC
CAGTGGATGAAAAAGTGGTGGGAGATGAAGCGAGAGATAGTTGGGGTGGTGGAACC
TGTGCCCCATGATGAAACATACTGTGACCCCGCATCTCTGTTCCATGTTTCTAATGAT
TACTCATTCATTCGATATTACACAAGGACCCTTTACCAATTCCAGTTTCAAGAAGCACT
TTGTCAAGCAGCTAAACATGAAGGCCCTCTGCACAAATGTGACATCTCAAACTCTACA
GAAGCTGGACAGAAACTGTTCAATATGCTGAGGCTTGGAAAATCAGAACCCTGGACC
CTAGCATTGGAAAATGTTGTAGGAGCAAAGAACATGAATGTAAGGCCACTGCTCAAC
TACTTTGAGCCCTTATTTACCTGGCTGAAAGACCAGAACAAGAATTCTTTTGTGGGAT
GGAGTACCGACTGGAGTCCATATGCAGACCAAAGCATCAAAGTGAGGATAAGCCTAA
AATCAGCTCTTGGAGATAGAGCATATGAATGGAACGACAATGAAATGTACCTGTTCCG
ATCATCTGTTGCATATGCTATGAGGCAGTACTTTTTAAAAGTAAAAAATCAGATGATTC
TTTTTGGGGAGGAGGATGTGCGAGTGGCTAATTTGAAACCAAGAATCTCCTTTAATTT
CTTTGTCACTGCACCTAAAAATGTGTCTGATATCATTCCTAGAACTGAAGTTGAAAAG
GCCATCAGGATGTCCCGGAGCCGTATCAATGATGCTTTCCGTCTGAATGACAACAGC
CTAGAGTTTCTGGGGATACAGCCAACACTTGGACCTCCTAACCAGCCCCCTGTTTCC
AGTGGTGGCGGAGGGACCAGCTACCCATACGATGTTCCAGATTACGCTTAATAAAAGG
GCGAATTCCAGCACACTGGCGGCCGTTACTAGTGGATCCGGCTGCTAACAAAGCCCGA
AAGGAAGCTGAGTTGGCTGCTGCCACCGCTGAGCAATAACTAGCATAACCCCTTGGGG
CCTCTAAACGGGTCTTGAGGGGTTTTTTG

ACE2-FLAG

https://benchling.com/s/seq-BLfvTnsAuLqTyAsPJTzc?m=slm-XRJLb6UlQdBomz068EBF

TAATACGACTCACTATAGGGAGACCACAACGGTTTCCCTCTAGAAATAATTTTGTTTAACT
TTAAGAAGGAGATATACCATGCAGTCCACCATTGAGGAACAGGCCAAGACATTTTTGG
ACAAGTTTAACCACGAAGCCGAAGACCTGTTCTATCAAAGTTCACTTGCTTCTTGGAA
TTATAACACCAATATTACTGAAGAGAATGTCCAAAACATGAATAACGCTGGGGACAAA
TGGTCTGCCTTTTTAAAGGAACAGTCCACACTTGCCCAAATGTATCCACTACAAGAAA
TTCAGAATCTCACAGTCAAGCTTCAGCTGCAGGCTCTTCAGCAAAATGGGTCTTCAGT
GCTCTCAGAAGACAAGAGCAAACGGTTGAACACAATTCTAAATACAATGAGCACCAT
CTACAGTACTGGAAAAGTTTGTAACCCAGATAATCCACAAGAATGCTTATTACTTGAA
CCAGGTTTGAATGAAATAATGGCAAACAGTTTAGACTACAATGAGAGGCTCTGGGCTT
GGGAAAGCTGGAGATCTGAGGTCGGCAAGCAGCTGAGGCCATTATATGAAGAGTAT
GTGGTCTTGAAAAATGAGATGGCAAGAGCAAATCATTATGAGGACTATGGGGATTATT
GGAGAGGAGACTATGAAGTAAATGGGGTAGATGGCTATGACTACAGCCGCGGCCAG
TTGATTGAAGATGTGGAACATACCTTTGAAGAGATTAAACCATTATATGAACATCTTCA
TGCCTATGTGAGGGCAAAGTTGATGAATGCCTATCCTTCCTATATCAGTCCAATTGGA
TGCCTCCCTGCTCATTTGCTTGGTGATATGTGGGGTAGATTTTGGACAAATCTGTACT
CTTTGACAGTTCCCTTTGGACAGAAACCAAACATAGATGTTACTGATGCAATGGTGGA
CCAGGCCTGGGATGCACAGAGAATATTCAAGGAGGCCGAGAAGTTCTTTGTATCTGT
TGGTCTTCCTAATATGACTCAAGGATTCTGGGAAAATTCCATGCTAACGGACCCAGGA
AATGTTCAGAAAGCAGTCTGCCATCCCACAGCTTGGGACCTGGGGAAAGGCGACTT
CAGGATCCTTATGTGCACAAAGGTGACAATGGACGACTTCCTGACAGCTCATCATGA
GATGGGGCATATTCAGTATGATATGGCATATGCTGCACAACCTTTTCTGCTAAGAAAT
GGAGCTAATGAAGGATTCCATGAAGCTGTTGGGGAAATCATGTCACTTTCTGCAGCC
ACACCTAAGCATTTAAAATCCATTGGTCTTCTGTCACCCGATTTTCAAGAAGACAATG
AAACAGAAATAAACTTCCTGCTCAAACAAGCACTCACGATTGTTGGGACTCTGCCATT
TACTTACATGTTAGAGAAGTGGAGGTGGATGGTCTTTAAAGGGGAAATTCCCAAAGAC
CAGTGGATGAAAAAGTGGTGGGAGATGAAGCGAGAGATAGTTGGGGTGGTGGAACC
TGTGCCCCATGATGAAACATACTGTGACCCCGCATCTCTGTTCCATGTTTCTAATGAT
TACTCATTCATTCGATATTACACAAGGACCCTTTACCAATTCCAGTTTCAAGAAGCACT
TTGTCAAGCAGCTAAACATGAAGGCCCTCTGCACAAATGTGACATCTCAAACTCTACA
GAAGCTGGACAGAAACTGTTCAATATGCTGAGGCTTGGAAAATCAGAACCCTGGACC
CTAGCATTGGAAAATGTTGTAGGAGCAAAGAACATGAATGTAAGGCCACTGCTCAAC
TACTTTGAGCCCTTATTTACCTGGCTGAAAGACCAGAACAAGAATTCTTTTGTGGGAT
GGAGTACCGACTGGAGTCCATATGCAGACCAAAGCATCAAAGTGAGGATAAGCCTAA
AATCAGCTCTTGGAGATAGAGCATATGAATGGAACGACAATGAAATGTACCTGTTCCG
ATCATCTGTTGCATATGCTATGAGGCAGTACTTTTTAAAAGTAAAAAATCAGATGATTC
TTTTTGGGGAGGAGGATGTGCGAGTGGCTAATTTGAAACCAAGAATCTCCTTTAATTT
CTTTGTCACTGCACCTAAAAATGTGTCTGATATCATTCCTAGAACTGAAGTTGAAAAG
GCCATCAGGATGTCCCGGAGCCGTATCAATGATGCTTTCCGTCTGAATGACAACAGC
CTAGAGTTTCTGGGGATACAGCCAACACTTGGACCTCCTAACCAGCCCCCTGTTTCC
AGTGGTGGCGGAGGGACCAGCGACTACAAAGACGATGACGACAAGTAATAAAGGGC
GAATTCCAGCACACTGGCGGCCGTTACTAGTGGATCCGGCTGCTAACAAAGCCCGAAA

GGAAGCTGAGTTGGCTGCTGCCACCGCTGAGCAATAACTAGCATAACCCCTTGGGGCCTCTAAACGGGTCTTGAGGGGTTTTTTG

NC-HA
https://benchling.com/s/seq-cZNiRHDjuuyqQ7DkNkbF?m=slm-4PCq6MvuVgR9KZxSFPbH
This negative control, non-fluorescent protein is the tail tubular protein of the T4 phage, gp11, and is unrelated to SARS-CoV
TAATACGACTCACTATAGGGAGACCACAACGGTTTCCCTCTAGAAATAATTTTGTTTAACTTTAAGAAGGAGATATACC**ATGAGTTTACTTAATAATAAAGCGGGAGTTATTTCCCGCTTAGCCGATTTTCTTGGTTTTAGACCTAAAACTGGCGACATTGATGTAATGAATCGTCAATCAGTCGGGTCAGTGACAATATCTCAATTAGCGAAAGGATTTTATGAACCAAACATAGAATCAGCTATTAATGACGTTCATAATTTTTCTATAAAAGACGTTGGCACAATTATTACTAATAAAACTGGTGTTTCTCCTGAGGGTGTTTCTCAAACTGATTATTGGGCATTTTCTGGAACTGTAACAGACGATTCTCTTCCTCCGGGTTCTCCTATTACGGTATTAGTATTTGGTCTTCCAGTTTCAGCAACAACTGGAATGACGGCAATTGAGTTTGTTGCAAAAGTTCGCGTTGCACTACAAGAAGCTATTGCGTCATTTACTGCTATCAATTCATATAAAGACCATCCAACTGATGGTAGTAAATTAGAAGTTACTTATTTAGATAATCAAAAACATGTATTAAGCACATATTCTACATATGGAATAACTATTTCCCAAGAAATTATATCTGAGTCTAAGCCTGGCTATGGTACATGGAATTTATTGGGCGCACAAACTGTAACTTTAGATAATCAGCAGACTCCTACAGTATTTTATCATTTTGAGAGAACAGCA**ACCAGCTACCCATACGATGTTCCAGATTACGCTTAATAAAAGGGCGAATTCCAGCACACTGGCGGCCGTTACTAGTGGATCCGGCTGCTAACAAAGCCCGAAAGGAAGCTGAGTTGGCTGCTGCCACCGCTGAGCAATAACTAGCATAACCCCTTGGGGCCTCTAAACGGGTCTTGAGGGGTTTTTTG

S1-GFP-HA for human cell-extract expression
https://benchling.com/s/seq-8Xpvg1caAgfXqfUCJNLo?m=slm-NvXUS4NU52YIND2mBfZX
T7 promoter, IRES, GFP, **protein of interest**, HA tag, *poly(A)*, T7 terminator
GATGTGCTGCAAGGCGATTAAGTTGGGTAACGCCAGGGTTTTCCCAGTCACGACGTTGTAAAACGACGGCCAGTGAATTGTAATACGACTCACTATAGGGCGAATTAATTCCGGTTATTTTCCACCATATTGCCGTCTTTTGGCAATGTGAGGGCCCGGAAACCTGGCCCTGTCTTCTTGACGAGCATTCCTAGGGGTCTTTCCCCTCTCGCCAAAGGAATGCAAGGTCTGTTGAATGTCGTGAAGGAAGCAGTTCCTCTGGAAGCTTCTTGAAGACAAACAACGTCTGTAGCGACCCTTTGCAGGCAGCGGAACCCCCCACCTGGCGACAGGTGCCTCTGCGGCCAAAAGCCACGTGTATAAGATACACCTGCAAAGGCGGCACAACCCCAGTGCCACGTTGTGAGTTGGATAGTTGTGGAAAGAGTCAAATGGCTCACCTCAAGCGTATTCAACAAGGGGCTGAAGGATGCCCAGAAGGTACCCCATTGTATGGGATCTGATCTGGGGCCTCGGTGCACATGCTTTACATGTGTTTAGTCGAGGTTAAAAAACGTCTAGGCCCCCCGAACCACGGGGACGTGGTTTTCCTTTGAAAAACACGATGATAATG**AGTCAGTGTGTTAATCTTACAACCAGAACTCAATTACCCCCTGCATACACTAATTCTTTCACACGTGGTGTTTATTACCCTGACAAAGTTTTCAGATCCTCAGTTTTACATTCAACTCAGGACTTGTTCTTACCTTTCTTTTCCAATGTTACTTGGTTCCATGCTATACATGTCTCTGGGACCAATGGTACTAAGAGGTTTGATAACCC**

TGTCCTACCATTTAATGATGGTGTTTATTTTGCTTCCACTGAGAAGTCTAACATAATAA
GAGGCTGGATTTTTGGTACTACTTTAGATTCGAAGACCCAGTCCCTACTTATTGTTAAT
AACGCTACTAATGTTGTTATTAAAGTCTGTGAATTTCAATTTTGTAATGATCCATTTTTG
GGTGTTTATTACCACAAAAACAACAAAAGTTGGATGGAAAGTGAGTTCAGAGTTTATT
CTAGTGCGAATAATTGCACTTTTGAATATGTCTCTCAGCCTTTTCTTATGGACCTTGAA
GGAAAACAGGGTAATTTCAAAAATCTTAGGGAATTTGTGTTTAAGAATATTGATGGTTA
TTTTAAAATATATTCTAAGCACACGCCTATTAATTTAGTGCGTGATCTCCCTCAGGGTT
TTTCGGCTTTAGAACCATTGGTAGATTTGCCAATAGGTATTAACATCACTAGGTTTCAA
ACTTTACTTGCTTTACATAGAAGTTATTTGACTCCTGGTGATTCTTCTTCAGGTTGGAC
AGCTGGTGCTGCAGCTTATTATGTGGGTTATCTTCAACCTAGGACTTTTCTATTAAAAT
ATAATGAAAATGGAACCATTACAGATGCTGTAGACTGTGCACTTGACCCTCTCTCAGA
AACAAAGTGTACGTTGAAATCCTTCACTGTAGAAAAAGGAATCTATCAAACTTCTAACT
TTAGAGTCCAACCAACAGAATCTATTGTTAGATTTCCTAATATTACAAACTTGTGCCCT
TTTGGTGAAGTTTTTAACGCCACCAGATTTGCATCTGTTTATGCTTGGAACAGGAAGA
GAATCAGCAACTGTGTTGCTGATTATTCTGTCCTATATAATTCCGCATCATTTTCCACT
TTTAAGTGTTATGGAGTGTCTCCTACTAAATTAAATGATCTCTGCTTTACTAATGTCTAT
GCAGATTCATTTGTAATTAGAGGTGATGAAGTCAGACAAATCGCTCCAGGGCAAACT
GGAAAGATTGCTGATTATAATTATAAATTACCAGATGATTTTACAGGCTGCGTTATAGC
TTGGAATTCTAACAATCTTGATTCTAAGGTTGGTGGTAATTATAATTACCTGTATAGATT
GTTTAGGAAGTCTAATCTCAAACCTTTTGAGAGAGATATTTCAACTGAAATCTATCAGG
CCGGTAGCACACCTTGTAATGGTGTTGAAGGTTTTAATTGTTACTTTCCTTTACAATCA
TATGGTTTCCAACCCACTAATGGTGTTGGTTACCAACCATACAGAGTAGTAGTACTTT
CTTTTGAACTTCTACATGCACCAGCAACTGTTTGTGGACCTAAAAAGTCTACTAATTTG
GTTAAAAACAAATGTGTCAATTTCAACTTCAATGGTTTAACAGGCACAGGTGTTCTTAC
TGAGTCTAACAAAAAGTTTCTGCCTTTCCAACAATTTGGCAGAGACATTGCTGACACT
ACTGATGCTGTCCGTGATCCACAGACACTTGAGATTCTTGACATTACACCATGTTCTT
TTGGTGGTGTCAGTGTTATAACACCAGGAACAAATACTTCTAACCAGGTTGCTGTTCT
TTATCAGGATGTTAACTGCACAGAAGTCCCTGTTGCTATTCATGCAGATCAACTTACT
CCTACTTGGCGTGTTTATTCTACAGGTTCTAATGTTTTTCAAACACGTGCAGGCTGTTT
AATAGGGGCTGAACATGTCAACAACTCATATGAGTGTGACATACCCATTGGTGCAGG
TATATGCGCTAGTTATCAGACTCAGACTAATTCTCCTCGGCGGGCACGTAAGCGAGC
CCCAGGAACGTCTATGGTAAGCAAAGGGGAGGAACTCTTCACAGGTGTAGTGCCGATT
TTGGTCGAACTTGACGGAGATGTCAATGGACATAAGTTCTCTGTGAGTGGTGAAGGTGA
GGGCGATGCCACTTACGGTAAGCTTACTCTGAAGTTCATATGCACAACTGGTAAGTTGC
CCGTACCTTGGCCTACGCTCGTCACGACACTCACCTACGGCGTCCAGTGTTTCTCCCG
CTACCCCGATCATATGAAGCAGCACGATTTCTTCAAGTCTGCGATGCCAGAAGGGTATG
TTCAGGAGAGGACTATCTTCTTTAAGGACGACGGGAATTACAAAACCAGAGCCGAAGTG
AAGTTTGAGGGTGATACCCTCGTTAACAGGATAGAACTGAAGGGGATTGATTTTAAGGA
GGATGGAAACATATTGGGACATAAATTGGAATATAATTACAACTCTCATAATGTTTACATT
ATGGCTGACAAACAAAAAAACGGCATAAAGGTCAACTTTAAGATTCGACATAACATCGAA
GACGGGAGCGTTCAACTGGCTGATCATTATCAACAGAACACTCCAATTGGCGACGGTC
CGGTATTGCTTCCTGATAACCATTACCTGTCAACCCAGTCTGCGCTGTCCAAGGATCCT
AATGAGAAAAGGGATCATATGGTGCTGTTGGAATTTGTTACAGCCGCAGGGATAACTCT
GGGGATGGACGAACTGTATAAGCGCCGCGCAAAGAAGGAGGCGTACCCATACGACGT

ACCGGACTACGCTTGAGATCTGACTG*AAAAAAAAAAAAAAAAAAAAAAAAAAAAAA*GTTTAA
ACACTAGTCCGCTGAGCAATAACTAGCATAACCCCTTGGGGCCTCTAAACGGGTCTTGA
GGGGTTTTTTGCTGAAAGGAGGAACTATATCCGGGCTTCCTCGCTCACTGACTCGCTGC
GCTCGGTCGTTCGGCTGCGGCGAGCGGTATCAGCTCACTCAAAGG

GFP-S1-HA for human cell-extract expression
https://benchling.com/s/seq-rwfpVokSGv295mnwNUYI?m=slm-0Gn5sNKvw4nEF3zR7XWG
GATGTGCTGCAAGGCGATTAAGTTGGGTAACGCCAGGGTTTTCCCAGTCACGACGTTGT
AAAACGACGGCCAGTGAATTGTAATACGACTCACTATAGGGCGAATTAATTCCGGTTATT
TTCCACCATATTGCCGTCTTTTGGCAATGTGAGGGCCCGGAAACCTGGCCCTGTCTTCT
TGACGAGCATTCCTAGGGGTCTTTCCCCTCTCGCCAAAGGAATGCAAGGTCTGTTGAAT
GTCGTGAAGGAAGCAGTTCCTCTGGAAGCTTCTTGAAGACAAACAACGTCTGTAGCGAC
CCTTTGCAGGCAGCGGAACCCCCCACCTGGCGACAGGTGCCTCTGCGGCCAAAAGCC
ACGTGTATAAGATACACCTGCAAAGGCGGCACAACCCCAGTGCCACGTTGTGAGTTGG
ATAGTTGTGGAAAGAGTCAAATGGCTCACCTCAAGCGTATTCAACAAGGGGCTGAAGGA
TGCCCAGAAGGTACCCCATTGTATGGGATCTGATCTGGGGCCTCGGTGCACATGCTTTA
CATGTGTTTAGTCGAGGTTAAAAAACGTCTAGGCCCCCCGAACCACGGGGACGTGGTT
TTCCTTTGAAAAACACGATGATAAATGGTAAGCAAAGGGGAGGAACTCTTCACAGGTGT
AGTGCCGATTTTGGTCGAACTTGACGGAGATGTCAATGGACATAAGTTCTCTGTGAGTG
GTGAAGGTGAGGGCGATGCCACTTACGGTAAGCTTACTCTGAAGTTCATATGCACAACT
GGTAAGTTGCCCGTACCTTGGCCTACGCTCGTCACGACACTCACCTACGGCGTCCAGT
GTTTCTCCCGCTACCCCGATCATATGAAGCAGCACGATTTCTTCAAGTCTGCGATGCCA
GAAGGGTATGTTCAGGAGAGGACTATCTTCTTTAAGGACGACGGGAATTACAAAACCAG
AGCCGAAGTGAAGTTTGAGGGTGATACCCTCGTTAACAGGATAGAACTGAAGGGGATT
GATTTTAAGGAGGATGGAAACATATTGGGACATAAATTGGAATATAATTACAACTCTCATA
ATGTTTACATTATGGCTGACAAACAAAAAAACGGCATAAAGGTCAACTTTAAGATTCGAC
ATAACATCGAAGACGGGAGCGTTCAACTGGCTGATCATTATCAACAGAACACTCCAATT
GGCGACGGTCCGGTATTGCTTCCTGATAACCATTACCTGTCAACCCAGTCTGCGCTGTC
CAAGGATCCTAATGAGAAAAGGGATCATATGGTGCTGTTGGAATTTGTTACAGCCGCAG
GGATAACTCTGGGGATGGACGAACTGTATAAGATG**AGTCAGTGTGTTAATCTTACAACC**
**AGAACTCAATTACCCCCTGCATACACTAATTCTTTCACACGTGGTGTTTATTACCCTGA**
**CAAAGTTTTCAGATCCTCAGTTTTACATTCAACTCAGGACTTGTTCTTACCTTTCTTTTC**
**CAATGTTACTTGGTTCCATGCTATACATGTCTCTGGGACCAATGGTACTAAGAGGTTT**
**GATAACCCTGTCCTACCATTTAATGATGGTGTTTATTTTGCTTCCACTGAGAAGTCTAA**
**CATAATAAGAGGCTGGATTTTTGGTACTACTTTAGATTCGAAGACCCAGTCCCTACTT**
**ATTGTTAATAACGCTACTAATGTTGTTATTAAAGTCTGTGAATTTCAATTTTGTAATGAT**
**CCATTTTTGGGTGTTTATTACCACAAAAACAACAAAAGTTGGATGGAAAGTGAGTTCA**
**GAGTTTATTCTAGTGCGAATAATTGCACTTTTGAATATGTCTCTCAGCCTTTTCTTATG**
**GACCTTGAAGGAAAACAGGGTAATTTCAAAAATCTTAGGGAATTTGTGTTTAAGAATAT**
**TGATGGTTATTTTAAAATATATTCTAAGCACACGCCTATTAATTTAGTGCGTGATCTCC**
**CTCAGGGTTTTTCGGCTTTAGAACCATTGGTAGATTTGCCAATAGGTATTAACATCACT**
**AGGTTTCAAACTTTACTTGCTTTACATAGAAGTTATTTGACTCCTGGTGATTCTTCTTCA**
**GGTTGGACAGCTGGTGCTGCAGCTTATTATGTGGGTTATCTTCAACCTAGGACTTTTC**

TATTAAAATATAATGAAAATGGAACCATTACAGATGCTGTAGACTGTGCACTTGACCCT
CTCTCAGAAACAAAGTGTACGTTGAAATCCTTCACTGTAGAAAAAGGAATCTATCAAA
CTTCTAACTTTAGAGTCCAACCAACAGAATCTATTGTTAGATTTCCTAATATTACAAACT
TGTGCCCTTTTGGTGAAGTTTTTAACGCCACCAGATTTGCATCTGTTTATGCTTGGAAC
AGGAAGAGAATCAGCAACTGTGTTGCTGATTATTCTGTCCTATATAATTCCGCATCATT
TTCCACTTTTAAGTGTTATGGAGTGTCTCCTACTAAATTAAATGATCTCTGCTTTACTAA
TGTCTATGCAGATTCATTTGTAATTAGAGGTGATGAAGTCAGACAAATCGCTCCAGGG
CAAACTGGAAAGATTGCTGATTATAATTATAAATTACCAGATGATTTTACAGGCTGCGT
TATAGCTTGGAATTCTAACAATCTTGATTCTAAGGTTGGTGGTAATTATAATTACCTGTA
TAGATTGTTTAGGAAGTCTAATCTCAAACCTTTTGAGAGAGATATTTCAACTGAAATCT
ATCAGGCCGGTAGCACACCTTGTAATGGTGTTGAAGGTTTTAATTGTTACTTTCCTTTA
CAATCATATGGTTTCCAACCCACTAATGGTGTTGGTTACCAACCATACAGAGTAGTAG
TACTTTCTTTTGAACTTCTACATGCACCAGCAACTGTTTGTGGACCTAAAAAGTCTACT
AATTTGGTTAAAAACAAATGTGTCAATTTCAACTTCAATGGTTTAACAGGCACAGGTGT
TCTTACTGAGTCTAACAAAAAGTTTCTGCCTTTCCAACAATTTGGCAGAGACATTGCT
GACACTACTGATGCTGTCCGTGATCCACAGACACTTGAGATTCTTGACATTACACCAT
GTTCTTTTGGTGGTGTCAGTGTTATAACACCAGGAACAAATACTTCTAACCAGGTTGC
TGTTCTTTATCAGGATGTTAACTGCACAGAAGTCCCTGTTGCTATTCATGCAGATCAA
CTTACTCCTACTTGGCGTGTTTATTCTACAGGTTCTAATGTTTTTCAAACACGTGCAGG
CTGTTTAATAGGGGCTGAACATGTCAACAACTCATATGAGTGTGACATACCCATTGGT
GCAGGTATATGCGCTAGTTATCAGACTCAGACTAATTCTCCTCGGCGGGCACGTAAG
CGAGCCCCAGGAACGTCTCGCCGCGCAAAGAAGGAGGCGTACCCATACGACGTACCG
GACTACGCTTGAGATCTGACTG*AAAAAAAAAAAAAAAAAAAAAAAAAAAA*GTTTAAACACT
AGTCCGCTGAGCAATAACTAGCATAACCCCTTGGGGCCTCTAAACGGGTCTTGAGGGG
TTTTTTGCTGAAAGGAGGAACTATATCCGGGCTTCCTCGCTCACTGACTCGCTGCGCTC
GGTCGTTCGGCTGCGGCGAGCGGTATCAGCTCACTCAAAGG

Supplementary Table 2 | Experimental conditions. Attached separately.

Supplementary Table 3 | Specificity and sensitivity of IgG detection in human sera.

| Antigen | N | Na | Nb | Nc | Nd | Ne | Nf | Ng | Nh | S1-G | S1a | S1b |
|---|---|---|---|---|---|---|---|---|---|---|---|---|
| Sensitivity (%) | 70 | 85 | 65 | 50 | 85 | 90 | 35 | 80 | 75 | 50 | 20 | 35 |
| Specificity (%) | 67 | 50 | 67 | 83 | 33 | 67 | 83 | 100 | 100 | 67 | 100 | 83 |

| Antigen | S1c | S1d | S1e | S1f | S1g | S1h | S1i | S1j | S1k | S1l | G-S1 |
|---|---|---|---|---|---|---|---|---|---|---|---|
| Sensitivity (%) | 5 | 5 | 20 | 10 | 20 | 20 | 25 | 25 | 35 | 15 | 30 |
| Specificity (%) | 100 | 100 | 100 | 100 | 100 | 100 | 100 | 100 | 100 | 100 | 100 |

| Antigen | R2-G | G-R2 | G-d | G-o | R1-G | G-R1 |
|---|---|---|---|---|---|---|
| Sensitivity (%) | 30 | 10 | 15 | 10 | 35 | 15 |
| Specificity (%) | 100 | 100 | 100 | 100 | 100 | 100 |

### References


1. Buxboim, A., Daube, S. S. & Bar-Ziv, R. Synthetic gene brushes: a structure–function relationship. Mol Syst Biol 4, 181 (2008).
2. Sprague, B. L. et al. Analysis of binding at a single spatially localized cluster of binding sites by fluorescence recovery after photobleaching. Biophys J 91, 1169–1191 (2006).
3. Cayley, S., Lewis, B. A., Guttman, H. J. & Record, M. T. Characterization of the cytoplasm of Escherichia coli K-12 as a function of external osmolarity: Implications for protein-DNA interactions in vivo. J Mol Biol 222, 281–300 (1991).
4. Bracha, D., Karzbrun, E., Shemer, G., Pincus, P. A. & Bar-Ziv, R. H. Entropy-driven collective interactions in DNA brushes on a biochip. Proceedings of the National Academy of Sciences 110, 4534–4538 (2013).
5. Karzbrun, E., Tayar, A. M., Noireaux, V. & Bar-Ziv, R. H. Programmable on-chip DNA compartments as artificial cells. Science (1979) 345, 829–832 (2014).
6. Landry, J. P., Ke, Y., Yu, G.-L. & Zhu, X. D. Measuring affinity constants of 1450 monoclonal antibodies to peptide targets with a microarray-based label-free assay platform. J Immunol Methods 417, 86–96 (2015).
7. Barton, M. I. et al. Effects of common mutations in the SARS-CoV-2 Spike RBD and its ligand, the human ACE2 receptor on binding affinity and kinetics. Elife 10, e70658 (2021).
8. Abcam. Anti-SARS-CoV-2 Spike RBD antibody [CV30].
9. Hurlburt, N. K. et al. Structural basis for potent neutralization of SARS- CoV-2 and role of antibody affinity maturation. Nat Commun (2020) doi:10.1038/s41467-020-19231-9.
